# H3.3-G34R Mutation-Mediated Epigenetic Reprogramming Leads to Enhanced Efficacy of Immune Stimulatory Gene Therapy in Diffuse Hemispheric Gliomas

**DOI:** 10.1101/2023.06.13.544658

**Authors:** Maria B. Garcia-Fabiani, Santiago Haase, Kaushik Banerjee, Ziwen Zhu, Brandon L. McClellan, Anzar A. Mujeeb, Yingxiang Li, Claire E. Tronrud, Maria L. Varela, Molly E.J. West, Jin Yu, Padma Kadiyala, Ayman W. Taher, Felipe J. Núñez, Mahmoud S. Alghamri, Andrea Comba, Flor M. Mendez, Alejandro J. Nicola Candia, Brittany Salazar, Fernando M. Nunez, Marta B. Edwards, Tingting Qin, Rodrigo T. Cartaxo, Michael Niculcea, Carl Koschmann, Sriram Venneti, Montserrat Puigdelloses Vallcorba, Emon Nasajpour, Giulia Pericoli, Maria Vinci, Claudia L. Kleinman, Nada Jabado, James P. Chandler, Adam M. Sonabend, Michael DeCuypere, Dolores Hambardzumyan, Laura M. Prolo, Kelly B. Mahaney, Gerald A. Grant, Claudia K Petritsch, Joshua D. Welch, Maureen A. Sartor, Pedro R. Lowenstein, Maria G. Castro

## Abstract

Diffuse hemispheric glioma (DHG), H3 G34-mutant, representing 9-15% of cases, are aggressive Central Nervous System (CNS) tumors with poor prognosis. This study examines the role of epigenetic reprogramming of the immune microenvironment and the response to immune-mediated therapies in G34-mutant DHG. To this end, we utilized human G34-mutant DHG biopsies, primary G34-mutant DHG cultures, and genetically engineered G34-mutant mouse models (GEMMs). Our findings show that the G34 mutation alters histone marks’ deposition at promoter and enhancer regions, leading to the activation of the JAK/STAT pathway, which in turn results in an immune-permissive tumor microenvironment. The implementation of Ad-TK/Ad-Flt3L immunostimulatory gene therapy significantly improved median survival, and lead to over 50% long term survivors. Upon tumor rechallenge in the contralateral hemisphere without any additional treatment, the long-term survivors exhibited robust anti-tumor immunity and immunological memory. These results indicate that immune-mediated therapies hold significant potential for clinical translation in treating patients harboring H3.3-G34 mutant DHGs, offering a promising strategy for improving outcomes in this challenging cancer subtype affecting adolescents and young adults (AYA).

**STATEMENT OF SIGNIFICANCE:** This study uncovers the role of the H3.3-G34 mutation in reprogramming the tumor immune microenvironment in diffuse hemispheric gliomas. Our findings support the implementation of precision medicine informed immunotherapies, aiming at improving enhanced therapeutic outcomes in adolescents and young adults harboring H3.3-G34 mutant DHGs.

## INTRODUCTION

Central nervous system (CNS) tumors are the leading cause of cancer related deaths in people ranging from 0 to 14 years in the United States (1). These tumors have an incidence of 5.74 per 100,000 population, and a mortality rate of 4.42 per 100,000 population for malignant tumors(1). Pediatric-type diffuse high-grade gliomas (PDHGGs) are diffuse, infiltrating, and highly aggressive tumors(2–4). Currently, they remain incurable, with a 5-year overall survival of less than 20%(2). Despite these adverse statistics, the last decade has been crucial in the understanding of the molecular features of PDHGGs (5–7). One of the breakthroughs in pediatric glioma biology was the discovery of mutations in the histone gene H3. These mutations are involved in glioma development, and their presence correlates with distinct genomic alterations, epigenetic landscapes, CNS locations and clinical features(5–10).

In this study, we focused on the subtype of PDHGGs harboring a mutation in the histone H3.3 gene *H3F3A*, that changes the glycine at the position 34 to an arginine (H3.3-G34R) or, less commonly, to a valine(5,7). This mutation is present in 9-15% of PDHGGs, is restricted to the cerebral hemispheres, and is found predominantly in the adolescent population (median 15.0 years)(5–7,11,12). G34-mutant Diffuse hemispheric gliomas (DHGs) are characterized by the presence of loss of function mutations in *ATRX/DAXX* and *TP53* genes, and a DNA hypomethylated phenotype(5–7). The prognosis of these tumors is dismal, with patients displaying a median survival of 18.0 months, and a 2-year overall survival of 27.3%(5). The mechanisms that underlie H3.3-G34R/V subgroup’s disease progression in DHGs are not well understood(6,13–16). Presently, the standard of care (SOC) for DHGs, irrespective of their H3 status, includes maximum surgical removal followed by targeted radiation. The benefit of chemotherapy in addition to radiation for G34-mutant DHGs is still under debate, largely because earlier cooperative group studies did not consistently test and stratify for G34 mutations(17).

The implementation of effective immunotherapies for DHG remains underdeveloped. Recent studies in DHG have shown a complex and dynamic interaction between the tumor cells and the immune system(18). DHGs are heterogeneous tumors with a significant portion of the tumor mass consisting of non-malignant cells, such as macrophages, microglia, and dendritic cells, as well as myeloid-derived suppressor cells (MDSCs), which are found extensively in the DHG tumor microenvironment (TME) and have been identified as major contributors to immunosuppression within the TME(19,20). Thus, the development of immunotherapies has been limited by several factors, notably the immunosuppressive TME frequently found in these tumors, which is influenced by many factors, including the specific tumor subtype, the genetic aberrations found within the tumor, and the host immune system.

Understanding the molecular characteristics of the DHG tumor cells and their interactions with the TME is critical to evaluate the implementation of immunotherapies for G34-mutant DHG. In this study, we aimed to understand the biological and molecular features of DHGs harboring H3.3-G34R mutations, as well as its immunological landscape. So far, the tumor immune microenvironment (TIME) of brain tumors in adolescents and young adults has not been understudied, holding back the successful development of targeted immunotherapies(21–23). We generated genetically engineered mouse models (GEMM) of DHGs harboring the H3.3-G34R mutation, in combination with *ATRX* and *TP53* downregulation, which recapitulates salient features of the human disease(17,24). A previous study from our lab using this model allowed us to identify that H3.3-G34-mutations drive DNA repair impairment and genetic instability in DHG, leading to the activation of the cGAS-STING pathway and enhanced anti-tumoral immunological responses(17). Thus, in this study, we aimed at characterizing the consequences of the G34-mutations in shaping the TIME.

Using ChIP-Seq and RNA-Seq, we uncovered the molecular landscape of H3.3-G34R DHG cells, unveiling the consequences of the H3.3-G34R mutation at the epigenetic and transcriptional levels. Our results revealed that the presence of H3.3-G34R correlates with the epigenetic and transcriptional upregulation of pathways related to immune-activation. We showed that this correlates with the posttranslational activation of the JAK-STAT signaling pathway in G34-mutant cells. Then we studied the immune cell composition of the H3.3-G34R DHG TME and discovered that H3.3-G34R DHG have a less immunosuppressive TME with respect to the control group, harboring the wild-type H3.3. Leveraging these differences, we tested the efficacy of Ad-TK/Ad-Flt3L immune stimulatory gene therapy(19,20,25–28). We observed an effective anti-tumor CD8 T cell immune response, an increase in the median survival of H3.3-G34R-DHG bearing animals, and the onset of anti-tumor immunological memory. Collectively, our results uncovered molecular features of H3.3-G34R DHGs and suggest that immune-stimulatory therapeutic approaches, such as the TK/Flt3L gene therapy, could be considered as an attractive adjuvant treatment for H3.3-G34R DHGs.

## RESULTS

### Development of a genetically engineered mouse model for H3.3-G34R DHGs

Using the Sleeping Beauty (SB) transposon system, we generated an immunocompetent mouse model for DHGs harboring H3.3-G34R mutation (H3.3-G34R group) or H3.3-wild type (WT) (control group)(17). This system allows the *de novo* induction of brain tumors using non-viral, transposon-mediated integration of plasmid DNA into cells of the sub-ventricular zone of neonatal mice by the SB-transposase (17,24,29–32) (Figures S1, A-B). The H3.3-G34R DHG were modelled using DNA plasmids encoding: i) *NRASG12V* (NRAS activating mutation) (the receptor tyrosine kinase (RTK)–RAS–phosphatidylinositol 3-kinase (PI3K) pathway is activated in a large number of DHGs (5,7,33–36)); short hairpin (sh) RNAs targeting ii) *Atrx* and iii) *Trp53* genes; with or without the mutated histone H3.3-G34R (H3.3-G34R group)(Figure S1, A)(17). Mice injected with these plasmids developed tumors and those expressing the H3.3-G34R displayed longer median survival (MS) with respect to the H3.3-WT group (H3.3-G34R, MS=161 days; H3.3-WT, MS=88 days) (Figure S1, A). We were able to verify the development of tumor as the expression of each genetic lesion is coupled to the expression of fluorescent proteins (Figure S1, B). H3.3-G34R expression in the genetically engineered DHG model was confirmed by immunohistochemistry (IHC) (Figure S2, A-B). These tumors were diffusely infiltrative and hypercellular (Figure S2, D-E). Also, they showed other salient characteristics of DHG such as pseudopalisades with necrotic areas, microvascular proliferation, hemorrhagic areas, multinucleated giant cells and nuclear atypia(13,37) (Figure S2, D-E). We confirmed by western blotting that primary cells generated from tumors expressed H3.3-G34R (Figure S2, C). Additionally, we observed that the oligodendrocyte marker OLIG2, a hallmark of these tumors, was downmodulated in the H3.3-G34R DHG cells (Figure S2, C) (5,6,37).

To better understand the effects of H3.3-G34R expression on the transcriptional landscape of glioma cells, we performed RNA-Seq on H3.3-G34R and H3.3-WT cells extracted from the TME of SB-derived DHG. Tumor cells were separated from the immune cells (CD45+) and the RNA-Seq analysis was conducted on tumor-enriched cell fractions(38)(Figure S1, C). Among the 22,828 genes that were measured, 353 were found to be differentially expressed between H3.3-G34R vs H3.3-WT tumor cells (Fold Change: 0.585, adjusted p value <0.05) (<u>Table S1</u>) (Figure 1, A). Amongst the upregulated Gene Ontologies (GOs) in the H3.3-G34R group, we observed terms related to immune-mediated processes, such as “Innate immune response” and “Regulation of adaptive immune response” (Figure 1, B). Particularly, we found GOs related to interferon I (IFN-I) (IFNα and IFNβ) and interferon II (IFN-II) (IFNγ) signaling, and several genes involved in these pathways were upregulated in H3.3-G34R tumor cells versus H3.3-WT tumor cells (Figures S1, D and S3, B). The upregulation of type I IFN pathways aligns with previous work from our lab showing that DNA damage deficiencies and genetic instability in G34-mutant DHG lead to activation of the cGAS-STING pathway in G34-mutant DHG(17) (Figure S1, E). Among the downregulated GOs in the H3.3-G34R group, we found several GOs related to CNS development (Figure S3, A).

**Figure 1.**
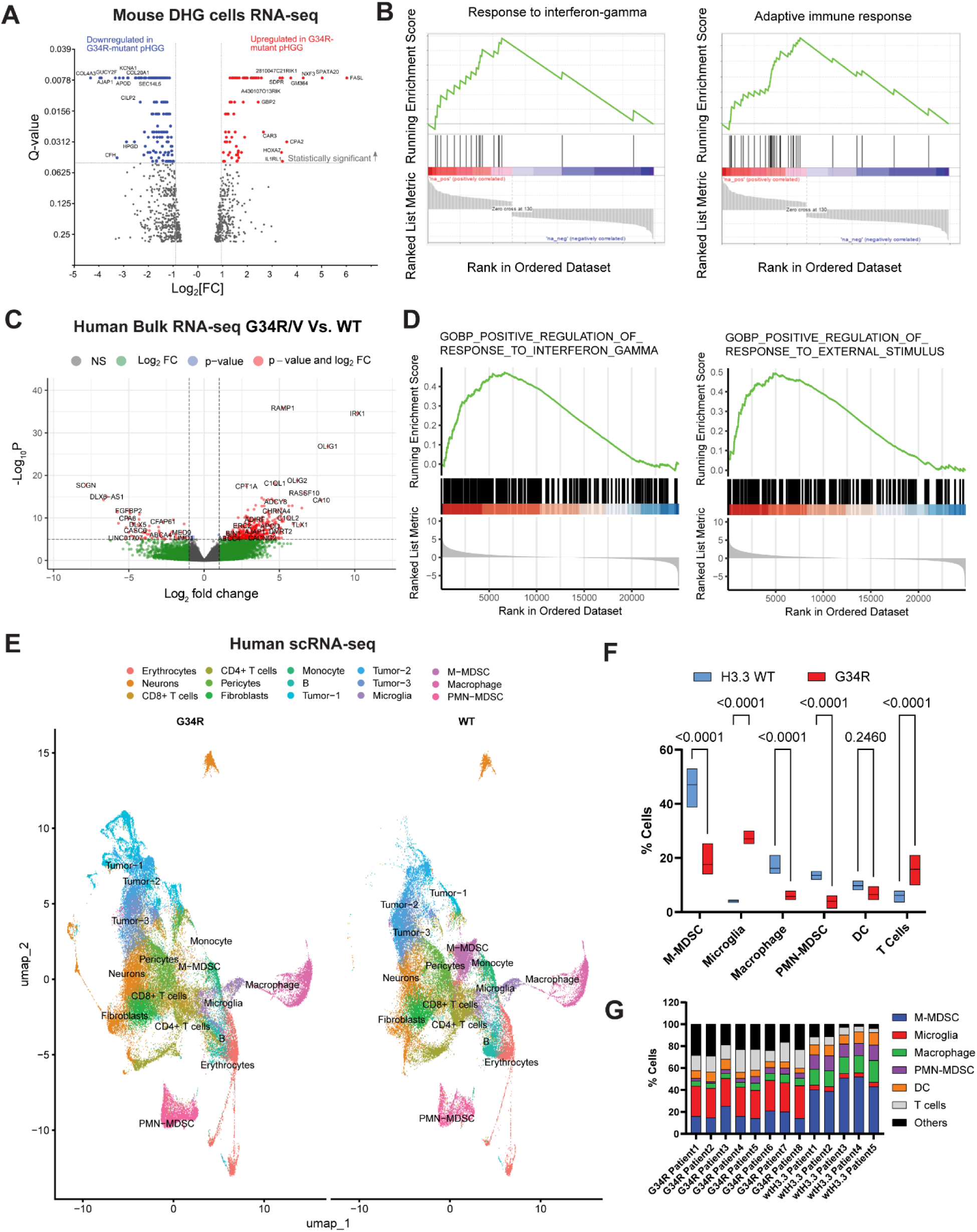
Transcriptomics analysis of mouse and human DHG. **(A)** Volcano plot depicting the transcriptional differences between mouse H3.3-G34R and H3.3-WT DHG tumor cells. **(B)** Gene set enrichment analysis (GSEA) showing the upregulation of interferon-gamma and adaptive immune pathways in mouse H3.3-G34R versus H3.3-WT DHG tumor cells. **(C)** Volcano plot depicting the transcriptional differences between H3.3 G34R/V tumors compared to H3.3 WT HGG tumors. **(D)** Gene set enrichment analysis (GSEA) showing the upregulation of interferon-gamma and immune pathways in H3.3 G34R/V versus H3.3 WT DHG tumors. **(E)** UMAP embedding of H3.3 G34R/V and WT DHG scRNA-seq. **(F)** Immune cell percentages from H3.3 G34R/V and WT samples. **(G)** Immune cell percentages from each individual patient sample.

### Characterization of the Tumor Immune Microenvironment (TIME) in human G34-Mutant DHGs

To investigate the impact of G34 mutations on the immune landscape of DHG, we performed a comprehensive analysis using both single-cell RNA sequencing (scRNA-seq) and bulk RNA sequencing data from human patient samples. First, we analyzed bulk RNA-seq data from human DHG samples, differential expression analysis revealed significant upregulation of immune-related gene ontologies in G34-mutant DHG compared to H3.3-WT DHG samples. The volcano plot (Figure 1, C) highlights these transcriptional differences. Gene Set Enrichment Analysis (GSEA) indicated a pronounced upregulation of interferon-gamma and other immune pathways in G34R/V samples (Figure 1, D). Notably, GOs related to STAT pathway activation were significantly upregulated in G34-mutant patient samples, corroborating our observations from mouse models. To further validate the observed trends, we conducted an analysis using independent bulk RNA-seq datasets from Chen *et al.*(*15*) and Mackay *et al.* (5), the Gene Set Variation Analysis (GSVA) scores consistently showed elevated immune pathway activity in G34-mutant DHGs compared to WT samples across both studies (Figure S4, A-B). To substantiate our findings, our scRNA-seq data from patient-derived samples revealed that G34-mutant DHGs exhibit a notably less immunosuppressive TME compared to histone WT patient samples. UMAP embedding analysis (Figure 1, E) demonstrated distinct clustering patterns between H3.3-G34R/V and H3.3-WT WT DHG samples. Analysis of immune cell percentages (Figure 1, F) revealed a significant decrease in monocytic myeloid-derived suppressor cells (M-MDSCs) and a slight decline in polymorphonuclear myeloid-derived suppressor cells (PMN-MDSCs). Additionally, there was a decrease in macrophages, accompanied by an increase in microglia, in H3.3-G34R/V samples. This trend was consistently observed across individual patient samples (Figure 1, G).

### The expression of the H3.3-G34R mutation leads to reprogramming of the molecular landscape of DHG cells, upregulating the expression of immune-related genes

The H3.3-G34 mutations occur close to a lysine located at the position 36 (H3K36) of the H3.3 histone tail. H3K36 can be trimethylated (H3K36me3) and this is regarded as a transcriptional activation mark (7,39). It has been demonstrated that, even though the H3.3-G34 mutations do not alter the global amounts of H3K36me3, they reduce the H3K36me2 and H3K36me3 deposition in *cis*, and affects the deposition of this epigenetic mark throughout the genome, shaping the transcriptional signature of the cell (5,8,40). Thus, to test the epigenetic consequences of the H3.3-G34R mutation in relation to histone mark deposition, we analyzed the differential binding of several histone marks throughout the H3.3-G34R and H3.3-WT DHG cells’ chromatin (Figure 2, A). Using ChIP-Seq, we assessed the binding of H3K4me1, H3K4me3, H3K27ac, H3K36me3 (transcriptional activation marks), and H3K27me3 (transcriptional repressive mark)(41), across the genome of three primary H3.3-WT and H3.3-G34R DHG cells (Figure 2). After the reads’ alignment and the fulfillment of quality controls, peak calling was performed to identify gene regions differentially enriched in specific histone marks, between H3.3-G34R and H3.3-WT DHG cells. Although no differences in global histone mark levels were found between H3.3-G34R and H3.3-WT DHG cells by Western blot (Figure S5, A), GSEA of differentially enriched genes in these marks allowed the identification of GO related to pathways or biological processes being epigenetically affected by H3.3-G34R expression (Figures 2, B-C, S5, B-D <u>and Table S2</u>). When exploring GOs related to the immune system activation/regulation, we observed differentially enriched immune-related GOs associated to H3K27me3 and H3K36me3 deposition (Figure 2, B-C). The H3K27me3 mark, which is associated to transcriptional repression, was enriched in GOs related to the regulation of signal transducer and activator of transcription (STAT) cascade and the activation of T cell-mediated, NK cell-mediated, B cell-mediated and humoral responses in H3.3-WT cells. On the other hand, in H3.3-G34R DHG cells, the H3K27me3 mark was enriched in GOs related to leucocyte apoptotic processes (Figure 2, B). Moreover, the H3K36me3 mark, which is associated to transcriptional activation, was enriched in GOs related to the negative regulation of T cell proliferation and activation, and the development of the immune system in H3.3-WT DHG cells; whereas this mark was found to be enriched in GOs related to leucocyte chemotaxis, myeloid cytokine production and STAT pathway activation in H3.3-G34R DHG cells (Figure 2, C). H3K27ac, H3K4me3 and H3K4me1 (Figure S5, B-D) were differentially bound to genes encountered in immune-related GOs. These results indicate that H3.3-G34R affects the deposition of different histone marks across the genome, specifically altering the epigenetic regulation of genes related to an active immune response.

**Figure 2.**
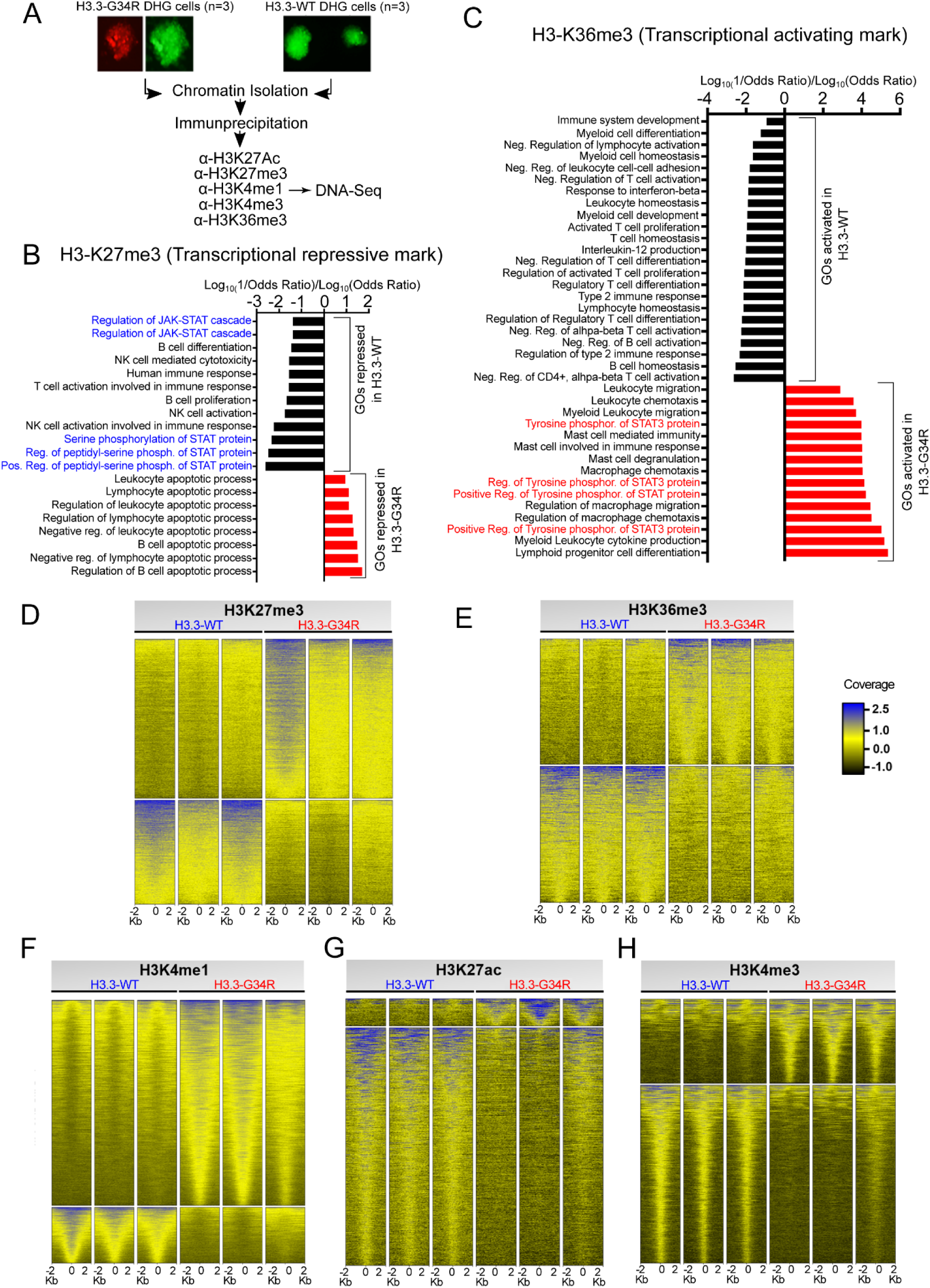
Characterization of the G34R DHG epigenome by ChIP-seq. **(A)** Experimental outline for the ChIP-Sequencing analysis. Three biological replicates of H3.3-WT and H3.3-G34R neurospheres (NS) were used to perform the ChIP-Seq. Five marks were analyzed: H3K27ac, H3K27me3, H3K4me1, H3K4me3 and H3K36me3. **(B-C)** Analysis of differentially enriched peaks for H3K27me3 **(B)** (transcriptional repression) or H3K36me3 **(C)** (transcriptional activation) histone marks revealed genes that are linked to distinct functional GO terms. GO enriched in the marks in H3.3-G34R versus H3.3-WT DHG are indicated with red bars and GO enriched in the marks in H3.3-WT versus H3.3-G34R DHG are indicated with black bars. The JAK/STAT related GO epigenetically active in H3.3-G34R cells (as indicated by enrichment of K36me3) are highlighted in red, and the same GO epigenetically downregulated in H3.3-WT cells are highlighted in blue. **(D-H)** Heat maps showing H3K27me3, H3K36me3, H3K4me1, H3K27ac and H3K4me3 differential peaks ± 2 kilo–base (Kb) pair from the gene center. Each row represents a distinct gene. The blue-to-yellow color gradient indicates high-to-low counts in the corresponding regions.

Since we found striking differences in histone mark deposition due to the expression of H3.3-G34R (Figure S6, A), we performed an analysis to define chromatin states (CS) at different gene regulatory regions (Figure S4, B-E). CS were determined by the combination and the relative abundance of different histone marks at a particular locus. Given that several genes related to IFN-I and IFN-II related pathways were upregulated in H3.3-G34R with respect to H3.3-WT DHG cells in the RNA-Seq analysis (Figure 1, A-B), we analyzed the epigenetic regulation and the CS changes of genes related to IFNγ signaling and STAT1 pathway activation. IFNγ is a cytokine secreted by activated CD4 and CD8 T cells, as well as by antigen presenting cells, and its pleiotropic function resides on its ability to modulate the expression of numerous genes (42–44). IFNγ canonical signaling pathway involves IFNγ binding to IFNGR1 and IFNGR2, the activation of JAK1 and JAK2, and the phosphorylation of STAT1 (pSTAT1) (42,44,45). The homodimerization of pSTAT1 forms a complex, which translocates to the nucleus and binds to gamma-activated sites (GAS), increasing the transcription of target genes (Figures 3, A and S7, A)(44).

**Figure 3.**
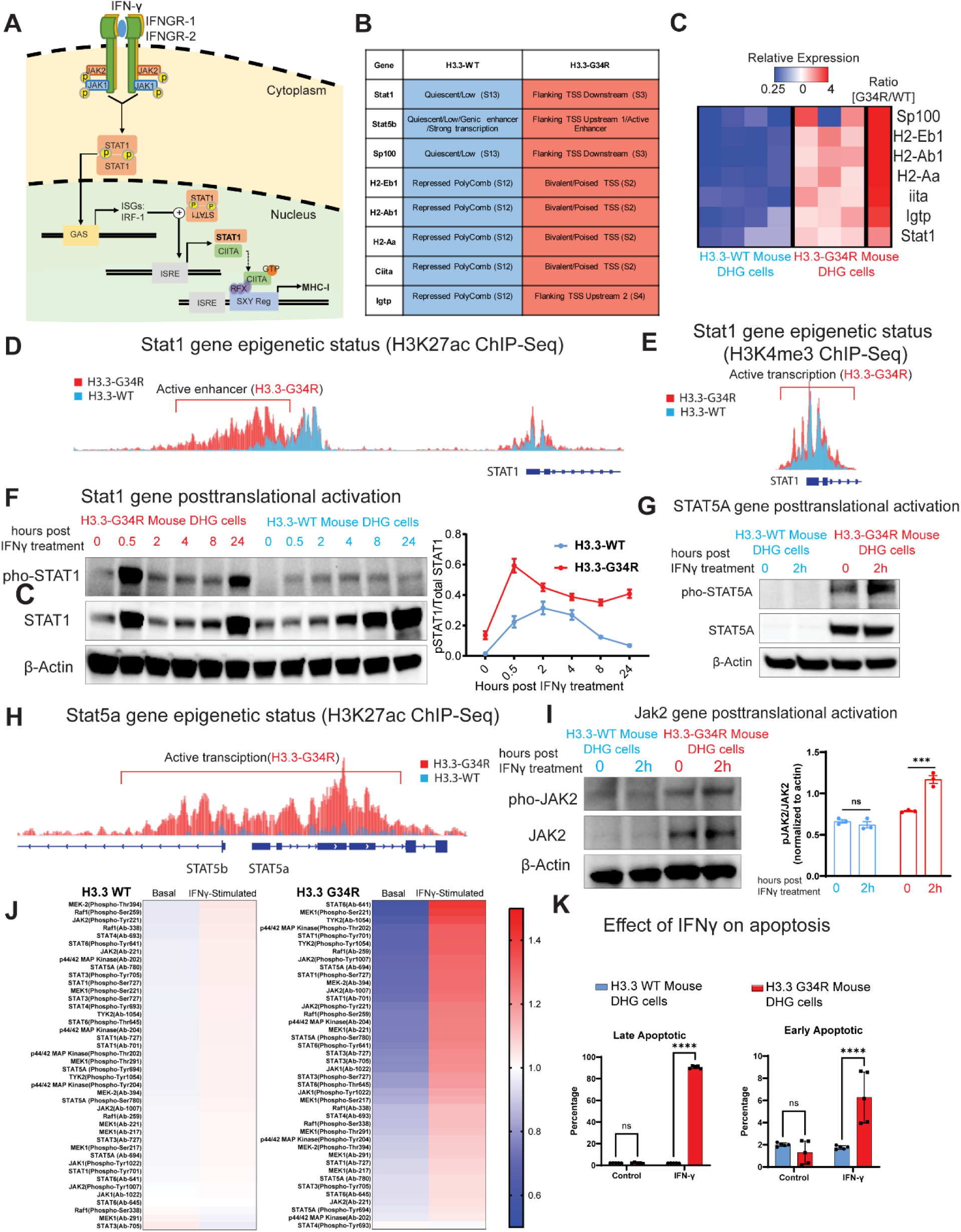
Epigenetic activation of the STAT pathway in G34R DHG. **(A)** Diagram depicting IFNγ inducing STAT1 pathway activation. GAS: interferon-gamma activated site; ISRE: interferon-stimulated response element; SXY reg: SXY regulatory module of MHC promoters; RFX: DNA-binding complex. **(B)** The combination of different histone mark deposition enabled the definition and the identification of chromatin states (CS). The table shows CS changes at specific genomic regions for STAT1 pathway related genes, based on the combination of different histone mark enrichments, considering a fold-change enrichment cut-off > 2 (n=3 biological replicates). TSS=transcription start site. **(C)** Heatmap illustrates RNA-Seq data of H3.3-WT (n=4) and H3.3-G34R (n=3) tumor cells from SB tumors. **(D-E)** Occupancy of H3K27Ac and H3K4me3 on the *Stat1* gene. **(F)** Western blot of phospho-STAT1 and STAT1 at different time points after stimulation with IFN-γ, in H3.3-WT and H3.3-G34R mouse DHG cells. **(G)** Western blot of phospho-STAT5a and STAT5a at basal levels and upon stimulation with IFN-γ in H3.3-WT and H3.3-G34R mouse DHG cells. **(H)** Occupancy of H3K27Ac on the *Stat5a* and *Stat5b* genes. **(I)** Western blot of phospho-JAK2 and JAK2 at basal levels and upon stimulation with IFN-γ in H3.3-WT and H3.3-G34R mouse DHG cells. **(J)** Heatmap of a protein array showing the relative normalized PTM levels of JAK/STAT related proteins in G34R mouse DHG cells versus H3.3-WT cells, highlighting the main upregulated and downregulated proteins and PTMs. **(K)** H3.3-G34R or H3.3–WT neurospheres were incubated with IFNγ (200 UI) or diluent (control) and apoptosis was measured at 48 hours by Annexin V and DAPI staining. *P < 0.05, **P < 0.01, ***P < 0.005, and ****P < 0.001, unpaired t test.

One of these genes is the transcription factor interferon-regulatory factor 1 (*IRF1*), which binds to interferon-stimulated response elements (ISRE) and enhances the transcription of several secondary response genes, such as Class II transactivator (*CIITA*) and Suppressor of cytokine signaling (*SOCS*) which negatively regulates the IFN-γ pathway by inhibiting JAKs and STAT1 phosphorylation (Figures 3A and S7, A)(44). CIITA is the mediator of IFNγ-inducible transcription of Major Histocompatibility Complex (MHC) class I and II genes(44,46). We found differences in the binding of single histone marks on genes related to STAT-pathway GOs (Figures 2, B-C and S7, B-E). The analysis of CS changes between H3.3-G34R and H3.3-WT DHG cells revealed transcriptionally-permissive CS for genes related to STAT1 pathway and IFNγ signaling in H3.3-G34R cells, which correlated with the transcriptional data obtained from the RNA-Seq analysis (Figure 3, B-E) *i.e.*, genes that were shown to be downregulated in H3.3-WT vs H3.3-G34R DHG cells by RNA-Seq, were associated to regions with CS related to transcriptional repression in H3.3-WT cells.

Overall, the presence of H3.3-G34R has profound effects on the epigenetic landscape of DHG cells, which is reflected in the distinctive transcriptional profile observed in these tumors by RNA-Seq. Specifically, we observed that H3.3-G34R expression impacts pathways related to immune system processes and IFNy signaling.

### The JAK/STAT pathway activity is increased in G34R-mutant DHG after IFNγ stimulation

As the RNA-seq performed on tumor cells isolated from the TME showed upregulation of STAT genes and gamma-interferon activated pathways, we hypothesized that interferon-gamma and other cytokines in the G34-mutant DHG TME stimulate the JAK/STAT pathway. Thus, we evaluated STAT1-phosphorylation (Tyr-705), STAT5a phosphorylation (phospho-Ser780) and JAK2 levels in H3.3-G34R DHG and H3.3-WT mouse and human DHG cells by Western blot. Our results show that normalized levels of phospho-STAT1-Tyr-705 are increased in both mouse and human H3.3-G34R DHG cells compared to their H3.3-WT counterparts under basal conditions and following IFNγ stimulation (Figure 3, F, S8, A-B). Similarly, we observed elevated JAK2 phosphorylation (phospho-Tyr1007/1008) normalized to total JAK2, as well as increased STAT5a phosphorylation (phospho-Ser780) normalized to total STAT5a, in H3.3-G34R DHG cells relative to H3.3-WT DHG cells upon IFNγ stimulation (Figure 3, G-I, S8, C-D).

To characterize in detail the JAK/STAT pathway activity in H3.3-G34R and H3.3-WT DHG cells, we analyzed the levels of proteins and their posttranslational modifications using an antibody array with 42 antibodies related to the JAK/STAT pathway. We characterized the levels of posttranslational modifications and basal proteins in DHG cells in basal conditions and after stimulation with IFNγ. Following stimulation with IFNγ for 2 hours, the levels of phosphorylation in proteins associated with posttranslational activation of the JAK/STAT pathway were significantly higher in H3.3-G34R cells compared to H3.3-WT cells (Figure 3, J) (with total STAT6, MEK (Phospho-Ser221),total TYK2, p44/42 MAP kinase (Phospho-Tyr202), STAT1 (Phospho-Tyr701), TYK2 (Phospho-Tyr1054), total Raf1, JAK2 (Phospho-Tyr1007), STAT5A (Phospho-Ser780), STAT1 (Phospho-Ser727), total JAK2, total STAT1, STAT6 (Phospho-Tyr-641), STAT3 (Phospho-Ser727), total STAT4 are differentially upregulated (Q value <0.05) in IFNγ-treated H3.3-G34R vs H3.3-WT DHG cells) (Figure S9, <u>Table S3</u>). These results indicate that G34R DHG cells are more sensitive to IFNγ stimulation, which leads to posttranslational activation of the JAK/STAT pathway.

### H3.3-G34R DHG cells are sensitive to IFNy stimulation and IFNy-mediated apoptosis

IFNγ, and IFNs in general, are known to have anti-tumor effects(47). The anti-tumor effects of IFNγ can be either indirect or direct. Indirectly, this cytokine is a powerful mediator of innate and adaptive immune response; whereas directly, it can inhibit tumor cell proliferation and induce tumor cell apoptosis(42–44,48). Thus, we aimed to test if H3.3-G34R DHG cells were also more sensitive to IFNγ-induced apoptosis. We incubated H3.3-G34R or H3.3-WT DHG cells with 200 UI of IFNγ and then, the proportion of apoptotic cells was assessed by Annexin V/DAPI staining (Figure 3, K). After IFNγ treatment, we observed a significant increase in the proportion of early apoptotic and late apoptotic cells in the H3.3-G34R group (Figure 3, K, S10). These results demonstrate that H3.3-G34R DHG cells are more sensitive to IFNγ stimulation and prone to IFNγ-induced apoptosis. Moreover, IFNγ-mediated apoptosis was confirmed through a caspase 3/7 activity assay performed on H3.3-G34R and H3.3-WT DHG cells after IFNγ treatment (Figure S11, A-B). To ensure this effect is consistent regardless of the cells’ genetic background, we employed a model independent of Ras pathway activation. Specifically, in this model, tumors were induced using shRNA against Tp53 and Atrx, along with the expression of H3.3-G34R and a constitutively active mutant Platelet-derived growth factor receptor A (PDGFRα) (D842V) in cyclin-dependent kinase inhibitor 2A (CDKN2A) knockout mice (Figure S12, Figure S11, C-D). Our results also revealed a significant induction of caspase 3/7 activation in human H3.3-G34R DHG cells compared to both human H3.3-WT cells and normal human astrocytes following IFNγ treatment (Figure S11, E-H).

### JAK/STAT activation leads to MHC-I expression in G34R-mutant DHG

JAK/STAT signaling and CIITA activation were shown to activate MHC I expression, as CIITA is a critical transcriptional regulator of the MHC class I and II complexes(49). MHC-I expression on tumor cells can drive antitumor immunity via antigen presentation. We evaluated CIITA and MHC-I levels in H3.3-WT and H3.3-G34R DHG mouse and human cells by Western blot. In line with ChIP-seq and RNA-seq results, we observed increased expression of these proteins in G34-mutant mouse (Figure 4, A-D) and human (Figure S13) DHG cells. We confirmed these results by assessing MHC-I expression by flow cytometry in mouse and human H3.3-WT and H3.3-G34R DHG cells, observing increased MHC-I expression in G34-mutant DHG cells (Figure 4, E-H). We then compared the levels of MHC I genes in H3.3-G34-mutant and H3.3-WT DHG cells by real-time PCR (qPCR) in our two mouse models (Figure S14, A-B). The levels of MHC class I genes upregulation were also confirmed in patient derived H3.3-G34-mutant DHG cells versus H3.3-WT DHG cells (Figure S14, C).

**Figure 4.**
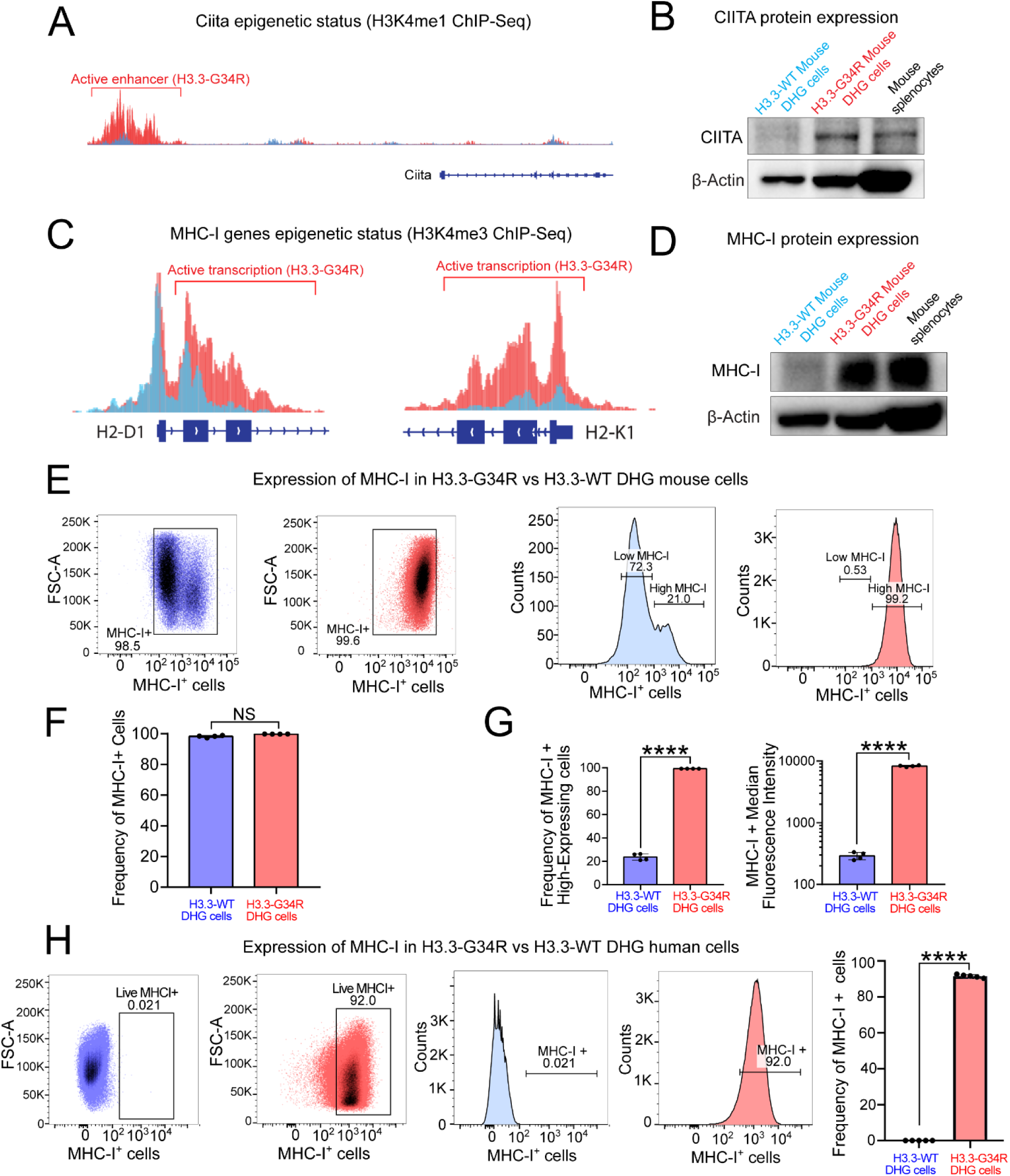
Expression of MHC-I in H3.3-G34R vs H3.3-WT DHG. **(A)** Occupancy of H3K4me1 on the *Ciita* gene. The y axis of each profile represents the estimated number of immunoprecipitated fragments at each position normalized to the total number of reads in each dataset. **(B)** Western blot for CIITA at basal levels and upon treatment with IFN-γ in H3.3-WT and H3.3-G34R mouse DHG cells. Mouse splenocytes were included as positive control. **(C)** Occupancy of H3K4me3 on the MHC-I genes H2-D1 and H2-K1. **(D)** Western blot for MHC-I at basal levels and upon treatment with IFN-γ in H3.3-WT and H3.3-G34R mouse DHG cells. **(E)** Expression of MHC-I in mouse DHG cells by flow cytometry. **(F-G)** Quantitative analysis of the experiment shown in (E). **(H)** Expression of MHC-I in mouse DHG cells by flow cytometry and quantification of the experiment. ****P < 0.001, unpaired t test.

### H3.3-G34R expression impacts on the immune cell infiltration into the glioma microenvironment

The molecular analysis of our H3.3-G34R GEMM indicates that there is an active crosstalk between tumor cells and the immune cells in the TME. Thus, we analyzed the transcriptional differences among immune cells (*i.e.*, CD45+ cells), purified from the H3.3-G34R and H3.3-WT DHG models. At symptomatic stage, tumors were dissected from the mouse brain; a cell suspension was prepared; and CD45+ cells were isolated from H3.3-G34R or H3.3-WT DHG by magnetic sorting (38). Total RNA was purified from the CD45+ cells and gene expression analysis were performed by RNA-Seq. The statistical analysis revealed 243 differentially expressed genes (adjusted p value < 0.05) in CD45+ cells purified from H3.3-G34R tumors with respect to H3.3-WT tumors (Figure 5, A-B; <u>Table S4</u>). We observed that genes related to the presence of T cells, such as *CD3g* and *CD3e*, and specifically to cytotoxic CD8 T cells, such as *CD8a* and *CD8b1*, were significantly upregulated in immune cells from H3.3-G34R DHG (Figure 5, A). On the contrary, we found that the *CD34* marker, which is related to hematopoietic progenitor and stem cells(50), and genes associated with immunosuppression, such as *Arg1*(51), were downregulated in CD45+ cells derived from H3.3-G34R DHG when compared to CD45+ cells derived from H3.3-WT DHG (Figure 5,C), implying that the immune cells of the H3.3-G34R DHG TME are more differentiated and less immunosuppressive than the immune cells from the H3.3-WT TMEs.

**Figure 5.**
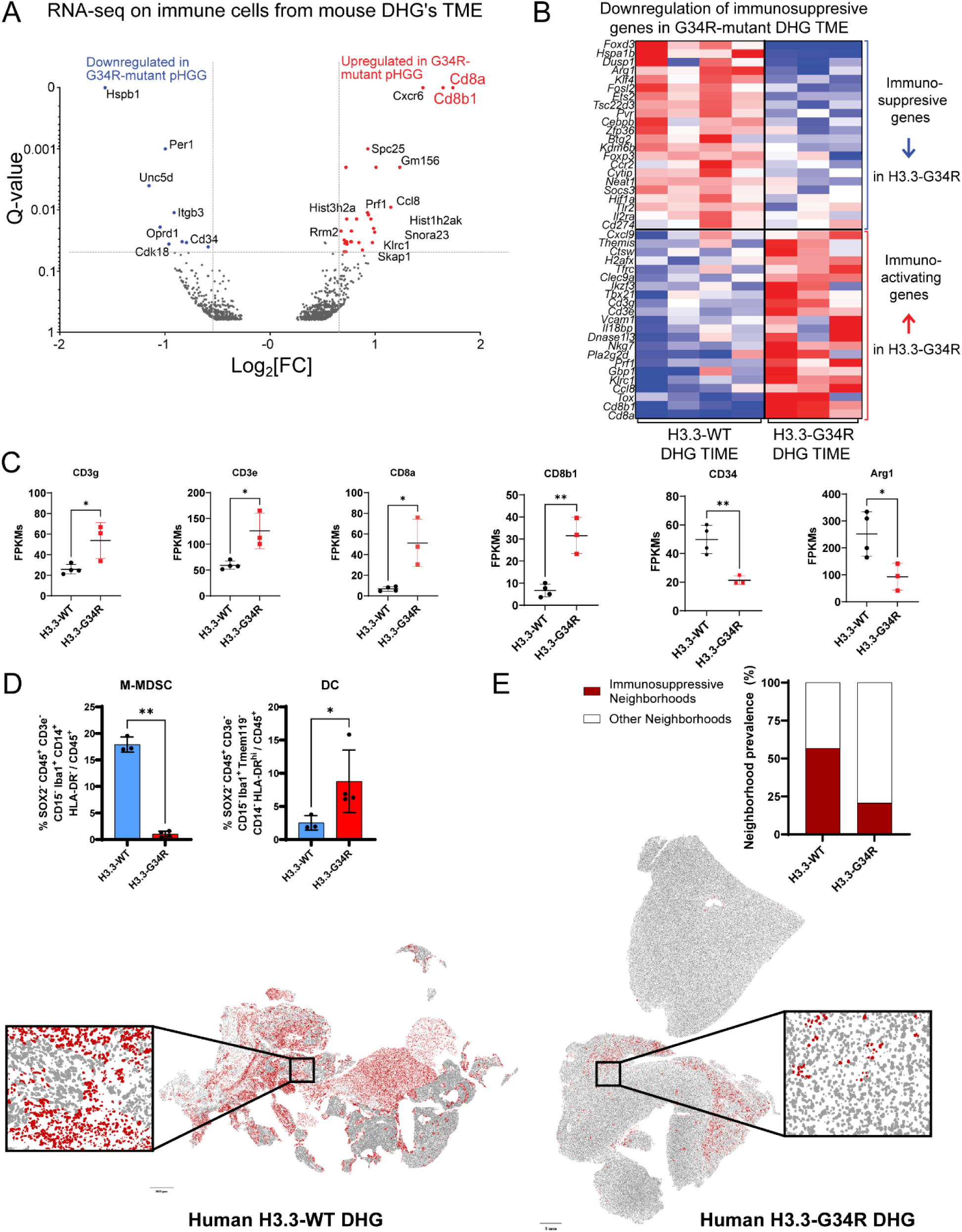
Analysis of H3.3-G34R DHG TME immune cells. **(A)** Volcano plot shows differentially expressed genes (-Log(p-value), y axis) versus the magnitude of change or fold change (Log2(Fold Change), x axis) when comparing gene expression in CD45 positive cells derived from H3.3-G34R versus CD45 from H3.3-WT SB-generated tumors. **(B)** Heatmap shows the expression level of genes related to immunosuppression in CD45+ cells obtained from H3.3-G34R and H3.3-WT SB-derived tumors. **(C)** Differentially expressed genes related to T-cell activation (CD3g, CD3e, CD8a and CD8b1), immunosuppression (Arg1) and stemness (CD34) were identified, between H3.3-WT (n=4) and H3.3-G34R (n=3) CD45+ cells. Expression levels are shown as Fragments Per Kilobase of transcript per Million mapped reads (FPKMs, y axis) ± SD. **(D)** The proportion of cells in human samples identified as DC (SOX2^-^ CD45^+^ CD3e^-^ CD15^-^ Iba1^+^ Tmem119^-^ CD14^-^ HLA-DR^hi^ / CD45^+^) and M-MDSC (SOX2^-^ CD45^+^ CD3e^-^ CD15^-^ Iba1^+^ CD14^+^ HLA-DR^-^ / CD45^+^). **(E)** Neighborhood analysis identifying immunosuppressive neighborhoods as those enriched in M-MDSC, PMN-MDSC or M2 macrophages. Plots illustrate representative samples from each experimental group with cell color corresponding to their neighborhood type. The bar graph displays the average prevalence (%) of each neighborhood for each experimental group.

Then, we performed a network analysis and clustering of the upregulated genes in H3.3-G34R-derived CD45+ cells using the Reactome Functional Interaction plugin in Cytoscape (52). After network building and clustering, we detected 5 clusters with highly interacting nodes (Figure S15, A). The analysis of the top-30 most significant pathways associated to these clusters displayed terms that were related to immune system activation, especially to T cell-mediated responses (Figure S15, A). In summary, the transcriptional analysis of CD45+ cells isolated from the TME of H3.3-G34R DHG showed the upregulation of genes related to an activated T cell response. To assess the immune cell infiltration in human DHGs, multiplex immunocytochemical analysis of the cells in three H3.3-WT and four H3.3-G34R human DHG samples was performed (Figure S15, B). There was a statistically significant decrease in M-MDSCs, as well as a statistically significant increase in DCs in the H3.3-G34R patient samples (Figure 5, D). Clustering neighborhoods of similar composition into regions resulted in seven distinct neighborhood types in all tissue samples. Four of these neighborhoods were enriched in immunosuppressive cell types in the H3.3-WT samples, which comprised 56.7% of the total neighborhoods (Figure 5, E). Only two of these neighborhoods were enriched in immunosuppressive cell types in the H3.3-G43R samples, which made up 20.6% of the total neighborhoods in those tissues (Figure 5, E). Overall, immunosuppressive neighborhoods accounted for much more of the tissue in H3.3-WT samples compared to H3.3-G34R samples.

To assess the role of T cells in G34-mutant DHG development, we compared the survival of C57BL/6, *Rag-1* KO (devoid of T and B-cells(53)), and Cd8 KO (devoid of CD8^+^ T-cells) mice, implanted with H3.3-WT or H3.3-G34R DHG cells. We observed a reduced survival of *Rag-1* KO (Figure S16, A) and *Cd8* KO (Figure S16, B) mice that were implanted with H3.3-G34R cells, in comparison to C57BL6. In the case of mice implanted with H3.3-WT DHG cells, there was no survival difference between C57BL/6 and *Rag-1* KO; and the survival difference between C57BL/6 and *Cd8* KO mice was only of 4 days. The marked reduction in the MS observed in mice bearing G34-mutant DHG demonstrates the relevance of cytotoxic T cell mediated anti-tumor activity in this model of DHG. Additionally, we implanted the CDKN2A KO/PDGFRA D842V/p53 KD/ATRX KD H3.3-WT or H3.3-G34R cells in NOD scid gamma (NSG) mice. Our results show that in this immunocompromised host there is no difference in the MS between H3.3-WT and H3.3-G34R DHG (Figure S16, C).

To assess the immune cell infiltration pattern in H3.3-G34R DHG and uncover how the presence of the H3.3-G34R mutation affects the TME, we implanted H3.3-G34R or H3.3-WT cells in the brain of C57BL6 mice. At symptomatic stage, DHG were dissected and processed for spectral flow cytometry analysis (Figure 6, A). UMAP embedding of the immune cell spectral flow cytometry (SFC) analysis revealed distinct clustering for H3.3-G34R Vs. WT DHG immune cells (Figure 6, B). MDSC phenotypic characterization also requires distinction between Monocytic (M) (CD45+ CD11b+ Ly6C^hi^ Ly6G^-^) MDSC and Polymorphonuclear (PMN) (CD45+ CD11b+ Ly6C^lo^ Ly6G^+^) MDSC. We found similar proportions of cells displaying PMN-MDSC phenotype in the TME of both tumors, but a significant decrease in the proportion of M-MDSCs in the TIME of H3.3-G34R DHG (Figure 6, C-D). For T cell analysis, pan T cells were identified as CD45^+^/CD3^+^ cells, and were classified into CD8^+^ or CD4^+^ T cells (CD45^+^/CD3^+^/CD8^+^ or CD45^+^/CD3^+^/CD4^+^, respectively), a significant increase in the proportion of CD4 T cell infiltrates were observed in H3.3-G34R DHG TIME (Figure 6, E-F). Microglia (CD45^low^/CD11b^+^/TMEM119^+^/P2Y12^+^) proportion was also significantly increased in H3.3-G34R TIME (Figure 6, G). Additionally, we measured CD206 and MHCII expression on macrophages to classify them as anti-tumor M1 (CD45^+^ F4/80^+^ Ly6C^-^ CD206^-^ MHCII^+^) or pro-tumor M2 macrophages (CD45^+^ F4/80^+^ Ly6C^-^ CD206^+^ MHCII^-^)(54). We observed that the proportion of M2 macrophages significantly decreased in H3.3-G34R TIME (Figure 6, H). Additionally, spectral flow cytometry analysis was also performed in CDKN2A KO/PDGFRA D842V/p53 KD/ATRX KD H3.3-WT or H3.3-G34R tumors, inwhich we observed similar results (Figure S17).

**Figure 6.**
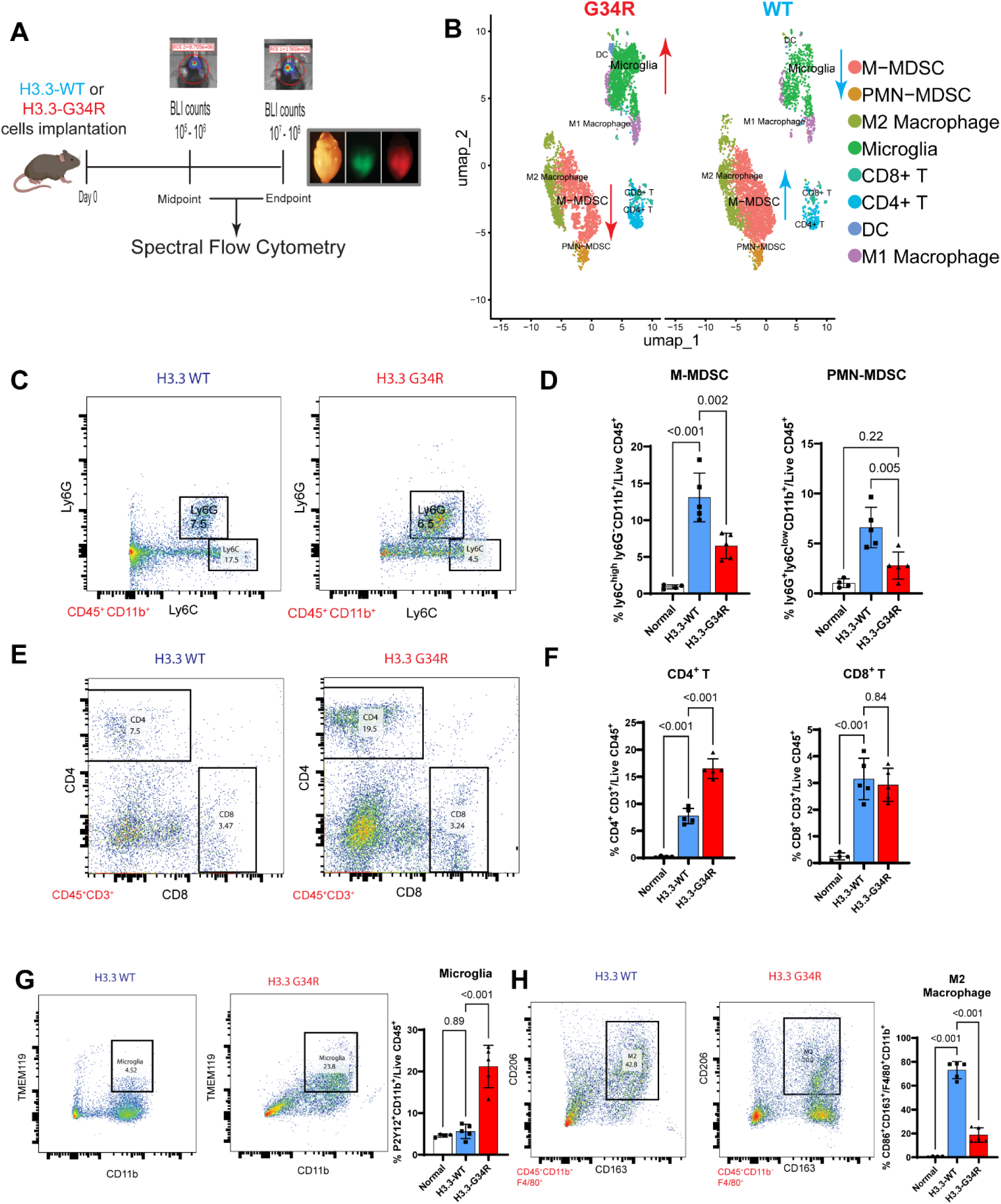
Spectral flow cytometry Analysis of the tumor immune microenvironment (TIME) of H3.3-WT and H3.3-G34R DHG at end stage. **(A)** Illustration of experimental procedure followed for the characterization of immune cells in H3.3-WT and H3.3-G34R mouse DHG TIME. **(B)** UMAP embedding of H3.3 G34R and WT DHG immune cell SFC analysis. **(C-D)** The proportion of cells with Monocytic (CD45+/CD11b+/Ly6G-Ly6Chi) or Polymorphonuclear (CD45+/CD11b+/Ly6G+Ly6Clo) myeloid derived suppressor cell. **(E-F)** Total pan T cells were identified as CD45+/CD3+ cells and further classified by the expression of CD8 or CD4 (CD45+/CD3+/CD8+ or CD45+/CD3+/CD4+, respectively). **(G)** The total proportion of macrophages is shown as the percentage of CD45+/CD11b+/F4/80+ cells. **(H)** The total proportion of microglia is shown as the percentage of CD45 low/CD11b+/TMEM119+ P2Y12+ cells. Data are represented as mean±SD. Statistical differences determined with one-way ANOVA Tukey’s post hoc test. **** p<0.0001; ***p<0.001, **p<0.01, * p<0.05, ns, not significant.

To rule out the effect on the TIME of the fluorescent protein, Katushka. We developed new cells expressing either the fluorescent marker Katushka alone or in conjunction with wild-type histone H3.3 (H3.3-WT), and we performed a detailed analysis of the immune TIME in these models (Figure S18). The analysis showed no significant differences attributed to Katushka expression alone or H3.3-WT with Katushka (Figure S19). These results confirm that the fluorescent marker Katushka does not interfere with the assessment of the TIME in our mouse model. This comprehensive approach ensures that any observed alterations in the TIME are due to biological changes in the model itself and not artifacts introduced by expression of fluorescence markers. To address the concerns regarding differences in growth rate between G34-mutant and H3.3-WT DHG tumors, we performed a characterization of the TIME in our mouse models at the same post-implantation time point. This ensured that the samples were collected when animals exhibited comparable signs of tumor burden. The immune cell infiltration differences are similar at midpoint in our two mouse models (Figure S20-21). These detailed analyses suggest that the differences in the TIME observed in H3.3-G34R versus H3.3-WT DHG tumors are not attributed to dissimilarities in tumor growth rates.

Taken together, the analysis of the TIME revealed that H3.3-G34R DHG tumors are infiltrated with a lower portion of immunosuppressive cells, contributing to a more immune-permissive tumor microenvironment.

### H3.3-G34R impacts the array of cytokines secreted by DHG cells, rendering the TME less immunosuppressive

Given that H3.3-G34R DHG cells display a transcriptional landscape and a TME infiltration pattern consistent with an activated immune response, analyzed how the presence of H3.3-G34R alters the array of cytokines secreted by H3.3-G34R DHG cells. We analyzed H3.3-WT or H3.3-G34R cell culture supernatants for cytokine concentration (Figure 7A). The secretion of G-CSF, VEGF, MCP-1, MCP-3, TGFβ and N-Gal was lower in H3.3-G34R DHG cells. The downregulation of G-CSF, VEGF and N-Gal were also confirmed in human H3.3-WT and H3.3-G34R DHG culture supernatants (Figure S22, A). We investigated our ChIP-Seq data to analyze the chromatin context of the genes encoding the differentially secreted cytokines. We observed an increase in H3K27me3 (transcriptional repressive) mark deposition on the promoter region of *CSF3* (gene that produces G-CSF) in H3.3-G34R DHG cells with respect to H3.3-WT. On the contrary, H3K4me1, H3K4me and H3K27ac (transcriptional activation) marks were found to be increased in the same region in H3.3-WT vs H3.3-G34R DHG cells (Figure 7, B). These results indicate the presence of a permissive chromatin for G-CSF expression in H3.3-WT with respect to H3.3-G34R DHG cells. We did not find differences in the deposition of histone marks for the other cytokines with significantly different concentrations.

**Figure 7.**
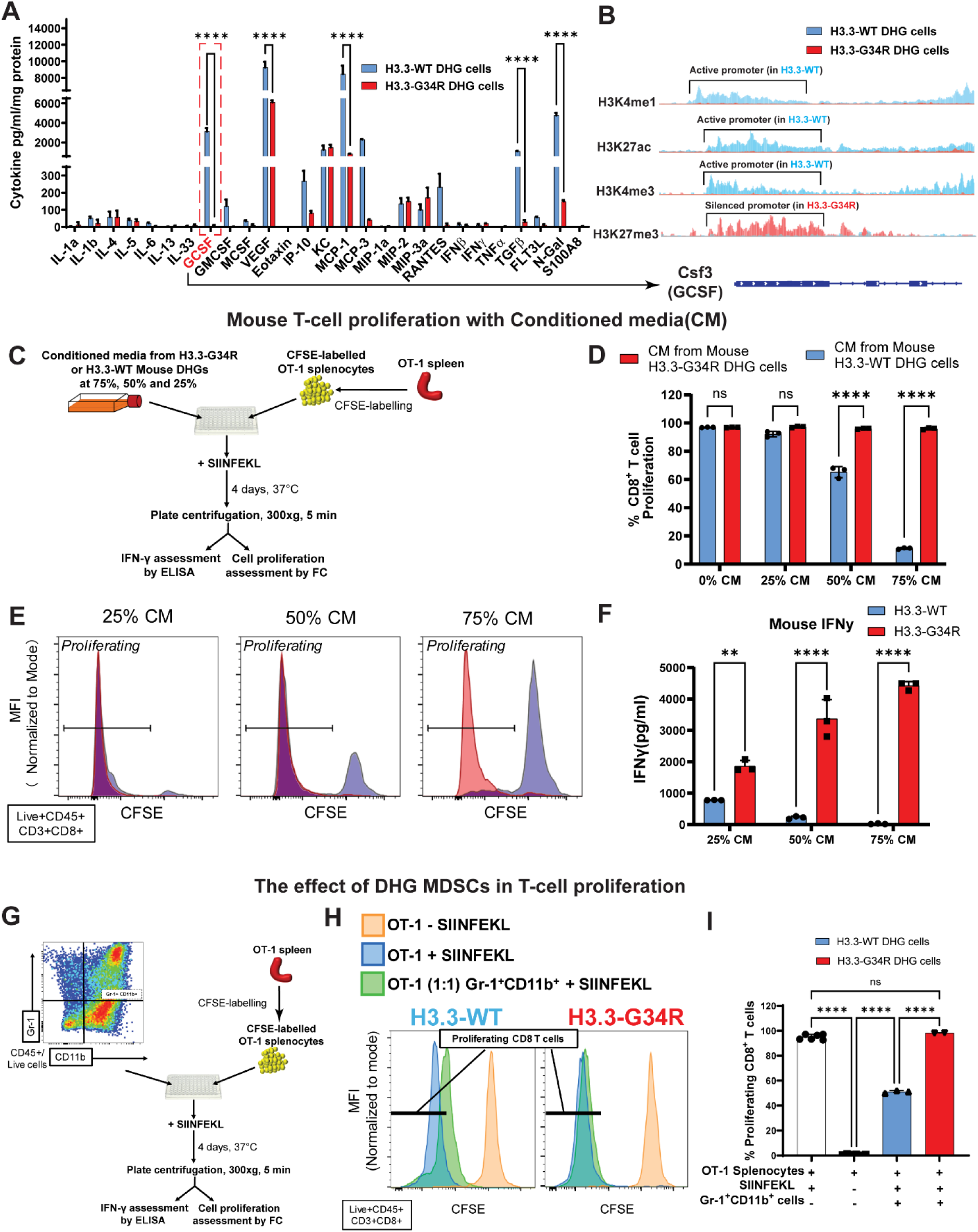
Cytokines profile of G34R DHG cells and immune properties of the conditioned media. **(A)** The concentration of different cytokines in the conditioned media from three different clones of H3.3-WT and H3.3-G34R neurospheres was analyzed by ELISA. ****p<0.0001, 2-way ANOVA and Sidak’s multiple comparisons test. **(B)** The plots show the occupancy of H3K27me3 (transcriptional repressing), H3K27ac, H3K4me3 and H3K4me1 (transcriptional activating) histone marks on the G-CSF (*Csf3*) promoter region (light blue dashed box). **(C)** Experimental layout of the T cell proliferation assay to test the immunosuppressive capacity of conditioned media (CM) from H3.3-G34R or H3.3-WT neurospheres. **(D)** The percentage of proliferating CD8+ T cells in the presence of CM from H3.3-WT or H3.3-G34R neurospheres, with respect to the positive control (0% CM). **(E)** Representative histograms show the CD8+ T cell proliferation in the presence of 25%, 50% or 75% CM from H3.3-WT and H3.3-G34R neurospheres. **(F)** IFNγ concentration was measured in the supernatant of the cultures by ELISA. Bars represent cytokine concentration in absolute numbers. **(G)** Experimental layout of the T cell proliferation assay to test the immunosuppressive capacity of Gr-1+ CD11b+ cells isolated from H3.3-WT and H3.3-G34R DHG. **(H)** T cell proliferation was measured as the reduction of CFSE staining in the CD45+/CD3+/CD8+ population with respect to the positive control (OT-1 splenocytes + SIINFEKL). Representative histograms showing the CD8+ T cell proliferation in the presence of Gr-1+ CD11b+ cells from H3.3-WT and H3.3-G34R TMEs. **(I)** Bars represent the percentage of proliferating CD8+ T cells in each treatment. **** p<0.0001, *** p<0.001; ns=non-significant; ANOVA and Tukey test. **** p<0.0001; ns=non-significant; ANOVA and Tukey test; mean ± SEM.

To assess the impact of the cytokines secreted by H3.3-G34R DHG cells on T cell survival, we performed T cell proliferation assays in the presence of conditioned media (CM) from H3.3-WT and H3.3-G34R DHG cells (Figure 7, C). The addition of increasing amounts of H3.3-WT CM correlated with decreasing proportions of proliferating CD8 T cells, whereas no inhibition in CD8 T cell proliferation was observed in the presence of H3.3-G34R CM (Figure 7, D-E). In accordance, IFNγ secretion decreased with the addition of increasing amounts of H3.3-WT CM, whereas IFNγ secretion increased with the addition of increasing amounts of H3.3-G34R CM (Figure 7, F). Additionally, we performed T cell proliferation assays in the presence of CM from H3.3-WT and H3.3-G34R human DHG cells using human PBMCs (Figure S22, B). The presence of H3.3-G34R CM was associated with higher proportions of proliferating CD8 T cells compared to H3.3-WT CM (Figure S22, C-D). Consistently, IFNγ secretion increased with the addition of H3.3-G34R CM compared to H3.3-WT CM (Figure S22, E).

As we discovered these differences in cytokine secretion and their implications on T cell proliferation, we analyzed the impact of H3.3-G34R or H3.3-WT DHG-derived factors in the immunosuppressive functionality of CD11b^+^ Gr-1^+^ MDCS cells encountered in the TME. To assess the immunosuppressive ability of the MDCS cells, we performed T cell proliferation assay(55) (Figure 7, G). Whilst the CD11b^+^ Gr-1^+^ cells from H3.3-WT inhibited T cell proliferation by 50%, CD11b^+^ Gr-1^+^ cells from H3.3-G34R did not affect it (Figure 7, H-I). These data demonstrates that the presence of H3.3-G34R impacts the array of cytokines secreted by tumor cells, creating a less immunosuppressive TME with respect to the H3.3-WT tumor cells. We observed that these secreted factors enable the proliferation of effector T cells and do not induce the production of *bona fide* MDSC or their homing into the TME.

### TK/Flt3L immune-stimulatory gene therapy enhances anti-glioma immune response in H3.3-G34R DHG mouse model

As we observed an immune-permissive tumor microenvironment associated with the expression of H3.3-G34R in the DHG mouse model, we evaluated the efficacy of a cytotoxic immune-stimulatory TK/Flt3L gene therapy (GT). This therapy entails injecting adenoviruses encoding Thymidine kinase (TK) and FMS-like tyrosine kinase 3 ligand (Flt3L) into the tumor, followed by ganciclovir (GCV) administration(25,26). The TK converts GCV into a competitive inhibitor of DNA synthesis, which leads to tumor cell lysis and the concomitant release of tumor antigens(26,28). Flt3L mediates DC recruitment and expansion in the TME, which uptake the tumor antigens, traffic them to the draining lymph nodes and prime a robust anti-tumor cytotoxic and memory T cell response, leading to tumor regression and long-term survival (20,25,26,28).

Thus, we treated H3.3-WT and H3.3-G34R DHG-bearing animals with the TK/Flt3L GT or saline as depicted in Figure 8, A. The mice harboring H3.3-WT gliomas that were treated with saline had a MS=20 days, mice that received GT exhibited a longer MS=29 days and around 20% long-term survivors (Figure 8, C). In contrast, we observed a significant extension in the survival of animals harboring H3.3-G34R gliomas treated with GT (MS=Not reached), with respect to the control group treated with saline (MS=44 days post-injection (DPI)) (Figure 8, B-C). GT led to an effective cure rate of 53.3% (Figure 8, C). To assess the safety of this therapy, complete blood cell counts and serum chemistry analysis of H3.3-G34R DHG-bearing mice treated with saline or GT was performed (Figure S23, A). We did not find any differences in the amount of white blood cells (WBC) and red blood cells (RBC); the percentage of hemoglobin; or in the hematocrit between the saline and GT treated groups (Figure S23, A). Also, we observed that the levels of AST (Aspartate transaminase), ALT (Alanine transaminase), Creatinin, and BUN (Blood urea nitrogen) were within the normal range for mice treated with GT, indicating that the liver and the renal system were functioning normally (Figure S23, A). The histological assessment of livers from animals treated with saline or GT did not display any abnormalities (Figure S23, B). Additionally, IHC analysis was performed to assess the brain anatomy on paraffin-embedded brain tissue from saline or long-term survivors from gene therapy treated animals. Sections were stained for Myelin Basic Protein (MBP) (myelin sheath marker, to assess neuronal structure), Glial fibrillary acidic protein (GFAP) (Glial marker, to assess glial inflammation) and Nestin (neural stem cell marker which stains for tumors cells). The analysis indicates that long term survivors have no signs of anatomical abnormalities or signs of tumor growth (Figure S23, C). We also used CPAH (CDKN2A^−/−^/PDGFRα/shTP53/shATRX/H3.3-G34R) implanted mice treated with saline or Flt3L-TK gene therapy. The saline group showed a MS of 23 days, while the group treated with the immunostimulatory gene therapy showed a significantly improved outcome, with the median survival not reached and 60% of long-term survivors (p=0.0256). (Figure S24).

**Figure 8.**
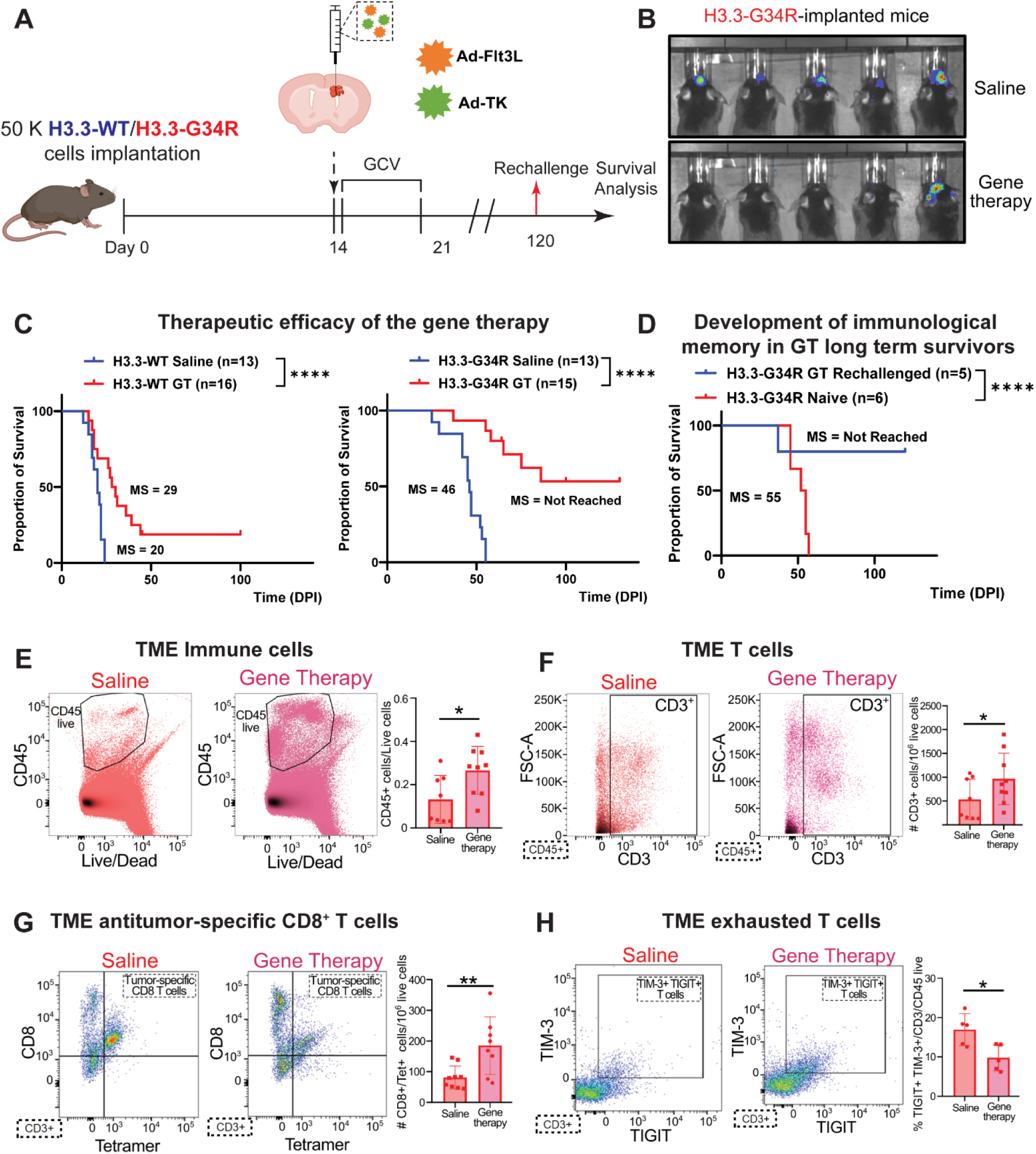
Evaluation of a GCV-inducible gene therapy for H3.3-G34R DHG. **(A)** Diagram depicting the experimental outline for the treatment of H3.3-WT and H3.3-G34R tumor bearing mice with Ad-Flt3L + Ad-TK gene therapy (GT). **(B)** IVIS scan of animals bearing H3.3-G34R tumors 14 days after being treated with Saline or GT. **(C)** Kaplan-Meier survival curve for the GT experiment. Median survival (MS) and number of animals used are detailed in the plot legend. **** p<0.0001, Mantel-Cox test **(D)** Long-term survivors were implanted in the contralateral hemisphere with 50.000 H3.3-G34R cells and animals were euthanized at symptomatic stage. **(E)** Percentage of CD45 infiltration in the TME of H3.3-G34R tumor bearing mice treated with Saline or GT. **(F)** Number of T cells (CD45+ CD3+) in the TME of H3.3-G34R tumor bearing mice treated with Saline or GT. **(G)** Number of tumor-specific CD8 T cells (CD45+ CD3+ CD8+ Tet+) in the TME of H3.3-G34R tumor bearing mice treated with Saline or GT. **(H)** Percentage of double positive T cells for TGIT and TIM-3, two T cell exhaustion markers. * p<0.05; **p<0.01 Student T test.

As tumors remained undetectable by luminescence for more than 100 DPI in the GT-treated long-term survivors, we rechallenged these animals in the contralateral hemisphere with H3.3-G34R DHG cells without further treatment to examine the development of immunological memory. As controls, naïve animals were implanted with H3.3-G34R DHG cells in the striatum. Most of long-term survivors from the GT experiment did not develop tumors after rechallenging at day 90 DPI, whereas those from the control group succumbed due to tumor burden (Figure 8, D). Rechallenged animals were euthanized at 90 DPI, perfused, and the brains were paraffin embedded and sectioned for histological analysis (Figure S25, A). After H&E staining, no tumor mass was detected in the rechallenged animals (Figure S25, A). Also, the histological assessment of the liver of these mice did not show any abnormalities (Figure S25, B). Additionally, serum metabolites were analyzed on the rechallenged animals, showing no alterations in any marker compared to the reference range (Figure S25, C), demonstrating that the gene therapy does not elicit toxic effects. These data demonstrates that TK/Flt3L GT is effective for the treatment of H3.3-G34R HGG and that it prompts the development of immunological memory.

To confirm the presence of anti-tumor specific CD8 T cells in the TME, we generated H3.3-G34R DHG cells expressing the subrogate antigen ovalbumin (OVA). OVA antigen is a model protein for studying antigen-specific immune responses in mice(20,28,56). H3.3-G34R-OVA cells were implanted in the striatum of C57BL/6 mice and at 14 DPI mice were treated with GT or saline, followed by treatment with GCV, twice a day for 7 days. Mice were euthanized at the end of GCV treatment and tumors were dissected to evaluate the immune response. The TME was examined to study the presence of total infiltrating T cells and anti-tumor specific CD8 T cells (Figure 8, E-G). Anti-tumor specific CD8 T cells were identified using SIINFEKL-H2K^b^-Tetramer (Figure 8, G). We observed that the TME of animals treated with GT had a higher amount of CD45+ cells (Figure 8, E) and CD3+ cells (Figure 8, F) compared to the saline group TME. Moreover, GT-treated animals displayed an increased number of tumor-specific CD8 T cells (CD45+ CD3+ CD8+ Tetramer+) than the saline group (Figure 8, G). In addition, we observed a decrease in exhausted T cells (CD45+ CD3+ TGIT+ TIM-3+) in the GT group compared to Saline animals (Figure 8, H).

Collectively, these results demonstrate that the TK/Flt3L GT increases the amount of tumor-specific CD8 T cells in the TME of H3.3-G34R gliomas, enhancing the MS and contributing to the development of anti-tumor immunological memory.

## DISCUSSION

We aimed to determine the biological features of a DHG subtype that harbors a mutation in the *H3F3A* gene, named H3.3-G34R. In the present work, we used an immunocompetent mouse model to better understand the tumor’s molecular characteristics and its immune microenvironment, and to test the effectiveness of an immune-mediated gene therapeutic approach for H3.3-G34R DHG. The immune response in the brain differs from the immune response mounted in other parts of the body due to anatomical particularities of the CNS (57);(58–61). Factors derived from neural cells promote quiescence and immunosuppression in the brain microenvironment (58,59). Nevertheless, in response to inflammatory stimuli, this quiescent immune state pivots, promoting the proliferation of brain resident microglial cells, as well as increasing the permeability of the blood brain barrier for the infiltration of myeloid cells and activated T cells(59,62–64). Glioma TME has been reported to be immunosuppressive(65–68).

Whilst most of the studies related to glioma TME have been performed on adult patient samples, the TME of children and young adult brain tumors has remained understudied, hindering the development of effective immune-mediated therapies. Using our immunocompetent H3.3-G34R DHG model, we were able to understand how the presence of this mutation impacts the epigenomic and transcriptomic landscape of DHG cells, affecting the tumor immune cell infiltration. By RNA-Seq we observed that H3.3-G34R glioma cells had a transcriptional pattern coherent with an active immune response in the TME. Specifically, we detected that H3.3-G34R tumor cells were responding to IFNs, as several genes related to IFN-I (IFNα and IFNβ) and IFN-II (IFNγ) signaling pathways were upregulated in H3.3-G34R DHG cells with respect to H3.3-WT DHG cells (Figure 1). We found the upregulation of genes that are downstream of the IFNγ signaling pathway (*Irf1*, *Ciita* and genes related to MHC-I complex, (Figures 3 and 4)), in H3.3-G34R DHG cells (42,44,46). Moreover, we detected that several genes involved in IFNγ signaling displayed transcriptionally permissive chromatin states (CS) in H3.3-G34R cells, in comparison to the transcriptionally inactive or repressed CS observed for the same regions in H3.3-WT (Figures 3 and 4), demonstrating that this transcriptional pattern is a consequence of the epigenetic signature induced by H3.3-G34R. We also observed an overall posttranslational activation of the JAK/STAT signaling evidenced by phosphorylation arrays upon IFNγ-stimulation. This pathway is widely conserved in mammals, and its activation impacts several biological processes, including cell proliferation, differentiation, and apoptosis(69), and aberrant activation of STAT proteins, particularly STAT1 and STAT5, has been described in glioma, and their activation may be related to oncogenesis(70,71).

Although the microenvironment of H3.3-G34R DHG mouse model is less immunosuppressive than that in H3.3-WT DHG, animals with H3.3-G34R tumors still eventually succumb to tumor burden. In response, we examined the effectiveness of the immune-boosting TK/Flt3L gene therapy (GT) (Figure 8A), a treatment our laboratory developed that has recently completed a phase I clinical trial for adult HGG patients (https://clinicaltrials.gov; identifier: NCT01811992)(20,25–28,72–74). Mice with H3.3-G34R DHG that were treated with this therapy lived longer than those receiving saline, and 53.3% became long-term survivors (Figure 8, C). Furthermore, these survivors remained tumor-free even without additional treatment. We also verified that this therapy increased the infiltration of CD45+ and pan T cells, thereby encouraging the growth and migration of tumor-specific CD8 T cells to the brain (Figure 8, E-G). The presence of CD8 T cells is crucial for combatting cancer as they can kill tumor cells and establish immunological memory, which is particularly critical to prevent tumor recurrence after surgery. However, these T cells can become exhausted and less responsive when continually exposed to antigens in non-ideal conditions, as evidenced by the expression of certain co-inhibitory receptors. In human HGG samples, these exhaustion markers are detectable on infiltrating T cells (75). In this study, we noticed decreased expression of these markers in CD8 T cells from GT-treated animals (Figure 8, H). Overall, the TK/Flt3L GT therapy showed promise in this pre-clinical H3.3-G34R DHG model, prolonging survival and creating anti-tumor immunological memory, while increasing the population of tumor-specific CD8 T cells and potentially delaying or preventing T cell exhaustion.

Our H3.3-G34R DHG GEMM enabled us find for the first time that H3.3-G34R presence affects the crosstalk between the PDHGG and the immune system. We demonstrated that H3.3-G34R DHGs are infiltrated with a pattern of immune cells that is consistent with an anti-tumor response being mounted, which was found to be further enhanced by the application of the TK/Flt3L GT. This immune stimulatory gene therapy (GT) increased the median survival of H3.3-G34R tumor bearing animals and prompted the development of tumor-specific immune response. Translationally, the pre-clinical data exhibited in this work represent robust evidence to consider the TK/Flt3L as a feasible immune-mediated GT for the treatment of H3.3-G34R malignant glioma patients.

Additionally, our results suggest that G34R-mutant DHG might benefit from other immunotherapies such as CAR-T or PD-L1 therapies. This highlights the relevance of the molecular stratification in the treatment decisions. To better understand the tumor microenvironment (TME) of pediatric patients, more research is necessary, particularly to elucidate the disparities attributable to various molecular subgroups. Most current research involving patient samples primarily focuses on tumor cell analysis. However, associating biological features with a specific molecular subgroup is challenging due to the scarcity of suitable control tumors in patients. For example, there are few histone-Wild type DHGs that are hemispherical and possess a genetic background— specifically, P53 mutant and ATRX mutant—comparable to that of G34R-mutant DHGs. In our study, we demonstrate a correlation between the findings from our mouse model and transcriptomic data from patients, specifically noting an observed upregulation of the STAT pathways in G34-mutant DHG patients.

In summary, this study advances our knowledge of the intricate interplay between the H3.3-G34R mutation, its impact on epigenomic and transcriptomic alterations, and the consequent modulation of the tumor immune microenvironment. Our findings underscore a less immunosuppressive tumor microenvironment in H3.3-G34R DHGs, driven by the complex interplay between molecular aberrations and ensuing immune responses. Additionally, our investigation revealed the potential of TK/Flt3L gene therapy in increasing survival and fostering an anti-tumor immune response, thereby providing robust preclinical evidence to consider this approach for clinical translation in H3.3-G34R DHG patients. As we continue to unveil the mechanisms within DHGs, it’s apparent that a better understanding of the tumor microenvironment associated with various molecular subgroups is an area that demands further research. Together, our findings set the stage for more personalized, effective therapeutic strategies, highlighting the crucial role of molecular stratification in clinical decision-making for the implementation of DHG treatments.

## Author Contributions

Conceptualization, M.B.G.F., P.R.L. and M.G.C.; Methodology, M.B.G.F., S.H., K.B. and Z.Z.; Investigation, M.B.G.F., S.H., K.B., Z.Z., B.L.M.,A.A.M.,Y.L.,C.E.T.,M.E.J.W.,J.Y.,P.K.,A.W.T.,F.J.M.,M.S.A.,A.C.,F.M.M.,A.J.N., B.S.,F.M.N. and M.B.E. Software, T.Q. J.D.W., M.A.S. Writing – Original Draft, M.B.G.F., S.H. and Z.Z.; Writing–Review & Editing, Z.Z., M.L.V. P.R.L. and M.G.C.; Funding Acquisition, P.R.L. and M.G.C.; Resources, R.T.C.,M.N.,C.K.,S.V.,M.P.,E.N.,G.P.,M.V.,C.L.K.,N.J.,J.P.C.,A.M.S.,M.D.,D.H.,L.M.P., K.B.M.,G.A.G. and C.K.P. Supervision, P.R.L. and M.G.C.

## Declaration of Interests

The authors declare no competing interests.

## ACKNOWLEDGMENTS

The laboratories of MGC and PRL are supported by the National Institutes of Health (NIH)/National Institute of Neurological Disorders and Stroke (NIH/NINDS) grants R37-NS094804, R01-NS105556, R01-NS122536, R01-NS124167, and R21-NS123879, and Rogel Cancer Center Scholar Award to M.G. Castro; NIH/NINDS grants R01-NS076991, R01-NS082311, R01-NS096756, R01NS122234, and National Institutes of Health/National Cancer Institute (NIH/NCI) R01-CA243916 to P.R. Lowenstein; the Department of Neurosurgery; and The Pediatric Brain Tumor Foundation, Leah’s Happy Hearts Foundation, Ian’s Friends Foundation (IFF), Chad Tough Foundation, Smiles for Sophie Forever Foundation to M.G. Castro and P.R. Lowenstein; and the American Brain Tumor Association Basic Research Fellowship ‘‘in Memory of Bruce and Brian Jackson’’ to M.B.G.F We would like to acknowledge the University of Michigan Epigenomics Core, the Advance Genomics Core, the Flow Cytometry Core, the In-Vivo Animal Core (IVAC) and the UMICH Single Cell Analysis Resource for providing services that contributed to this study.

A.M.S. is supported by NIH R01NS110703, U19CA264338, The Northwestern Nervous System Tumor Bank is supported by the P50CA221747 SPORE for Translational Approaches to Brain Cancer. C.K.P. is supported by BRAF LGG consortium research fund, NCI; Stanford Pediatrics CMDC (Cancer Model Development Center)/Human Cancer Model Initiative; Contract 17X074 and NIH/NCI U54CA261717.

## STAR Methods

## RESOURCE AVAILABILITY

### Lead Contact

Further information and requests for resources and reagents should be directed to and will be fulfilled by the Lead Contact, Maria G. Castro.

### Materials Availability

Plasmids and mouse cell primary cultures generated in this study are available upon request to the lead contact.

### Data and Code Availability

Bulk RNA-Seq, ChIP-Seq and Single-cell RNA-seq data have been deposited at Gene Expression Omnibus (GEO). Accession numbers are listed in the key resources table. This paper does not report original code.

## EXPERIMENTAL MODEL AND SUBJECT DETAILS

### Sleeping beauty transposase-based DHG model

All animal studies were conducted according to guidelines approved by the Institutional Animal Care and Use Committee at the University of Michigan. All animals were housed in an AAALAC accredited animal facility and were monitored daily. Studies did not discriminate sex, and both male and females were used. A murine model of glioma and the appropriate comparisons/controls were created by the Sleeping Beauty (SB) Transposase system, which is used to integrate genetic lesions into the genomic DNA of brain cells of neonatal mice. Tumors then develop intracranially de novo from neural progenitor cells. The plasmids encoding the different genetic lesions used in this study are: (i) pT2C-LucPGK-SB100X, henceforth referred to as pSB, encoding SB transposase and luciferase, (ii) pT2CAG-NRASV12, henceforth referred to as pNRAS, encoding a constitutively active mutant of NRAS, NRAS-G12V (iii) pT2-shATRX-GFP, henceforth referred to as pshATRX, encoding a short hairpin against ATRX (iv) pT2-shp53, henceforth referred to as pshp53, encoding a short hairpin against Trp53 and (v) pKT-H3.3(G34R)-IRES-Katushka, henceforth referred to as pH3.3-G34R, encoding H3.3-G34R mutation. This plasmid was generated by the directed insertion of a customized H3.3-G34R cDNA (Origene) into the pKT-IRES-Katushka construct via NotI/XhoI restriction sites. The resultant pH3.3-G34R plasmid was confirmed by Sanger sequencing. These specific genetic lesions, encoded in different plasmids, are flanked by inverted repeats/directed repeats (IR/DR) (24,32), that will be recognized by the SB-transposase, enabling transposon genomic integration(24). The SB-encoding plasmid also contains the coding sequence for the Luciferase enzyme flanked by IR/DR, which will get inserted into the host genome as well. This enables the confirmation of plasmid uptake, one day after plasmid injection, and monitor tumor progression by bioluminescence(24).

Neonatal P01 wild-type C57BL/6 mice of both sexes were used to generate H3.3-G34R and H3.3-WT groups. The genotype of SB-generated high-grade gliomas (HGG) involved these combinations: (i) pSB, pNRAS, pshATRX and pshp53 (H3.3-WT group), (ii) pSB, pNRAS, pshATRX, pshp53 and pH3.3-G34R (H3.3-G34R group). Mice were injected according to a previously described protocol(24). Briefly, plasmids were mixed in mass ratios of 1:2:2:2, or 1:2:2:2:2 with in vivo-JetPEI (VWR). After inducing anesthesia to the pups by hypothermia, and placing the head into the stereotactic frame, the lateral ventricle (1.5 mm from lambda, 0.8 mm lateral, and 1.5 mm deep) of neonatal P01 mice was injected with 0.75 μL plasmid mixture (0.5 μL/min) that included: (1) pSB, (2) pNRAS, (3) pshp53, (4) pshATRX, and (5) with or without pH3.3-G34R.

To monitor plasmid uptake in neonatal pups, 30 μL of luciferin (30 mg/mL; GOLDBIO) was injected subcutaneously into each pup 24-48 hours after plasmid injection. *In vivo* bioluminescence was measured on an IVIS® Spectrum imaging system (Perkin Elmer). For the IVIS spectrum, the following settings were used: automatic exposure, large binning, and aperture f = 1. For *in vivo* imaging of tumor formation and progression in adult mice, 100 μL of luciferin solution was injected intra-peritoneally (ip) and mice were then anesthetized with oxygen/1.5-2.5% isoflurane (VETONE). To score luminescence, Living Image Software Version 4.3.1 (Perkin Elmer) was used. A region of interest (ROI) was defined as a circle over the head, and luminescence intensity was measured using the calibrated unit’s photons/s/cm^2^/sr. Multiple images were taken over a 25-minute period following injection, and maximal intensity was reported.

For survival studies, animals were monitored daily for signs of morbidity, including ataxia, impaired mobility, hunched posture, seizures, and scruffy fur. Animals displaying symptoms of morbidity were transcardially perfused, then tissues were fixed and processed for histology.

### CDKN2A^-/-^ Alternative model

The DHG were induced using the Sleeping Beauty (SB) Transposase system as in the previous section. In this alternative model, the genetic lesions were incorporated into neonatal CDKN2A homozygous knocked-down (CDKN2A^-/-^) mice. CDKN2A^-/-^ mice were obtained from the Frederick National Library for Cancer Research as frozen embryos.

Embryos were implanted into receptive adult B6 females by the Transgenic Animal Core of the University of Michigan and the obtained transgenic mice, which were CDKN2A heterozygous, were mated with B6 mice. The CDKN2A KO heterozygous mice of second generation were mated and the progeny obtained was genotyped to confirm the homozygous CDKN2A deletion. From this point, the CDKN2A^-/-^ colony was maintained by mating CDKN2A^-/-^ mice and monitored regularly by CDKN2A genotypification. The genotype of the genetically engineered mice involved these combinations: (i) CDKN2A deletion, p53 knock down, PDGFRα-D842V expression, and shATRX. The tumors were developed using the Sleeping Beauty transposon system as explained in the previous. This model is described in previous a previous study from our group(17).

### Mouse Cell cultures

Primary glioma neurospheres (NS) (H3.3-G34R NS and H3.3-WT NS) were generated from our SB transposase HGG model. Briefly, tumors were harvested at the time of euthanasia by transcardial perfusion with Tyrode’s solution. The brains were excised, and the tumors were identified by GFP or by GFP/Katushka expression under an epi-fluorescence stereo microscopes (Olympus) at the time of resection. The tumor mass was dissociated using non-enzymatic cell dissociation buffer (Gibco), filtered through a 70 μm strainer and maintained in neurosphere cell medium (DMEM/F12 (Gibco) with L-Glutamine (Gibco), B-27 supplement (1X; Gibco), N-2 supplement (1X; Gibco), penicillin-streptomycin (100 U/mL; Gibco), and Normocin (1X; InvivoGen)) at 37 °C, 5% CO_2_. hFGF and hEGF (Peprotech) were added twice weekly at 1 μL (20 ng/μL each stock, 1000x formulation) per 1 mL medium.

### Human DHG cells

The cell culture and generation of human DHG stably transfected cells expressing G34R and cell cultures of DHG patient–derived cells expressing the G34R mutation is described in(17). SJ-GBM2 cells (CVCL_M141) were a gift of the COG Repository at the Health Science Center, Texas Tech University (Lubbock, Texas, USA). SJ-GBM2 cells were grown in IMDM (Gibco, Thermo Fisher Scientific) supplemented with 20% FBS at 37.0°C, 5% CO_2_, and were used in early passages and tested regularly for mycoplasma. SJ-GBM2 transgenic cells (SJ-GBM-2-H3.3-G34R and SJ-GBM-2-H3.3-WT) were maintained with puromycin (10 μg/μL, Goldbio). These cells are referred to as H3.3-G34R and human DHG cells.

All human glioma cells derived from patients were cultured at 37.0°C with 5% CO_2_. Specifically, KNS-42 cells were maintained in DMEM/F12 medium supplemented with 20% FBS. The OPBG-GBM-001, VUMC, and HSJD cells were cultured in a tumor stem medium composed of Neurobasal-A and DMEM/F12 (1:1 ratio), further supplemented with 10 mM HEPES buffer, 1 mM MEM sodium pyruvate, 0.1 mM MEM non-essential amino acids, GlutaMAX-I (1x), B-27 without vitamin A (1x), Antibiotic-Antimycotic (1x), and Normocin (1x). EGF, FGF, PDGF-AA, and PDGF-BB were added twice a week.

### Intracranial glioma mouse NS implantation model

Both male and female C57BL/6 (Taconic) or *Rag-1* KO (B6.129S2-Cd8^atm1Ma^k/J; Jackson Laboratory) or CD8 KO (B6.129S7-Rag1^tm1Mom^; Jackson Laboratory) mice, aged 6-8 weeks old, were used for implantation models. Intracranial tumors were established by stereotactic injection of 5 x 10^4^ H3.3-WT or H3.3-G34R mouse tumor NS into the right striatum using a 22-gauge Hamilton syringe (1 μL per minute) with the following coordinates: +1.00 mm anterior, 2.5 mm lateral, and 3.5 mm deep. The presence of tumors was verified 5 days post-implantation (DPI) by bioluminescence imaging. Usually, five to ten animals were used per treatment.

### Isolation of PBMCs from Human Blood

Heparinized whole blood from healthy human donors was obtained by venipuncture from the Platelet Pharmacology and Physiology Core at the University of Michigan. The PBMCs were isolated as previously described(76). In brief, whole blood was mixed at a 1:1 volume of 3% dextran in 0.9% NaCl and incubated upright for 20 min at RT. The upper, white, leukocyte layer is transferred to another conical tube. This conical tube was centrifuged at 400g for 5 min and resuspened in HBSS. Histopaque-1077 was added gently to underlie the leukocyte suspension. This solution was centrifuged at 400*g* for 20 min at room temperature to separate the human PBMCs (hPBMCs) from neutrophils and remaining erythrocytes. Any residual erythrocytes in the hPBMC pellet were removed using RBC lysis buffer.

## METHOD DETAILS

### Perfusion, fixation and paraffin embedding

After SB-plasmid or NS implantation, animals were monitored daily for signs of morbidity, including ataxia, impaired mobility, hunched posture, seizures, and scruffy fur. Animals displaying symptoms of morbidity were transcardially perfused using Tyrode’s solution (0.8% NaCl (Sigma Aldrich), 0.0264% CaCl_2_2H_2_O (Sigma Aldrich), 0.005% NaH_2_PO_4_ (Sigma Aldrich), 0.1% glucose (Sigma Aldrich), 0.1% NaHCO_3_ (Sigma Aldrich), 0.02%KCl (Sigma Aldrich)), followed by fixation with 4% paraformaldehyde (PFA; VWR) in Dulbecco’s modified phosphate buffered saline (DPBS; Gibco). Mouse brains were then removed from the skull and fixed in 4% PFA for an additional 48 hours at 4 °C. Then, brains were dehydrated by immersing them in solutions of increasing amounts of ethanol and lastly xylene using an automatic processor (HistoCore PELORIS 3 Premium Tissue Processing System, Leica), at the Histology Unit of the Orthopaedic Research Laboratories, at the University of Michigan. Brains were then embedded in paraffin using a Leica ASP 300 paraffin tissue processor/Tissue-Tek paraffin tissue embedding station (Leica) at the Histology Unit of the Orthopaedic Research Laboratories, at the University of Michigan.

### Hematoxylin and eosin and immunohistochemistry

Paraffin-embedded tissue was sectioned using a rotary microtome (Leica) set to 5 μm in the z-direction. Hematoxylin and eosin (H&E) staining was performed as described by us previously(24). For immunohistochemistry (IHC), antigen retrieval slides were placed in a pressure cooker in citrate buffer (10 mM Citric Acid (Sigma Aldrich), 0.05% Tween 20 (Sigma Aldrich), pH 6). After endogenous peroxidase quenching in 0.3% H_2_O_2_ (Sigma Aldrich) and blocking in 10% goat serum in TBST for 2 hours, samples were incubated with primary antibodies (GFAP (Millipore Sigma, AB5541, 1:200), MBP (Millipore Sigma, MAB386, 1:200, CD68 (Abcam, ab125212 1:1000), and Nestin (Novus Biologicals, (NB100-1604), 1:1000)) overnight at RT. The next day, sections were washed 3 times with TBS-Tx and incubated with biotinylated secondary antibody at 1:1000 dilution in 1% goat serum TBST in the dark for 2 h. Biotin-labeled sections were developed using 3,3′-diaminobenzidine (DAB) with nickel sulfate precipitation according to the manufacturer’s instructions. Brain sections were then washed three times, followed by dehydration with xylene, and finally, coverslips were placed on the slides using DePeX Mounting Medium (Electron Microscopy Sciences). Images were obtained using brightfield/epifluorescence (Carl Zeiss MicroImaging) and analyzed using LSM5 software (Carl Zeiss MicroImaging).

### Multiplex immunofluorescence

Multiplex immunofluorescence imaging of human FFPE tissue was performed on a PhenoCycler-Fusion by Akoya Biosciences per manufacturer’s instructions with a photobleaching protocol added to minimize autofluorescence. Akoya antibodies, listed in the materials section, were used, except for a fraction that were custom conjugated to Akoya barcodes. Per Akoya’s protocol, samples are stained with antibodies conjugated to unique oligonucleotide sequences, or barcodes. Reporters, composed of complementary barcodes labeled with fluorophores, are hybridized to their associated antibodies, then imaged and removed, three at a time, in consecutive cycles on the Phenocycler-Fusion platform.

The QPTIFF images generated were imported into QuPath version 0.4.3(77) for sample annotation, cell segmentation, and cell phenotyping. Samples were annotated to include only tumor bulk, which was determined based on cell density and SOX2 staining. Next, areas with artifacts were excluded. Segmentation was performed on the final annotation using the StarDist segmentation script(78). Approximately 1.4 million cells from seven samples were analyzed. After segmentation, phenotyping was performed using a hierarchical scheme implemented with a Groovy script using object classifiers that were trained using artificial neural networks.

After classification of all cells was complete, data was exported into CytoMAP(79) for spatial analysis. Raster scanned neighborhoods with a 50-micrometer radius were defined. Resulting neighborhoods were clustered into 10 regions using the NN self-organizing map algorithm, and similar neighborhoods were combined, resulting in seven final neighborhood types. Cell type enrichment, defined as a fold enrichment of 1.25 or above, of immunosuppressive cell types, including M-MDSC, PMN-MDSC or M2 macrophages, was determined for each neighborhood. Those neighborhoods enriched in any of these cell types were designated as immunosuppressive.

### Isolation of tumor and immune cells

We established SB glioma mouse model harboring H3.3-G34R mutation or H3.3-WT as previously described. Mouse were euthanized at symptomatic stage and the tumor was dissected and smashed through a 70 μm strainer. After red blood cell (RBC) lysis with 1X RBC Lysis Buffer (BioLegend), cells were resuspended in 90 μL of cold BSA-EDTA buffer (0.5% BSA (Sigma Aldrich), 2 mM Ethylenediaminetetraacetic acid (EDTA; Sigma Aldrich) in DPBS pH 7.2). CD45+ cells were labeled with 10 μL of CD45 MicroBeads (Miltenyi Biotec) per 10^7^ total cells and sorted using a MACS column (Miltenyi Biotec) placed in the magnetic field of a suitable MACS Separator (Miltenyi Biotec). Tumor cell-enriched and CD45+ cell-enriched fractions were obtained for downstream applications such as RNA purification. Cell purity was assessed by flow cytometry (38).

### RNA-sequencing

RNA-sequencing (RNA-Seq) was performed in collaboration with the University of Michigan Advanced Genomics core as previously described(28). Briefly, total RNA was isolated from cells using RNeasy Plus Mini Kit (Qiagen), and 100 ng of purified RNA were sent for analysis. RNA quality was assessed using a TapeStation (Agilent Technologies) following the manufacturer’s recommended protocol. RNA sequencing libraries were prepared according to the TruSeq Stranded Total RNA Human/Mouse/Rat (Illumina) and were sequenced on the HiSeq 2000 platform. The Tuxedo Suite software package was used for alignment, differential expression analysis, and post-analysis diagnostics. The volcano plot was produced using an R base script and encompasses all genes identified by our RNA-seq analysis. To identify significantly differentially expressed genes, we set a log2 fold change cut-off of 0.585 and –log10 (FDR) greater than 1.3. The enrichment map was generated using the Cytoscape platform with a set of the p-value cut off of 0.05 and FDR of 0.1.

### ChIP-Sequencing

For each immunoprecipitation (IP) performed, 1×10^6^ NS were collected, washed in HBSS (Gibco), and pelleted in low binding Corning Costar microtubes. A 95 μL aliquot of digestion buffer (50 mM Tris-HCl (Sigma Aldrich), 1mM CaCl_2_ (Sigma Aldrich), 0.2% Triton X-100 (Sigma Aldrich), pH 8.0) supplemented with 1:100 protease inhibitors (Sigma Aldrich) was added per 1×10^6^ NS and immediately pipetted gently up and down to lyse. Chromatin digestion was initiated by the addition of 5 μL of Micrococcal Nuclease (ThermoFisher Scientific) at 3.83 x 10^-3^ U/μL final concentration, for 12 minutes at 37 °C, and stopped by the addition of 10X stopping buffer (110 mM Tris-HCl, 55 mM EDTA (Sigma Aldrich), pH 8). The lysate was then diluted with an equal volume of X2 RIPA buffer (280 mM NaCl (Sigma Aldrich), 1.8% TritonX-100, 0.2% SDS (Sigma Aldrich), 0.2% sodium deoxycholate (Sigma Aldrich), 5 mM EGTA (ThermoFisher Scientific), protease inhibitor diluted 1:100). At this stage, 10% of the lysate volume was withdrawn as input. The remaining lysate was then precleared with 10 μL of Dynabeads (1:1 mixture of Protein A (ThermoFisher Scientific) and protein G (ThermoFisher Scientific)) and washed in RIPA buffer (10 mM Tris-HCl, 1 mM EDTA, 140 mM NaCl, 1% Triton X-100, 0.1% SDS, 0.1% sodium deoxycholate, pH 8). Samples were then incubated overnight at 4 °C with 1 μg of anti-H3K27ac antibody (Diagenode), anti-H3K27Me3 antibody (Diagenode), anti-H3K4Me1 antibody (Diagenode), anti-H3K4Me3 antibody (Diagenode), anti-H3K36Me3 antibody (Diagenode) or anti-total mouse IgG antibody (Santa Cruz Biotech). Antibodies were validated prior to use with Active Motif’s MODified Histone Peptide Array (Active Motif). Immunoprecipitation was performed in the following day by incubating the samples for 3 hours with 10 μL of Dynabeads (1:1 mixture of Protein A and Protein G (ThermoFisher Scientific)) in 1x RIPA, followed by washing steps with 1X RIPA (5 times, supplemented with 1:1000 dilution of protease inhibitor (Sigma Aldrich)), LiCl buffer (1 time, 250 mM LiCl (Sigma Aldrich, 10 mM Tris-HCl, 1 mM EDTA, 0.5% NP-40, 0.5 % sodium deoxycholate, pH 8, supplemented with 1:1000 dilution of protease inhibitor), and finally TE buffer (1 time, protease inhibitor-free (Corning)). Immunoprecipitated chromatin was freed from histones and Dynabeads by incubating the samples 1 hour at 37°C with proteinase K (Qiagen) at a final concentration of 0.50 mg/mL. Chromatin samples were then purified by the Qiagen QIAquick PCR Purification Kit (Qiagen) and eluted with 50 μL of the EB buffer provided in the kit.

For ChIP-seq, libraries were prepared at the Epigenomics Core using the TruSeq ChIP Library Preparation Kit (Illumina). Input and IP DNA (5-10 ng) was blunt-ended and phosphorylated, followed by the addition of a single adenine nucleotide prior to ligation with an adapter duplex with a T overhang. Ligated products were size-selected by agarose gel electrophoresis, purified, and PCR-amplified for the final library preparation. Quality control was performed on each input/IP library generated using loci-specific primers to verify that the original enrichment is retained in the final library. Input/IP libraries that passed this quality control were then sequenced on an Illumina NovaSeq 6000 platform.

For ChIP-seq data analysis, the ChIP-seq raw data quality was assessed using FastQC 0.11.3, and the adapters were trimmed using TrimGalore 0.4.4. ChIP-seq and input reads were aligned to the mouse reference genome (mm10) using bowtie2 (80) with default options, and the unique mapped reads were extracted for peak calling. The post-alignment quality control and visualized were performed by Deeptools2 3.5.1 (81). PePr (82) was used to identify the differential H3K27ac, H3K27Me3, H3K4Me1, H3K4Me3 or H3K36Me3 peaks between H3.3-WT and H3.3-G34R groups using FDR < 0.05 and fold change > 3 as the cutoffs. The H3K4me3 peaks were set as sharp peaks, while the other four marks were set as broad peaks. The identified peaks were annotated to genomic features (1 to 5 kb upstream of promoter, promoter, intron, exon, UTR, CDS, and intergenic) using Bioconductor *annotatr* package (83). As a comparison, the same number of peaks were randomly generated across the same chromosome and annotated in the same way by *annotatr.* Gene set enrichment testing was performed on the identified differential peaks by Bioconductor package *chipenrich* (84) to detect enriched Gene Ontology Biological Process terms.

ChromHMM algorithm was applied to identify the Chromatin States (CS) for each sample by integrating the five chromatin marks (85). The alignment BAM files were binarized into 200-bp window by BinarizeBAM, and the mouse enhancers (86) were included for state annotation. An 18-state model was learned by ChromHMM, and the regions and states enriched in specific combinations of histone modifications were identified using the default setting. These states were manually annotated based on the combination of the different type of chromatin marks and their genomic locations, using the 18-state annotations in the RoadMap project as a reference. After CS assignment, CS changes were analyzed between H3.3-WT and H3.3-G34R NS epigenomes by applying the method developed by Fiziev *et al.*(87). Briefly, the overlapped bins between H3.3-WT and H3.3-G34R were identified, and the number of overlapped bins was counted for each pair of CS change. The percentage of each state was normalized to the expected number of that state in each group, assuming that the states in each group are independent. To control for the state similarity between each transition pair, the relative enrichment of each pair pf CS change was calculated by taking the ratio of normalized state change from H3.3-WT to H3.3-G34R and that from H3.3-G34R to H3.3-WT. Those CS changes with relative enrichment > 2 were considered to be significant.

### Western blot (WB)

Neurospheres were harvested and total protein extracts were prepared in RIPA lysis buffer (ThermoFisher Scientific) with 1x of protease/phosphatase inhibitor cocktail (ThermoFisher Scientific). Twenty-five μg of protein extract (determined by bicinchoninic acid assay (BCA; ThermoFisher Scientific)) were separated by 4-12% SDS-PAGE (Invitrogen) and transferred to nitrocellulose membranes (BioRad). After blocking with 5% non-fat milk in TBS (BioRad) with 0.1% Tween (Sigma Aldrich) (TBST), the membrane was probed with primary antibodies overnight at 4 °C. Then, membranes were washed with TBST and the secondary antibody was added: goat anti-rabbit (Agilent Technologies) 1:4000 or rabbit anti-mouse (Agilent Technologies) 1:4000. Enhanced chemiluminescence reagents (ThermoFisher Scientific) were used to detect the signals following the manufacturer’s instructions using a ChemiDoc XRS+ System (BioRad).

To assess the magnitude of response of tumor cells to IFNγ by pSTAT1/STAT1, pJAK2/JAK2, pSTAT5A/STAT5A quantification, NS were incubated with IFNγ (200UI; Preprotech Inc.) for 0, 0.5, 2, 4, 8 and 24 hours, when total protein extracts were obtained for Western blotting. To assess the levels of MHC-I and CIITA in both human and mouse tumor cells, total protein extracts were obtained for western blotting. Total protein extracts from human THP-1 cells and mouse splenocytes were kept as control. All the western blots presented in the figures were performed using the same protein extracts and quantified via BCA assay. The protein concentrations were normalized to 1.5 µg/µl, aliquoted into 20 µl portions, and stored at −80°C. For each western blot, corresponding to different phospho or total JAK/STAT signaling molecules, either actin or vinculin was used as a loading control, depending on the molecular weight of the target protein. Since the protein extracts used in all the western blots for these experiments were the same for each experimental time point and cell type, we included only one representative actin loading control in the figure.

To assess the levels of specific histone modifications, histone extracts were obtained by Histone Purification Mini Kit (Active Motif) following the manufacturer’s protocol and WB was performed on 15 μg of histone extract using 1:1000 of anti-H3K27ac antibody (Diagenode), 1:1000 of anti-H3K27Me3 antibody (Diagenode), 1:1000 of anti-H3K4Me1 antibody (Diagenode), 1:1000 of anti-H3K4Me3 antibody (Diagenode) and 1:1000 of anti-H3K36Me3 antibody (Diagenode). Anti-total-H3 antibody was used as loading control (1:2000) (Cell Signaling Technologies). WB quantification was performed using ImageJ (National Institutes of Health), and the reported data are from three to four individual biological replicates.

### Site-directed mutagenesis

Q5 Site-Directed Mutagenesis Kit (New England Biolabs) was used to mutate specific nucleotides. NEBaseChanger tool was used to design primers carrying H3.3 WT and PCR was performed using the designed primers and master mix solution of Q5 Hot Start High-Fidelity DNA polymerase. Post-PCR, the amplified product was subjected to treatment with a Kinase-Ligase-Dpn1 (KLD) enzyme mixture for 5 minutes at room temperature. The product was transformed in high-efficiency NEB 5-alpha competent E. coli cells. Finally, positive site-directed mutagenesis was confirmed by sending the constructs for DNA sequencing (Eurofins Genomics).

### Flow cytometry

At symptomatic stage due to tumor burden, mice were euthanized and tumors were dissected from the brain and made into single cell suspensions as described above. Bone marrow was extracted from the femur, by flushing the cells with DMEM (Gibco) 10% fetal bovine serum (FBS; Peak Serum) using a 10 ml syringe and 30G needles. Blood and spleen were subjected to RBC lysis at 25 °C for 10 minutes.

For live/dead staining, cells were resuspended in 1/1000 dilution of LIVE/DEAD™ Fixable Aqua Dead Cell (Invitrogen) in DPBS (Gibco) and incubated for 30 minutes, at 25 °C, protected from light. After one wash with flow buffer (DPBS (Gibco) + 2% FBS (Peak Serum)), non-specific antibody binding was blocked by resuspending the samples in flow buffer and anti-CD16/CD32 antibody (1/200) (Biolegend). After a 15 minutes’ incubation, on ice, protected from light, samples were washed 2X with flow buffer, and stained with the specific antibodies. Antibody staining was carried out for 30 minutes, on ice, protected from light. Flow data was measured by using a BD FACSAria™ II Flow equipment (Certified Genetool, Inc.) and analyzed using FlowJo version 10 (Treestar).

### Spectral Flow cytometry

At symptomatic stage due to tumor burden, mice were euthanized and tumors were dissected from the brain and made into single cell suspensions. The strainer was washed two times with 10 mL complete media to force cells through and to minimize cell loss. Then, tumor-infiltrating immune cells were enriched using with 30% / 70% Percoll (GE Lifesciences) density gradient. Briefly, cells were centrifuged, and the supernatant was discarded. Then, cells were resuspended in 7 mL of complete media in 15 mL Falcon tube, and 3 ml of 90% stock isotonic percoll™ (SIP, GE Healthcare) were added and mixed well by pipetting up and down 3–5 times. To layer the 70% percoll™ under the 30% percoll™ gradient, 1 mL serological pipette was filled with 1 mL of 70% percoll™ and pushed to the bottom of the 15 mL falcon tube, and the 70% percoll™ was slowly released making sure a clear interface is formed between the two gradients. Finally, the solution was spun at 800 × g for 20 min at room temperature, and the tumor-infiltrating immune cells were collected by carefully isolating the white band that formed at the interface between the two gradients. For antibody staining, all the staining was performed at 4°C to minimize cellular metabolic activity and marker expression changes during the staining procedure.

Cells were first stained with viability dye (Alexa Fluor® 780, Fisher Scientific) to label the dead cells, after which cells were blocked with either fluorescence conjugated CD16/32 for 10 min at 4° C. Monocytic and Polymorphonuclear MDSCs were labeled with anti-CD45 (Biolegend), anti-CD11b (Biolegend), anti-Ly6G (Biolegend) and anti-Ly6C (Biolegend) antibodies (M-MDSC phenotype: CD45^+^ CD11b^+^ Ly6C^hi^ Ly6G^+^; PMN-MDSC phenotype: CD45^+^ CD11b^+^ Ly6C^lo^ Ly6G^-^). Mature dendritic cells were labeled with anti-CD45 (Biolegend), anti-CD11c (Biolegend) and anti-MHC II (Biolegend) antibodies (DC phenotype: CD45^+^ CD11c^+^ MHCII^hi^). Macrophages were labeled with anti-CD45 (Biolegend) and anti-F4/80 (Biolegend) antibodies. M1/M2 macrophages were distinguished by labeling samples with anti-MHCII (Biolegend) and anti-CD206 (Biolegend) antibodies (M1 phenotype: CD206^-^ MHCII^+^; M2 phenotype: CD206^+^ MHCII^-^). NK cells were labeled with anti-CD45 (Biolegend), anti-CD3 (Biolegend) and anti-NK1.1 (Biolegend) antibodies (NK phenotype: CD45^+^ CD3^-^ NK1.1^+^). Total T cells were labeled with anti-CD45 (Biolegend) and anti-CD3 (Biolegend) antibodies.

CD8 T cells were labeled with anti-CD45 (Biolegend), anti-CD3Ɛ (Biolegend) and anti-CD8a (Biolegend) antibodies (CD8 T phenotype: CD45^+^ CD3^+^ CD8^+^). CD4 T cells were labeled with anti-CD45 (Biolegend), anti-CD3 (Biolegend) and anti-CD4 (Biolegend) antibodies (CD4 T phenotype: CD45^+^ CD3^+^ CD4^+^). Antibody staining was carried out for 30 minutes, on ice, protected from light. Flow data was measured by using a Cytek Aurora spectral flow cytometer (Cytek Biosciences) and analyzed using FlowJo version 10 (BD).

### Single-cell RNA Sequencing

Nuclei were prepared from frozen tissue (5-25 mg) Chromium Nuclei Isolation Kit (10X Genomics) following the manual. As nuclei capture was 55%–60% less efficient than cell capture, we aimed to capture 10,000 nuclei per sample. The Chromium Single Cell 3′ (10X Genomics, Ve3) protocol was strictly followed to prepare libraries. The 10X libraries were sequenced on the Illumina NovaSeq 6000 Sequencing System. Cell Ranger V7 (10X Genomics) was used with default parameters to demultiplex reads and align sequencing reads to the genome, distinguish cells from background, intronic counts were included, and obtain gene counts per cell. Alignment was performed using the hg38 reference genome build, coupled with the Ensembl transcriptome (v110). For each sample, cells were filtered using the following quality control (QC) metrics: mitochondrial content (indicative of cell damage), number of genes, and number of unique molecular identifiers (UMIs), using the Seurat v5(88). Thresholds for each sample were selected according to the distribution of each metric within the sample, which varies with sequencing coverage and the number of cells captured. UMI counts and mitochondrial content were regressed from normalized gene counts and the residuals z-scored gene-wise. Dimensionality reduction was performed using principal component analysis (PCA) applied to the most variant genes, and PCA was used as input for projection to two dimensions using uniform manifold approximation and projection (UMAP) and clustering using a shared nearest neighbor modularity optimization algorithm. Cell types were classified by reference mapping with Azimuth against GBmap reference dataset(89).

### RNA-seq Analysis of from DHG patient samples

For RNA-seq data from Chen *et al.*(*15*), cleaned reads were aligned to the human reference genome build hg38 and quantified reads uniquely mapped to exonic regions defined by ensGene annotation set from Ensembl Release 110 using STAR v2.7.10b (90). Normalization, variance-stabilized transformation of the data and differential gene expression analysis were performed using DESeq2 v1.44.0(91). Or published normalized expression Z-score data was download from PedCBioportal (https://pedcbioportal.org/). Only samples with specified mutation status on ATRX and P53 were included. The control group consisted of DHG ATRX mutant, P53 mutant, and H3.3-WT, whereas the G34R group consisted of ATRX mutant, P53 mutant, and H3.3-G34R/V. Pathway analysis of differentially regulated genes was performed using GSEA (http://software.broadinstitute.org/gsea/) or GSVA v 1.52.3(92).

### IFNγ-induced cell apoptosis of mouse NS

Five hundred thousand H3.3-G34R or H3.3-WT NS were incubated with mIFNγ (200 UI; Preprotech Inc) or mIFNγ diluent (0.5% BSA in DPBS) as controls for 24 hours. Cell apoptosis of was assessed by anti-Annexin V/DAPI staining. One million cells were incubated with 5 µl of anti-Annexin V Alexa Fluor 680 antibody (ThermoFisher Scientific) for 10 minutes at 25 °C in 1X Annexin V binding buffer. Then, 400 µl of Annexin V Binding Buffer was added to each tube. DAPI was added at the moment of flow cytometry analysis. Tumor cells negative for both, Annexin V and DAPI staining, were identified as live cells. Tumor cells labeled as Annexin V+ and DAPI-were identified as early apoptotic cells. Tumor cells labeled as Annexin V+ and DAPI+ staining was identified as late apoptotic cells. Tumor cells labeled as Annexin V- and DAPI+ were identified as necrotic cells.

### Cytokine analysis

To assess and quantify the cytokines secreted by the H3.3-G34R or H3.3-WT-NS, one million cells were incubated in T25 flasks for 48 hours and then the supernatants were centrifuged for 5 minutes at 500 xg to remove cell debris. Levels of cytokines in the NS conditioned media were measured by ELISA, at the Cancer Center Immunology Core, University of Michigan. Briefly, to a 96-well plate 50 μl of assay diluent was added followed by 50 μl of sample, or standard; plate was incubated at 25 °C for 2 hours. Then, wells were washed five times with wash buffer and incubated for 2 h at 25 °C with 100 μl of conjugate solution. Wells were washed a further five times with wash buffer and incubated for 30 minutes at 25 °C with 100 μl of substrate solution. The reaction was stopped by adding 100 μl of stop solution to each well and the absorbance was acquired at 450 nm (background was subtracted by setting the wavelength at 540 nm).

### In vitro mouse T cell proliferation

For T cell Proliferation Assay, splenocytes from OT-1 mice (C57BL/6-Tg(TcraTcrb)1100Mjb/J; Jackson Laboratory) were labeled with 5-(and 6)-carboxyfluorescein diacetate succinimidyl ester (CFSE) according the manufacturer’s instructions (ThermoFisher Scientific). Splenocytes (5.0 x 10^5^) were stimulated with 100 nM SIINFEKL peptide (MBL International) in the presence of 0, 25%, 50% 75% conditioned media from H3.3-WT or H3.3-G34R NS or 1:1 Gr-1+CD11b+ cells isolated from H3.3-G34R or H3.3-WT TMEs, for 4 days in RPMI 1640 (Gibco), 10% FBS, 2-Mercaptoethanol (BME) 55 μM (Gibco). After incubation, T cells were stained with anti-mouse CD45 (Biolegend), anti-CD3Ɛ (Biolegend) and anti-CD8 (Biolegend) antibodies, and proliferation was assessed by CFSE dye dilution using a flow cytometer. Levels of IFN-γ in the cultures’ supernatants were measured by ELISA at the Cancer Center Immunology Core, University of Michigan.

### Human T cell proliferation in glioma cell conditioned media

For T cell proliferation assay, Human PBMCs were stained with 5uM CFSE for 5 min followed by CFSE quenching with HBSS. Using fresh culture media [RPMI 1640 (Gibco 11875093), 10% FBS (Gibco 10437028), 55uM beta-mercaptoethanol (Gibco 21985023), 100unit/mL Penicillin-Streptomycin (Gibco 15140122)], 100uL of media was prepared per well by adding 30 U/ml rIL-2 (Sigma-Aldrich SRP3085) and 4uL prewashed Dynabeads per well. The cells were then counted and seeded at 2 x 10^5^ cells per well in a 96-well plate. The volume in each well was brought to 100uL of fresh culture media, and 100uL of SJ-GBM-2-H3.3-G34R or SJ-GBM-2-H3.3-WT conditioned media was added to each well to generate the 50% conditioned media conditions. To collect the human glioma conditioned media, SJ-GBM-2-H3.3-G34R and SJ-GBM-2-H3.3-WT cells were seeded for 48 hours at a density of 2 x 10^6^ cells in T25 tissue culture flask. The cells were incubated at 37 degrees C for 4 days, after which they were rinsed and stained with CD45, CD4, CD8 antibodies for flow cytometry analysis of T cell proliferation by CFSE dilution.

### IFNγ induced Caspase3/7 activation and apoptosis assay

Ten thousand H3.3-G34R or H3.3-WT human and mouse DHG cells were incubated with IFNγ (200 IU; Peprotech Inc.) or IFNγ diluent (0.5% BSA in DPBS) as a control in a 96-well plate for 24 and 48 hours. The degree of cellular apoptosis was assessed using the Caspase-Glo® 3/7 Assay Kit following the manufacturer’s instructions with some modifications (Promega, Madison, WI, USA). In brief, cells were seeded in a 96-well plate at a density of 10,000 cells/well. Cells were then treated with IFNγ 12 hours after seeding.

Staurosporine (0.5 µM for 12 hours) and boiling the cells at 65 degrees Celsius for 15 minutes were used as positive and negative (vehicle) controls, respectively. Following 24 and 48 hours of incubation, the cells were centrifuged at 2500× g for 5 minutes, and 100 µL of the resulting supernatant was transferred to a clean, opaque white 96-well plate. Reconstituted Caspase-Glo® 3/7 reagent was then added, and the plates were incubated at room temperature with shaking for 15 minutes. Luminescence readings were recorded using a Tecan SPARK Microplate Reader (Grodig, Tecan Austria GmbH, Austria).

### Neurosphere nucleofection

NS were nucleofected (Lonza) with a plasmid expressing the cytoplasmic ovalbumin (cOVA) protein following the manufacturer’s recommendations. Briefly, 350,000 neurospheres were nucleofected with 2 μg of cOVA-expressing plasmid, in 16.4 μl of Nucleofector solution and 3.6 μl of Supplement, using EN-138 setting and a microcuvette. Selection of nucleofected neurospheres was performed by adding 1 µg/ml Puromycin (Sigma Aldrich). cOVA expression was confirmed by Western blotting and cells were maintained in NS media plus Puromycin for stable cOVA expression.

cOVA plasmid with Puromycin resistance was cloned by digesting the pCL-neo-cOVA plasmid (Addgene) with NdeI and MfeI enzymes and inserting this portion between NdeI and EcoRI sites in the pLVX-Puro vector (Clontech).

### Gene therapy and rechallenge assay experiment

Tumor implantation with 50,000 of NS in the right striatum was done as previously described. Fourteen days after implantation, each mouse received an intratumorally injection of either saline or the combination of the two viruses: AdV 5 × 10^8^ plaque-forming units (pfu) of Ad-Flt3L and 2 × 10^8^ pfu of Ad-TK in 1.5 μL volume, in three locations: at 3.7 mm, 3.5 mm, and 3.3 mm ventral from the dura. Ganciclovir (GCV; Biotang) was dissolved in DPBS and was given in a dose of 25 mg/kg i.p, twice a day for 7 days starting immediately the day after viral injection. Animals were monitored daily and euthanized at symptomatic stage due to tumor burden, or they were euthanized by the end of the 7th day, for tissue isolation and flow cytometry analysis. Long-term survivors from the Ad-Flt3L+Ad-TK gene therapy treatment group were rechallenged in the contralateral hemisphere. Fifty-thousand NS were implanted in gene therapy treated animals and in naïve animals as controls. No further treatment was performed in these animals and survival was documented.

### Complete Blood Cell Counts and Serum Biochemistry

Whole blood from mice was collected into EDTA anticoagulant tubes (BD Biosciences) and complete blood counts (CBCs) were run on a Heska Element HT5 (Heska Corporation, Loveland, CO) automated veterinary hematology analyzer. Or whole blood was collected into serum separator tubes (ThermoFisher Scientific), allowed to clot, and separated into serum by centrifugation. Serum chemistries were run on an AU480 chemistry analyzer (Beckman Coulter Inc, Brea, CA). Assays were performed within the ULAM Pathology Core laboratory at the University of Michigan.

### Quantitative real-time PCR (qPCR) analysis

The cultured cells were collected, and RNA was extracted using RNeasy Plus Mini Kit (Qiagen) following the manual. cDNA was synthesized using AffinityScript QPCR cDNA Synthesis Kit (Agilent). qPCR was performed on a ViiA 7 Real-Time PCR system (Applied Biosystems) using SYBR Green qPCR Master Mix Fast SYBR Green Master Mix (Applied Biosystems).

### Posttranslational modifications Phospho-Array

To assess the levels of posttranslational activation of JAK/STAT signaling proteins, we used an antibody array (Jak/Stat Phospho Antibody Array, Cat. # PJS042 FullMoon BioSystems) encompassing 42 site-specific and phospho-specific antibodies involved in the JAK/STAT signaling. Protein was extracted from 2×10^6^ cells (Mouse H3.3-WT and H3.3G34R DHG cells, respectively, treated with mIFNγ (200 UI; Preprotech Inc.) or mIFNγ diluent (0.5% BSA in DPBS), following the protocol from Antibody Array Assay Kit (FullMoon BioSystems, Cat. # KAS02). After purification, proteins were quantified by BCA assay (Assay Kit Pierce BCA, Cat # PI23227, Thermo Scientific) according to the manufacturer instructions, and 100 µg of protein per cell type were biotinylated according to the protocol from Antibody Array Assay Kit (FullMoon BioSystems, Cat. # KAS02). Antibody array slides were blocked, and biotinylated proteins were coupled to the arrays following (Cell Cycle Control Phospho Antibody Array, Cat # PCC238 FullMoon BioSystems) instructions. After protein coupling and washing, the biotinylated proteins that bound the antibodies spots on the array were stained with Cy3-streptavidin according to the manufacturer instructions, and the arrays were developed with a Genepix 4100A Microarray Scanner (UMICH Single Cell Analysis Resource Core).

Images were analyzed using ImageJ software, and spots were quantified into intensity values using the Plugin Protein Array Analyzer for ImageJ (93). Each value was normalized by dividing by the Actin average intensity, and each phospho-specific mark was normalized to each corresponding total antibody average intensity. As the array provides six spots for each site-specific and phospho-specific antibody, the analysis resulted in six normalized intensity values per protein-specific and phospho-specific mark. The values were analyzed in GraphPad Prism, and Unpaired T-tests were performed to compare individual protein and phospo-site intensity between H3.3-G34R and H3.3-WT (mIFNγ-treated and basal control), providing Significance Statistical analyses (False discovery rates) and Fold changes for each protein and phospo-site mark between H3.3-G34R and H3.3-WT DHG mouse cells. Fold change values were depicted in the JAK/STAT pathway (Wikipathways Type II interferon signaling (IFNG) (WP1253).

## QUANTIFICATION AND STATISTICAL ANALYSIS

### Statistics

All statistical analysis were performed using the Prism (GraphPad) software package. Data is represented as mean ± standard deviation (SD) or standard error of the mean (SEM). Statistical significance between 2 groups was determined by a student’s t-test. Statistical significance between groups of 3 or more was determined by a one-way or two-way ANOVA, followed by the Tukey’s multiple comparison test or Sidak’s multiple comparison test. Overall survival data including in animal models were plotted as Kaplan-Meier curves with Mantel-Cox test. For all tests, data were considered significant if p-values were below 0.05 (95% confidence intervals). The sample size (n) along with the statistical test performed and corresponding p-values are indicated in each figure legend. P-values are shown as follows: * p < 0.05, ** p < 0.01, *** p < 0.001, and **** p < 0.0001.

## Supplemental information

Figures S1–S25 and Table S1-S5

Table S1 RNA-seq Tumor cells H3.3-G34R vs H3.3-WT, related to Figure 1

Table S2 ChIP-Seq Differential Peaks results, related to Figure 2

Table S3 JAK-STAT PTM array results, related to Figure 3

Table S4 RNA-seq CD45+ cells, related to Figure 4

Table S5 MHC-1 primer list

---

**Figure S1.**
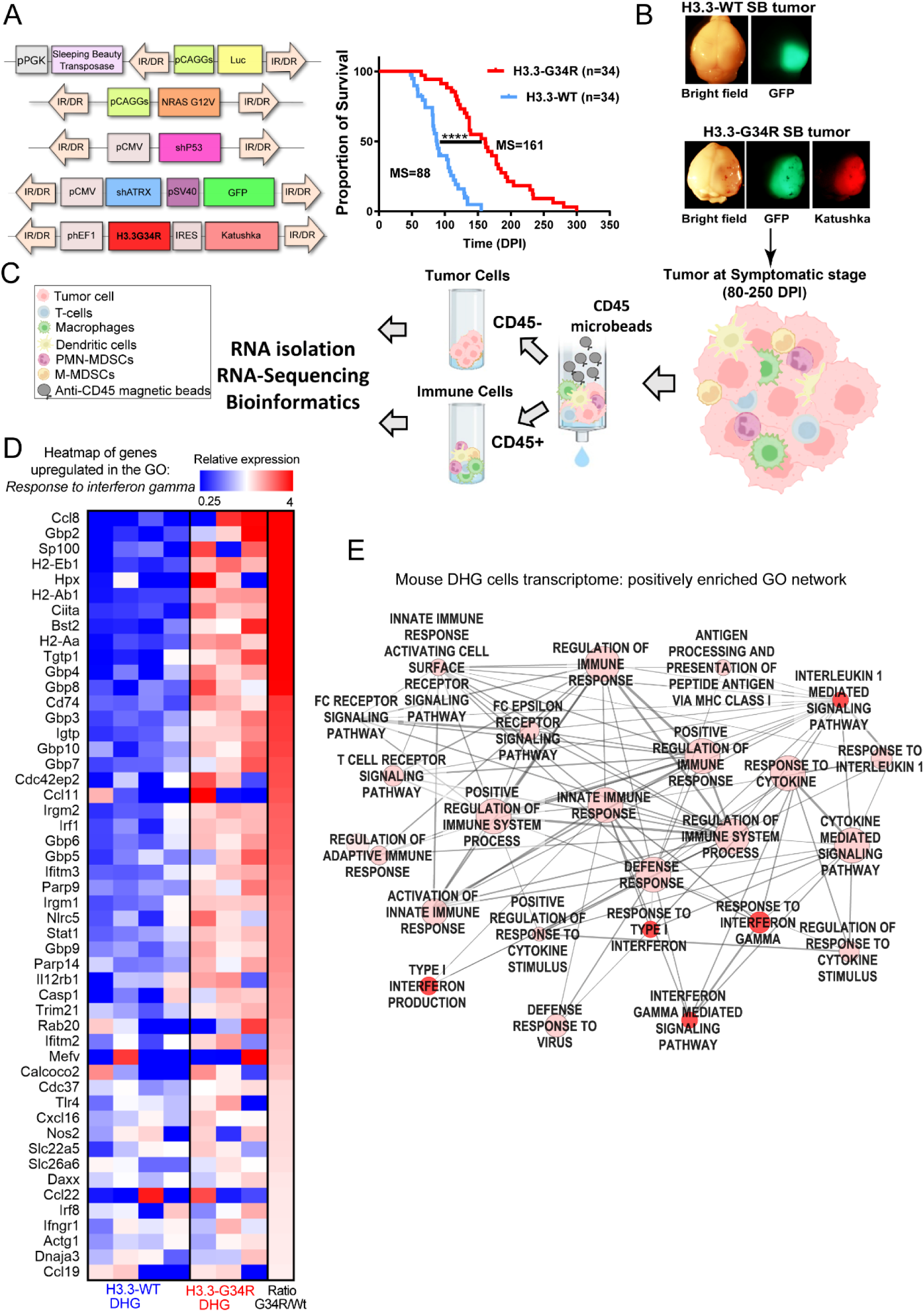
Development of a H3.3-G34R mutant DHG mouse model, related to Figure 1. **(A)** Diagrams of the plasmids used to generate H3.3-WT and H3.3-G34R high grade gliomas. H3.3-WT gliomas were induced by using plasmids encoding receptor tyrosine kinase (RTK)/RAS/PI3K pathway activation (pNRASG12V) in combination with p53 and Atrx downregulation (pshp53 and pshATRX plasmids). To study the impact of H3.3-G34R mutation on tumor’s biology, a plasmid encoding for this mutated histone was added to the same plasmid combination to generate H3.3-G34R high grade gliomas. IR/DR: inverted repeats/directed repeats. Kaplan-Meier survival curve for SB-derived H3.3-WT (n=34) or H3.3-G34R (n=34) tumor bearing animals. **** p<0.0001, Mantel-Cox test. **(B)** Dissecting microscope images showing the fluorescent markers in H3.3-WT DHG (green) and in H3.3-G34R DHG (green and red fluorescence) in sleeping beauty-derived GEMM DHG. **(C)** Scheme illustrating the procedure of sample extraction and analysis of sleeping beauty derived GEMM DHG. Tumors were subjected to dissection assisted by the fluorescence markers and immune cells and tumor cells were separated based on the expression of CD45. Tumor and immune cells were separately subjected to RNA-Seq. **(D)** Heatmap of genes belonging to the GO “*Response to interferon gamma*” upregulated in mouse H3.3-G34R versus H3.3-WT DHG tumor cells. **(E)** Network of selected gene ontologies (GO) upregulated in pathways in mouse H3.3-G34R versus H3.3-WT DHG tumor cells.

**Figure S2.**
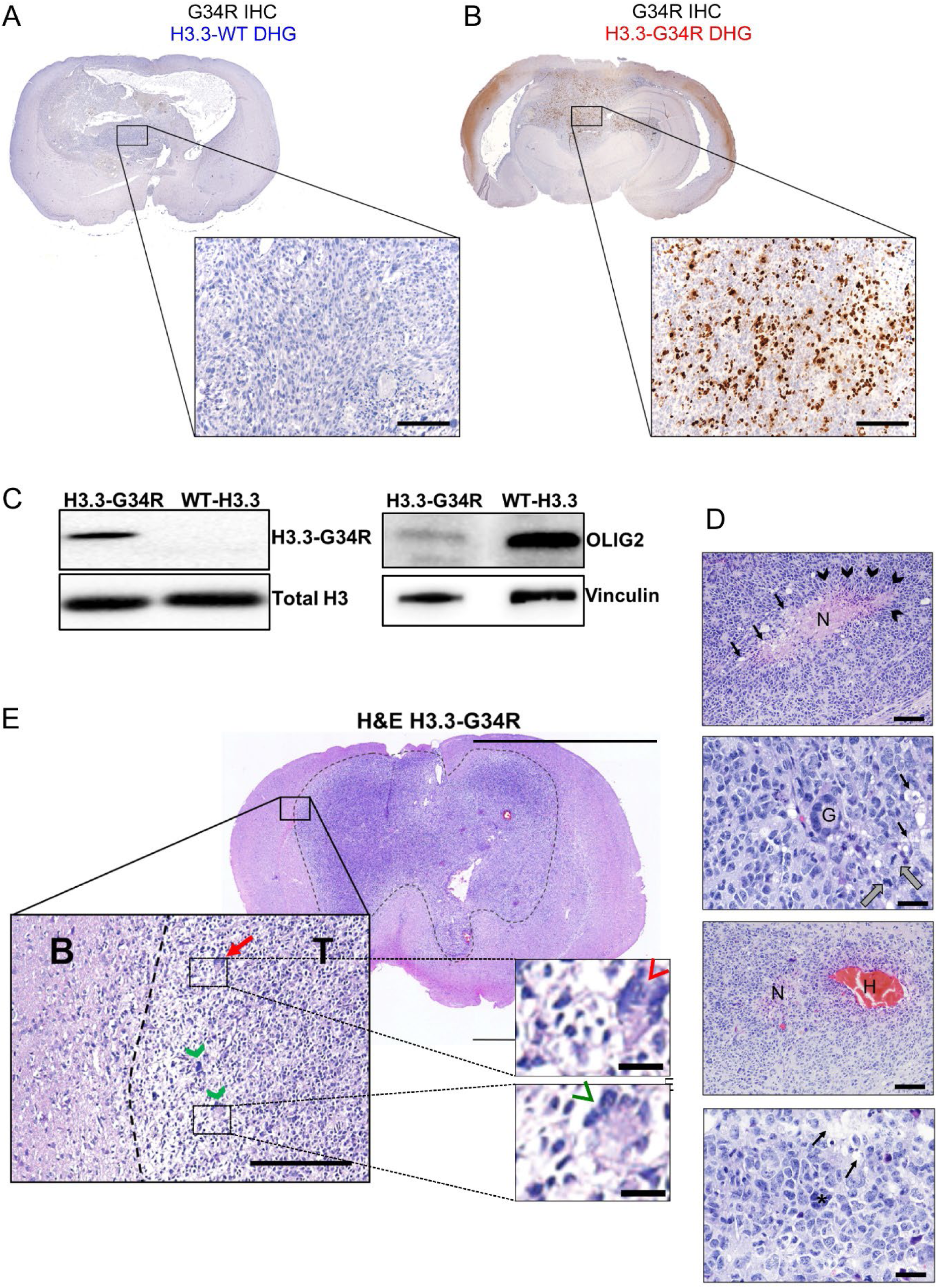
Characterization of a H3.3-G34R mutant DHG mouse model, related to Figure 1. **(A-B)** Immunohistochemistry on SB-derived DHG brain sections. Symptomatic mice bearing SB-H3.3-WT or H3.3–G34R DHG were euthanized, and their brains were processed for histological analysis. Paraffin-embedded brains were cut in 5 µm thick sections and the presence of H3.3-G34R was analyzed by immunohistochemistry (IHC) in H3.3-WT and H3.3-G34R DHG. Scale bars: 500 µm. **(C)** Western blot (WB) analysis of H3.3-G34R expression using total histone extraction from H3.3-WT and H3.3-G34R NS. Total Histone 3 was used as loading control and WB analysis of OLIG2 expression in H3.3-WT and H3.3-G34R NS. Vinculin was used as loading control. **(D)** Histological characteristics of the G34R-mutant DHG genetically engineered mouse model (GEMM). Hematoxylin and eosin (H&E) staining of H3.3-G34R SB-high grade glioma. The dashed line indicates tumor border. Coronal section scale bar = 0.5 cm. The insert image shows a magnified section of the tumor border (dashed line) and histological features of HGG such as hypercellularity, nuclear atypia or multinucleated cells (green arrow heads) and mitosis (red arrow) (Inlets scale bars = 10 μm). T: tumor, B: normal brain. Scale bar = 100 μm. **(E)** H3.3-G34R SB-derived tumors stained with H&E staining displayed common HGG histological features. Scale bars: 100 µm (first and third image) and 50 µm (second and fourth image); N: necrosis; arrowheads: pseudo-palisades; black arrows: microvascular proliferation; H: hemorrhage; G: giant cell; grey arrows: mitotic cells; black arrows: microvascular proliferation.

**Figure S3.**
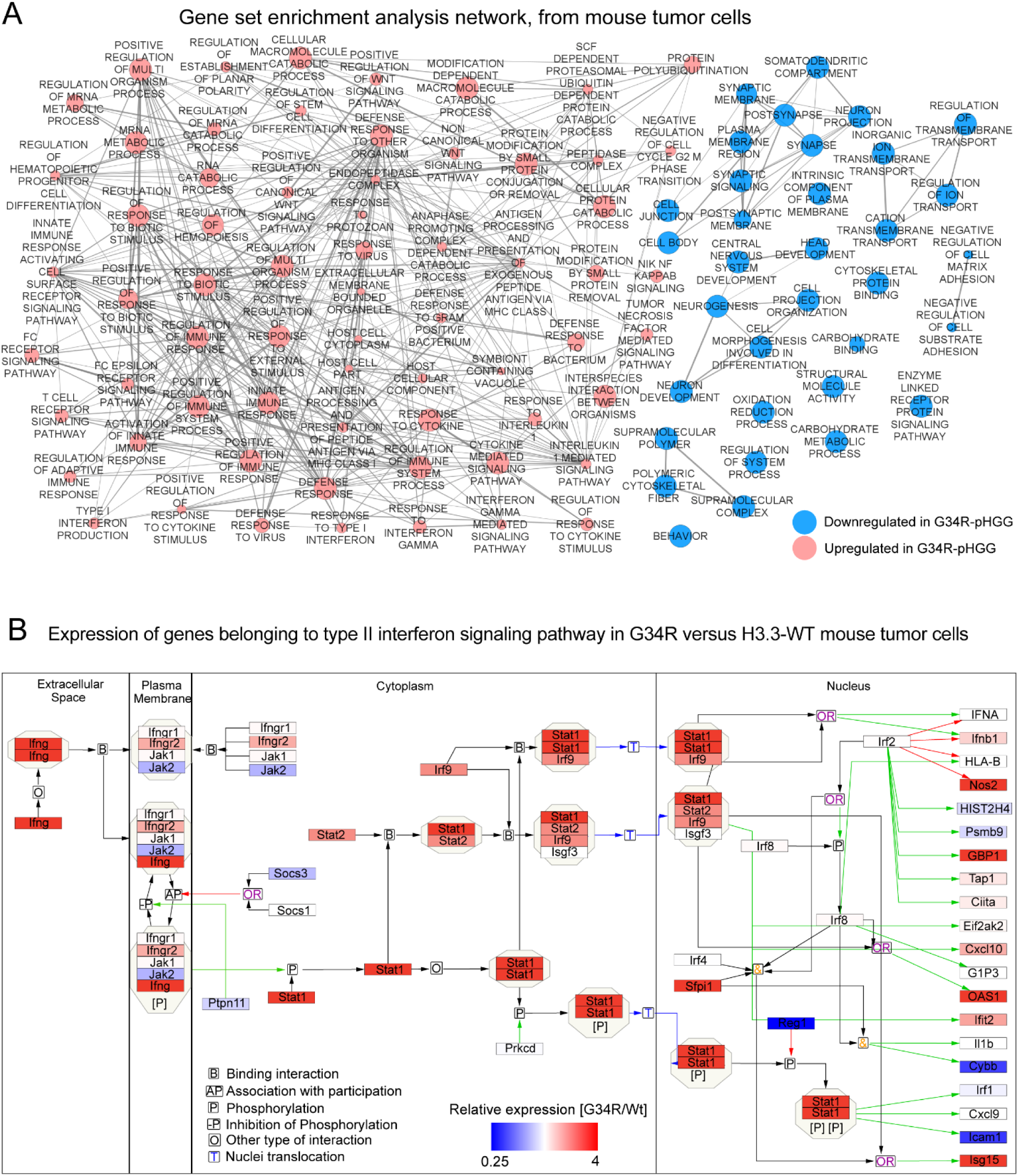
Transcriptome analysis of mouse DHG cells, related to Figure 1. **(A)** Complete Gene ontology (GO) enrichment network resulting from the GSEA of differentially expressed genes in H3.3-G34R versus H3.3-WT tumor cells. Nodes in blue and red illustrate downregulated or upregulated GOs, respectively (p<0.05), FDR<0.1. **(B)** Analysis of the transcriptional regulation of the JAK/STAT pathway in G34R-mutant versus H3.3-WT mouse DHG tumor cells. The relative expression is depicted with a color code, and the pathway was adapted from wikipathways Type II interferon signaling (IFNG) (WP1253).

**Figure S4.**
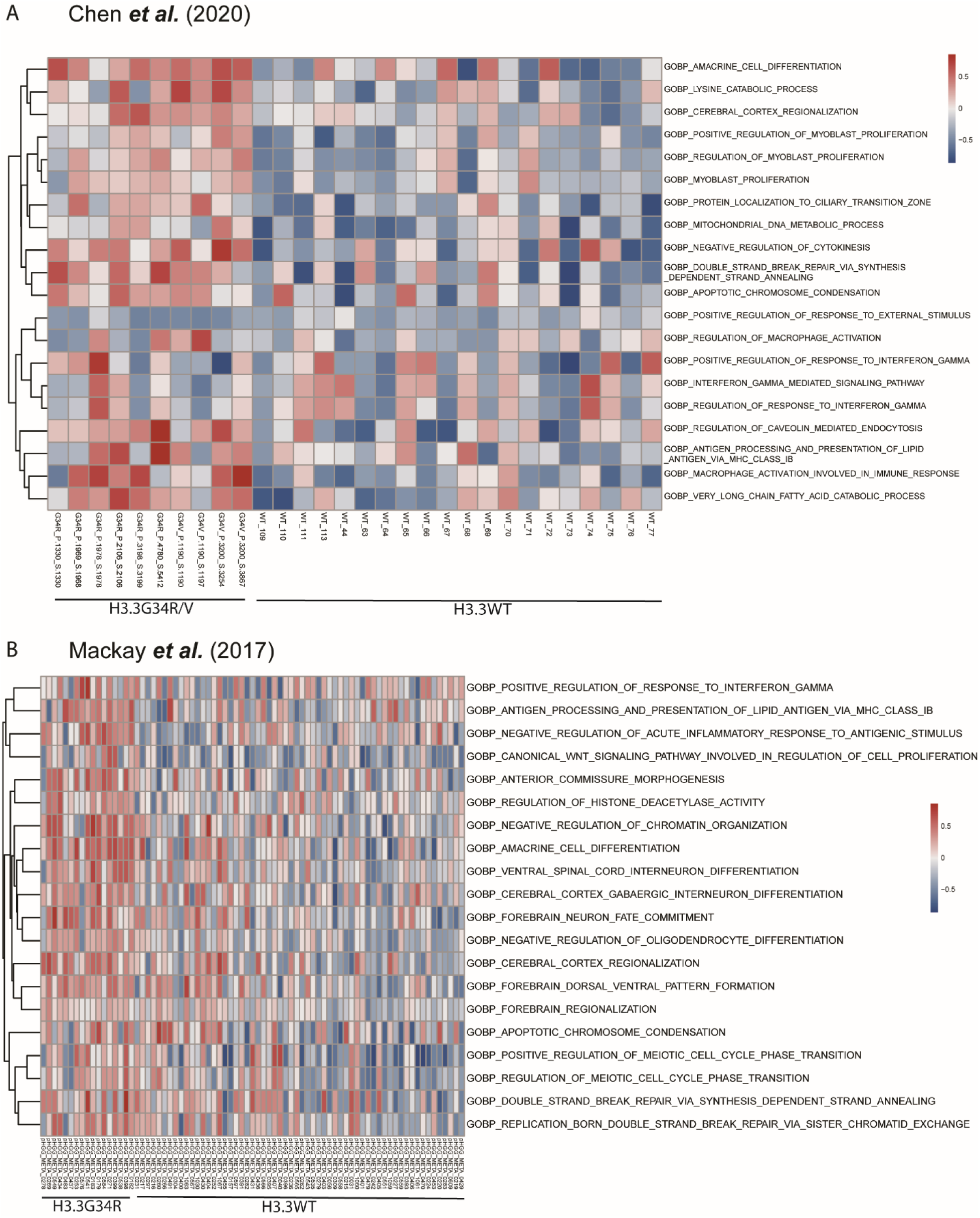
Bulk RNA-Sequencing analysis of DHG patient samples, related to Figure 1. **(A)** Heatmap of Gene set variation analysis (GSVA) scores in G34R/V tumors compared to WT DHG patient tumors from Chen *et al*. **(B)** Heatmap of Gene set variation analysis (GSVA) scores in G34R/V tumors compared to WT DHG patient tumors from Mackay *et al*.

**Figure S5.**
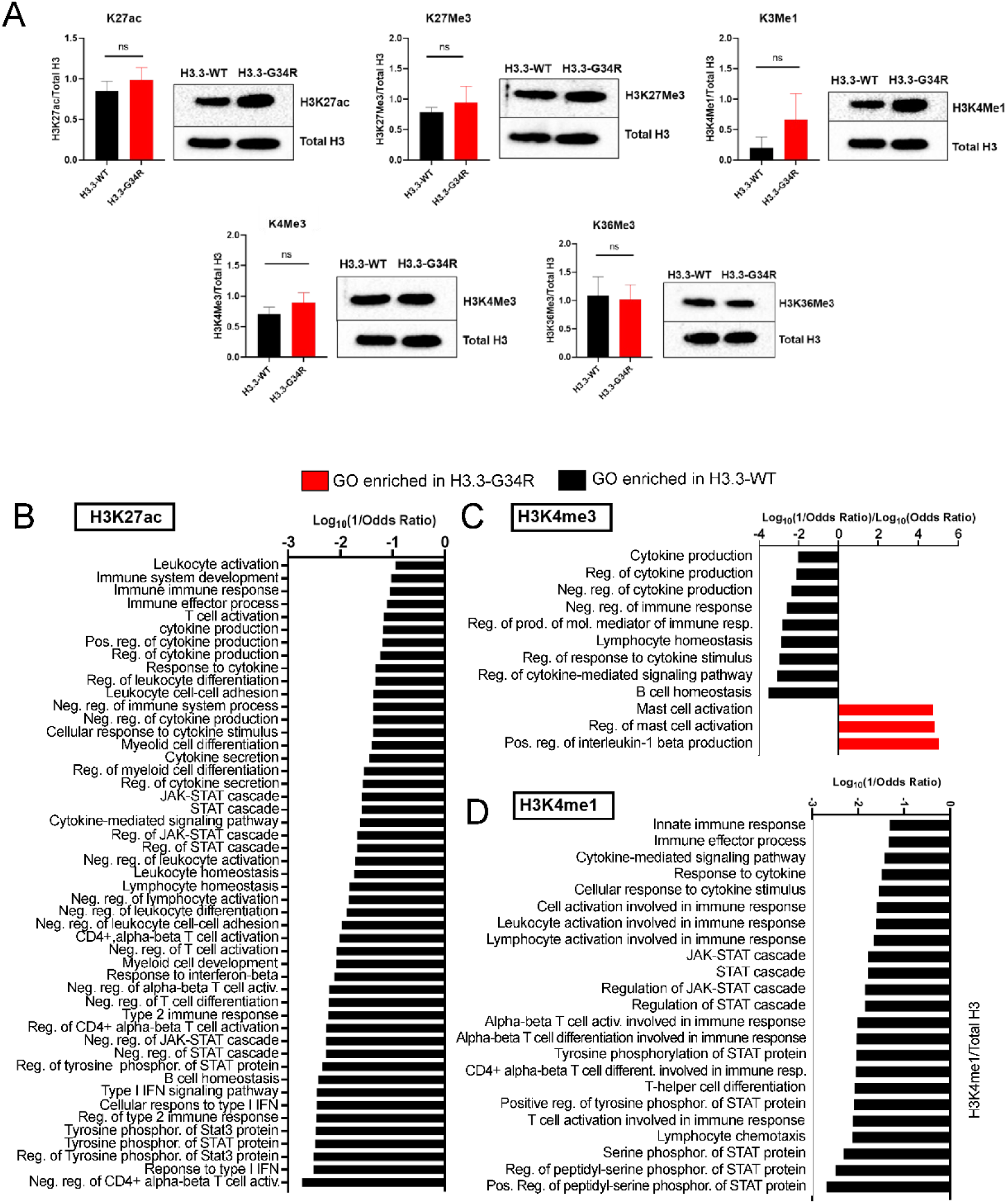
Analysis of histone mark deposition in H3.3-WT and G34R cells, related to Figure 2. **(A)** Western blot analysis of total histone marks in H3.3-WT and H3.3-G34R NS. Total Histone 3 was used as loading control. Representative images of three biological replicates and two independent experiments are displayed. Histograms show the semi-quantification of Western blots. Data shown as means ± SEM, ns= non-significant, Student T test. **(B-D)** Analysis of the GO terms related to immune response resulting from the GSEA of genes enriched in (B) H3K27ac, (C) H3K4me3 and (D) H3K4me1 marks. Red bars represent the Log10 of the Odds Ratio for immune-related-GO-terms enriched in each histone mark in H3.3-G34R NS (positive values), whereas the black bars represent the Log10 of 1/Odds Ratio for immune-related-GO-terms enriched in each histone mark in H3.3-WT NS (negative values).

**Figure S6.**
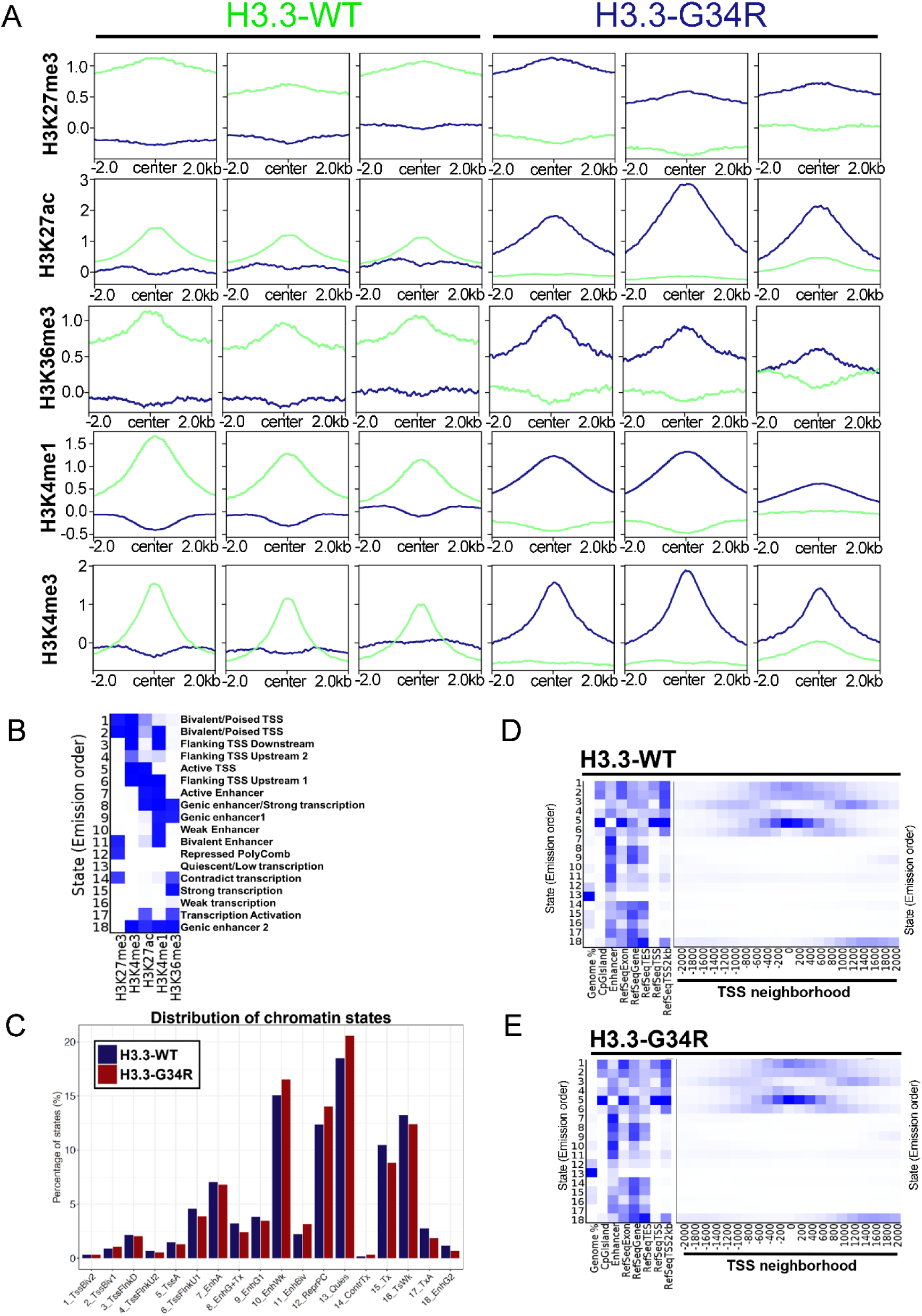
H3.3-G34R affects the binding of different histone marks, impacting on the chromatin state, related to Figure 2. **(A)** Plots show the average histone mark deposition in H3.3-WT (green) and G34R (blue) neurospheres. Differential peak calling was determined using PePr (v1.1.14), p-value < 1e^-5^, n=3. **(B)** Chromatin states were manually labeled based on bibliographic information and they were predicted based on H3.3-WT or H3.3-G34R ChIP-Seq data using ChromHMM. TSS: transcriptional start site. **(C)** Chromatin state distribution is shown as the % of a particular chromatin state (y axis) vs the different chromatin states (x axis) in H3.3-WT and H3.3-G34R groups. Chromatin states are defined as indicated in B). **(D-E)** Abundancy of different chromatin states around the TSS in (D) H3.3-WT NS or (E) H3.3-G34R NS. In both samples, the TSS region was enriched in chromatin states with terms related to transcriptional initiation.

**Figure S7.**
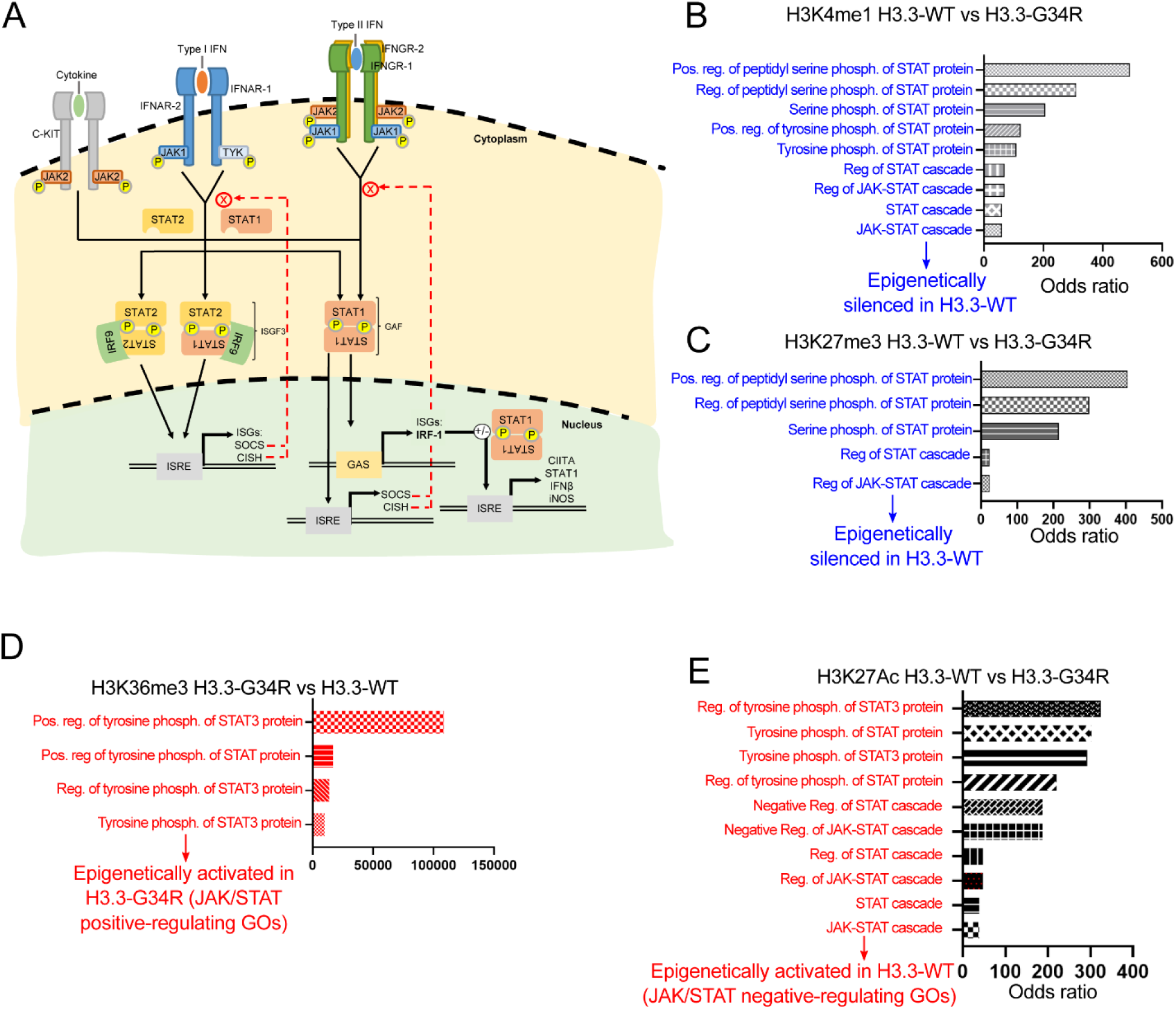
H3.3-G34R affects the epigenetic regulation of STAT pathway-related genes, related to Figure 2. **(A)** Diagram depicting STAT pathways related to Type I and Type II IFNs signaling. GAS: interferon-gamma activated sites; ISRE: interferon-stimulated response element; ISG: interferon-stimulated gene; ISGF: interferon-stimulated gene factor; GAF: gamma-activated factor; *CISH*: Cytokine Inducible SH2 Containing Protein; IRF: interferon regulatory factor. To perform the ChIP-Seq analysis, three biological replicates of H3.3-WT and H3.3-G34R neurospheres were used and five marks were analyzed: H3K27ac, H3K27me3, H3K4me1, H3K4me3 and H3K36me3. **(B-E)** Gene ontologies related to the STAT pathway were enriched in different histone marks. Bar graphs represent the enrichment score (Odds Ratio) for each GO. Those GO terms enriched in H3.3-WT vs H3.3-G34R neurospheres are shown in black (B-D), whereas GO terms enriched in H3.3-G34R vs H3.3-WT are shown in red (E). The GO terms’ significance was determined by FDR<0.05.

**Figure S8.**
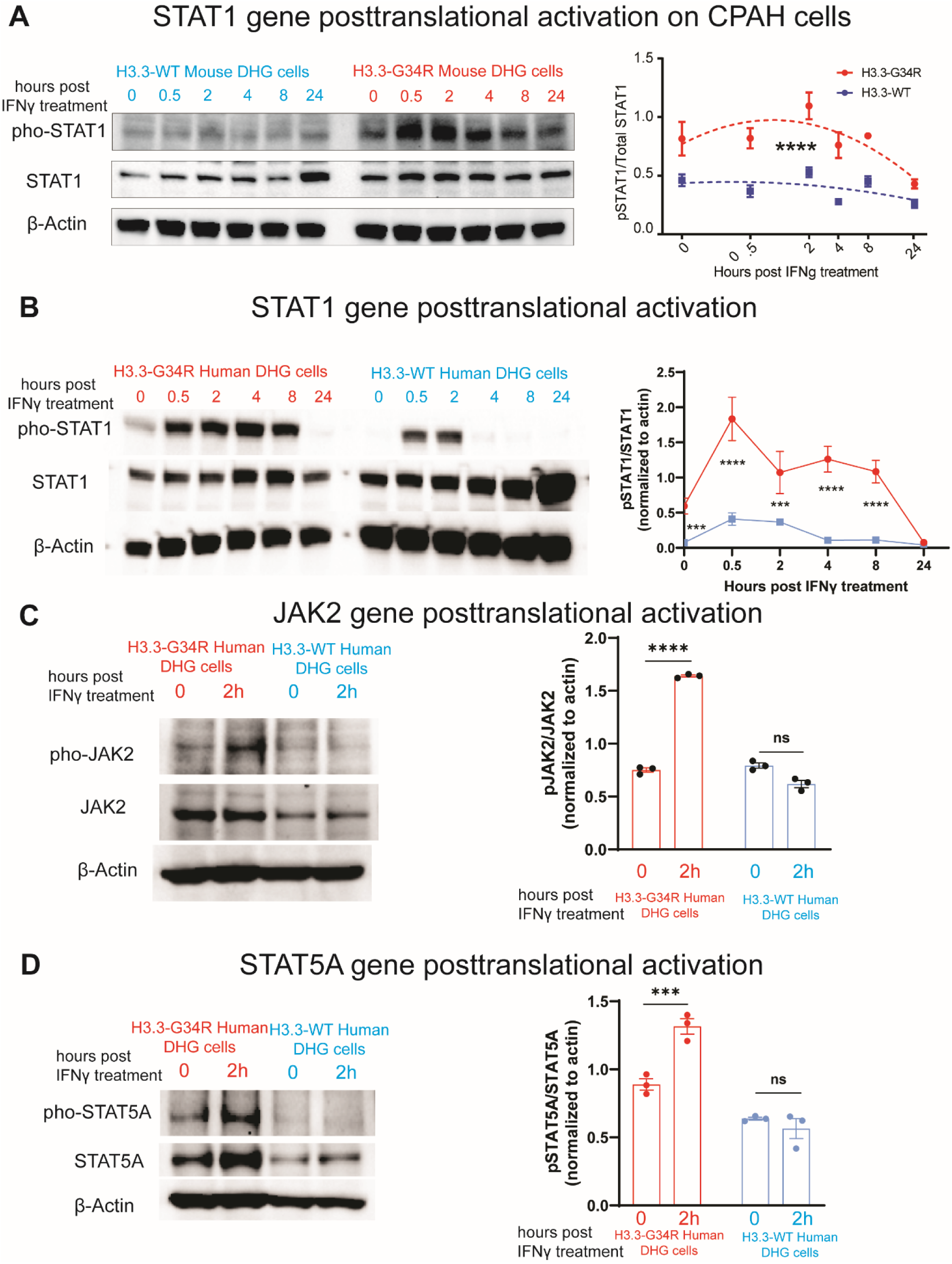
Post-translational activation of the JAK/STAT pathway in H3.3-G34R DHG cells following IFNγ treatment, related to Figure 3. **(A)** Western blot of phospho-STAT1 and STAT1 at different time points after stimulation with IFN-γ, in H3.3-WT and H3.3-G34R CDKN2A KO, P53 KD and ATRX KD mouse DHG cells and quantification of phospho-STAT1 normalized levels. ****P < 0.001, analysis of curve difference from the curve regression model. **(B-D)** Analysis of the JAK/STAT pathway posttranslational activation in H3.3-G34R versus H3.3-WT human DHG cells under basal condition and upon IFNγ (200 U/ml) treatment. **(B)** Western blot of phospho-STAT1 and STAT1 at different time points after stimulation with IFNγ, in H3.3-WT and H3.3-G34R human DHG cells (i.e., SJ-GBM2 H3.3-WT and SJ-GBM2 H3.3-G34R DHG cells). **(C)** Western blot of phospho-JAK2 and JAK2 at basal levels and upon stimulation with IFNγ for 2h in H3.3-WT and H3.3-G34R human DHG cells. **(D)** Western blot of phospho-STAT5A and STAT5A at basal levels and upon stimulation with IFNγ for 2h in H3.3-WT and H3.3-G34R human DHG cells.

**Figure S9.**
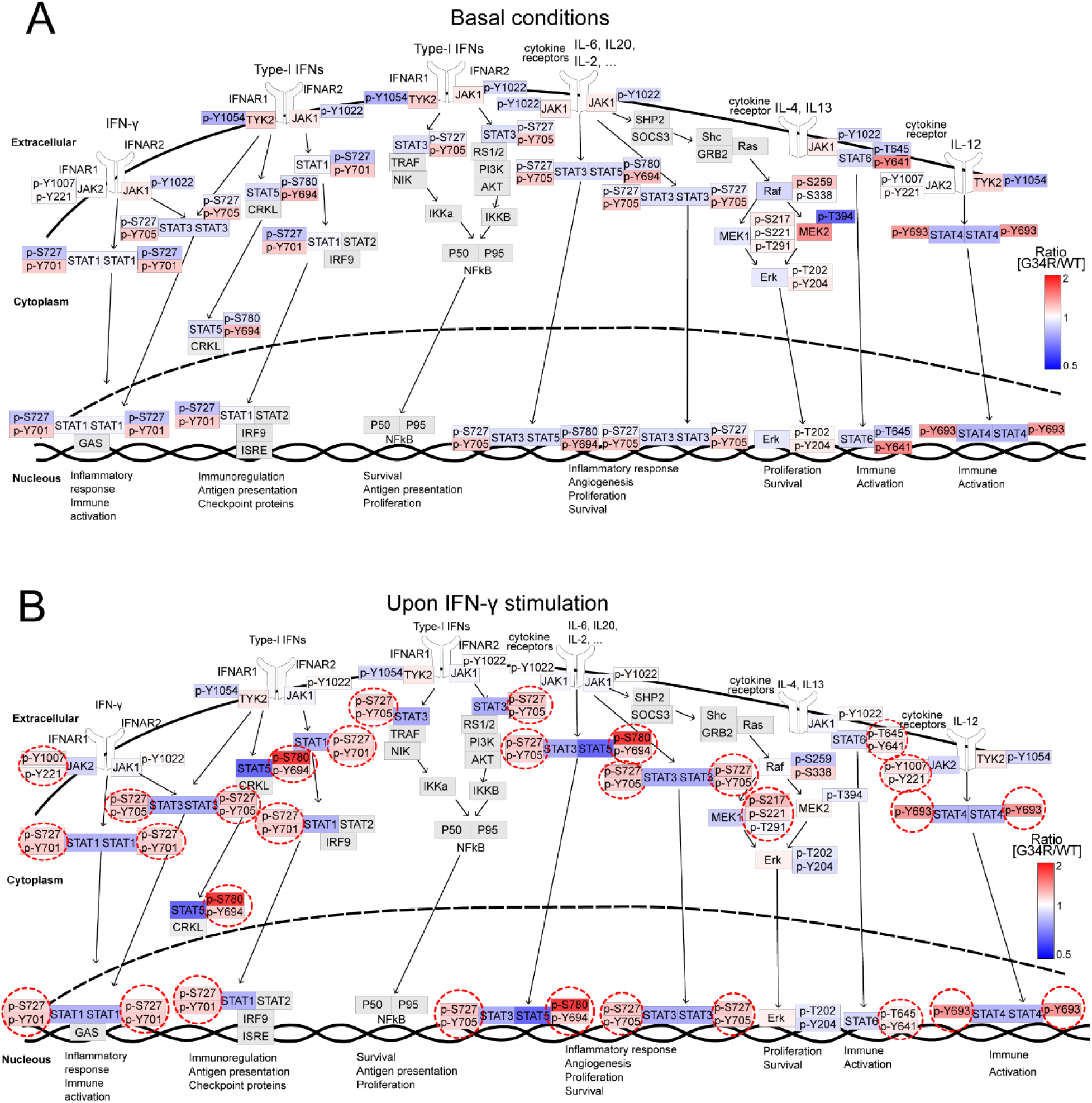
Analysis of the JAK/STAT pathway posttranslational activation in H3.3-G34R versus H3.3-WT DHG mouse cells in basal conditions and upon treatment with IFN-γ, related to Figure 3. **(A-B)** Scheme depicting the JAK/STAT pathway, and the results of the array in each protein and PTM in a color code defined by relative expression (Fold change) between G34R versus WT DHG cells (Blue: downregulated, red: upregulated), in basal conditions (A) and after stimulation with IFN-γ (B).

**Figure S10.**
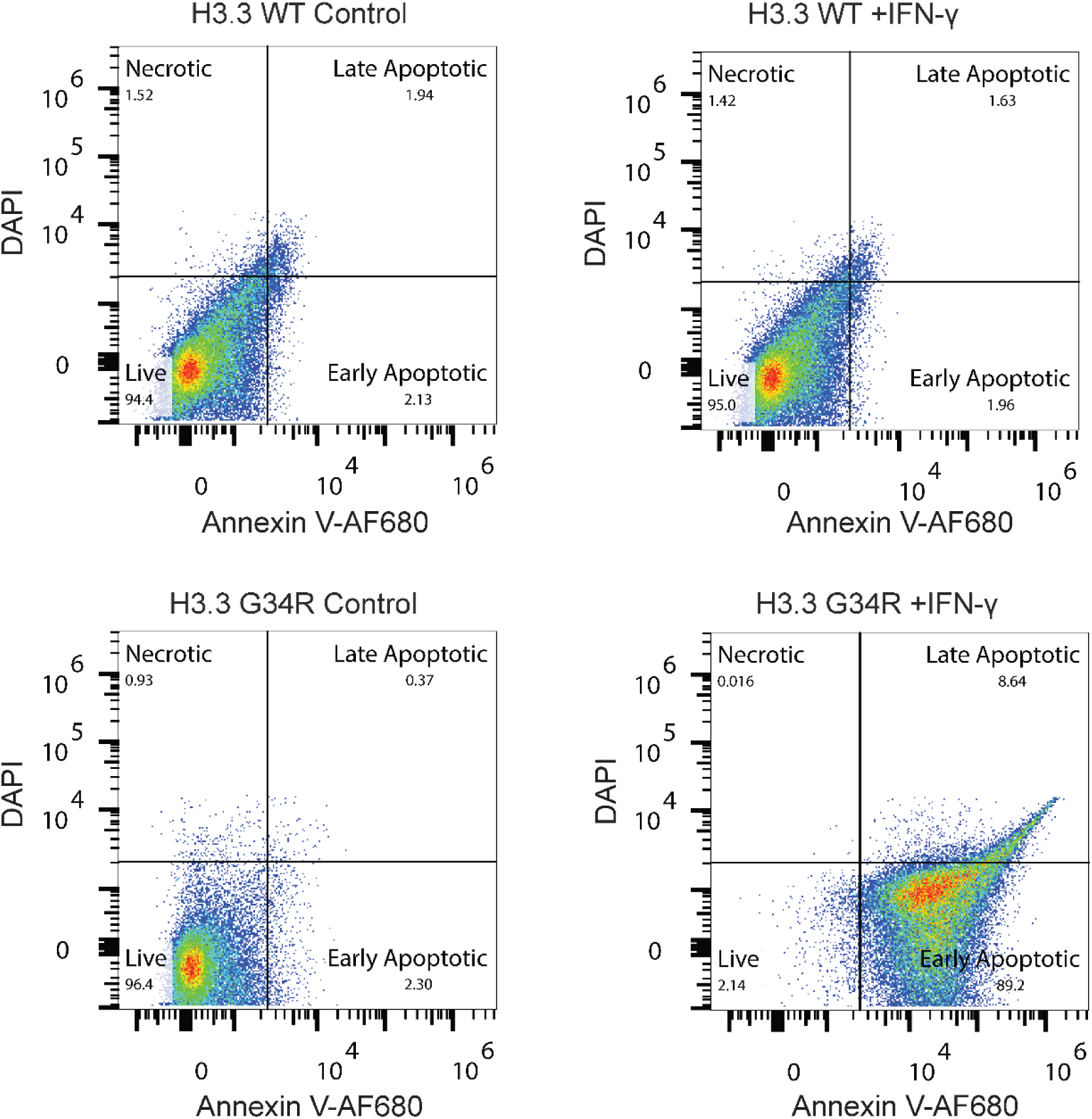
IFN-γ induced apoptosis in H3.3-G34R versus H3.3-WT DHG mouse cells, related to Figure 3. H3.3-G34R or H3.3–WT neurospheres were incubated with IFNγ (200 U) or diluent (control) and apoptosis was measured at 24 hours by Annexin V and DAPI staining.

**Figure S11.**
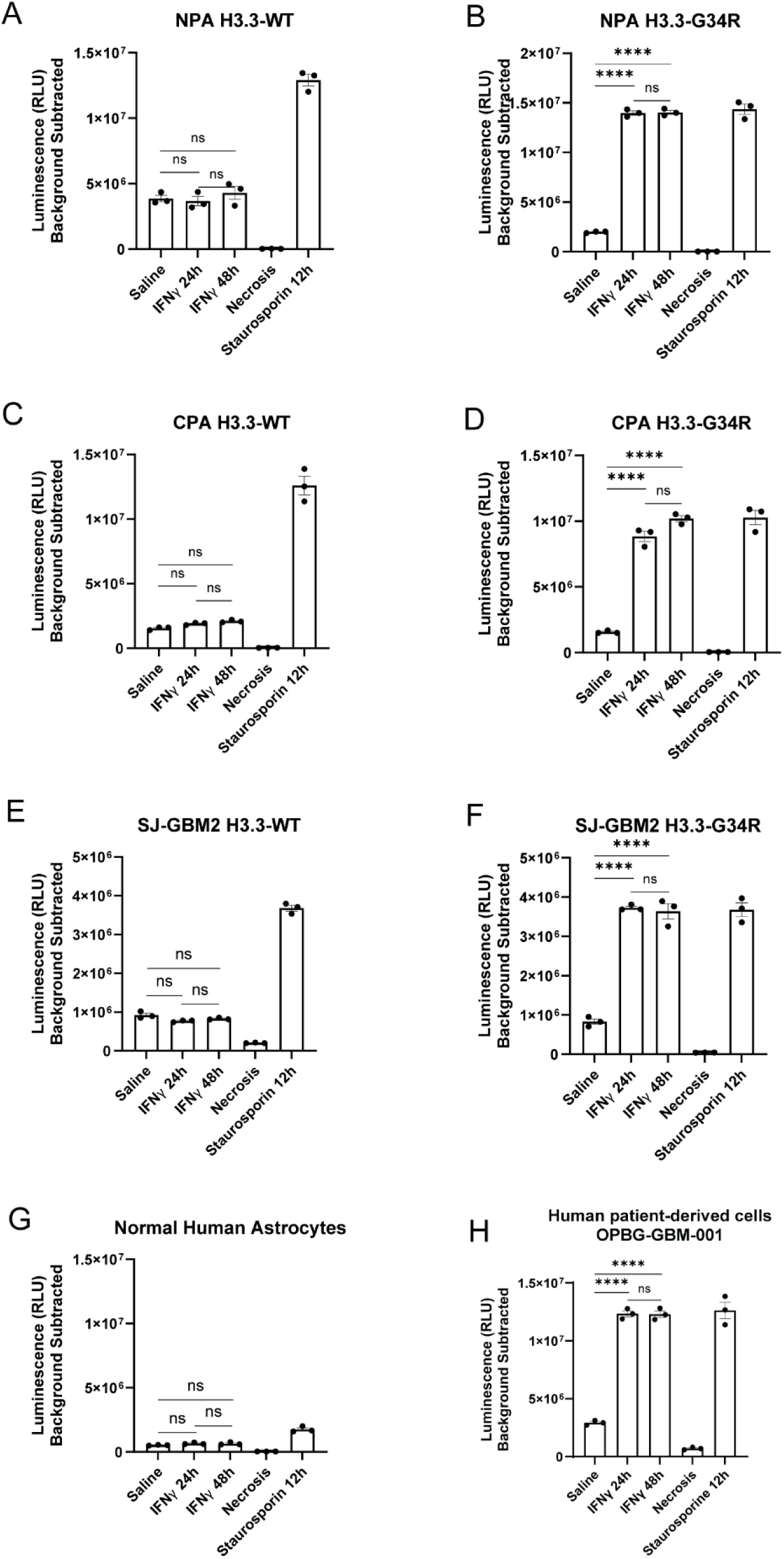
Effect of IFNγ stimulation on apoptosis in human and mouse DHG cells, related to Figure 3. **(A-D)** H3.3-G34R (**B & D**) and H3.3-WT (**A & C**) mouse DHG cells were treated with IFNγ (200 IU) or diluent (control) for 24 and 48 hours to evaluate the level of apoptosis or left untreated. (**E-H.** Similarly, H3.3-G34R human DHG cells (**F**), patient-derived endogenous H3.3-G34R cells (**H**), H3.3-WT human DHG cells (**E**) and normal human astrocytes (**G)** were treated with IFNγ (200IU) or diluent (control) for 24 and 48 hours to evaluate the level of apoptosis after IFNγ treatment. Cells were either treated with staurosporine (0.5µM) for 12 hours as positive control or boiled at 65^0^C for 15 minutes as a necrosis control. Data are presented as relative luminescence unit (RLU). ****P < 0.0001 and ns: not significant unpaired t-test.

**Figure S12.**
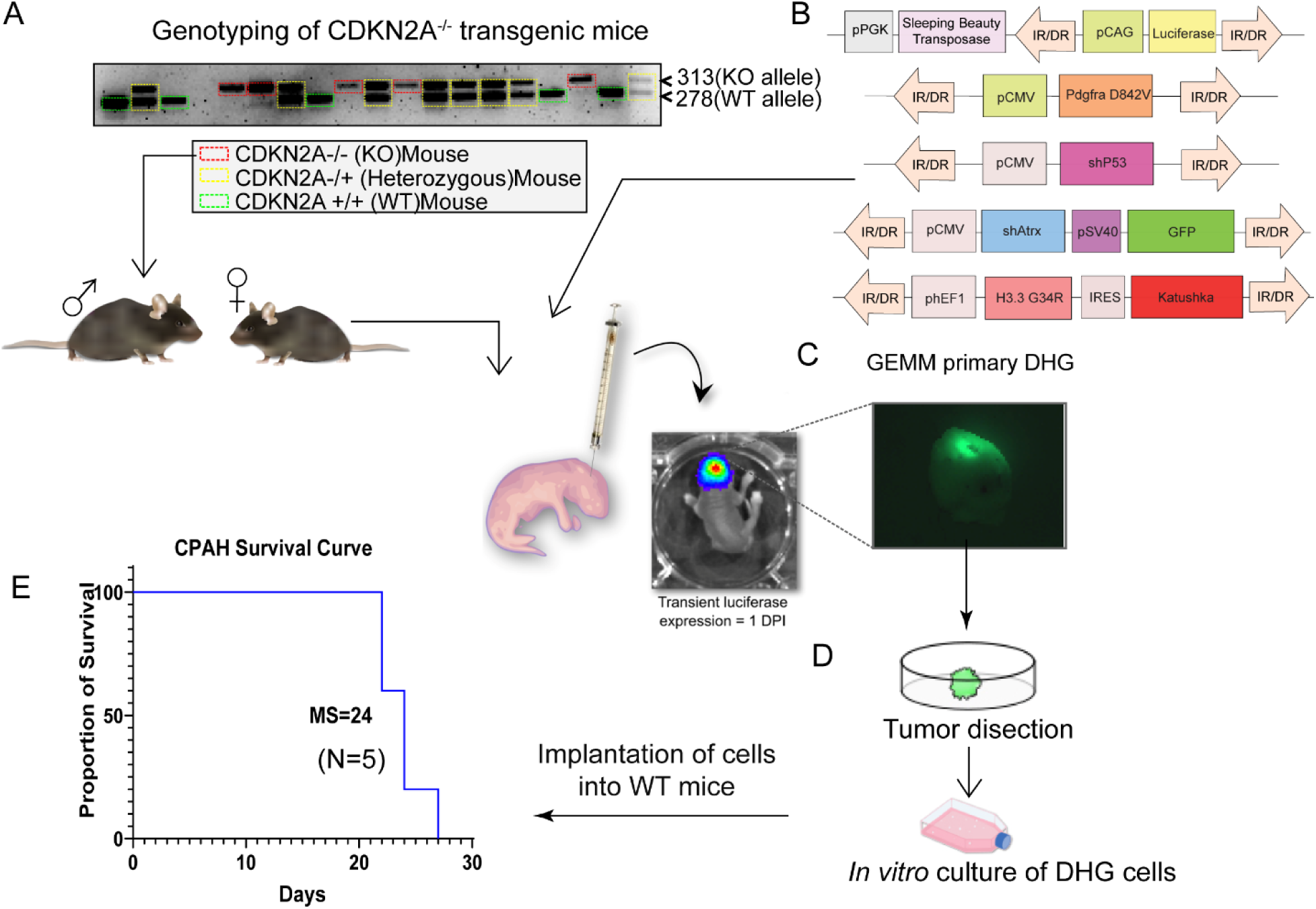
Development and characterization of a CDKN2A Knock out (KO), P53 Knock down (KD) and Atrx Knock down (KD) DHG model, related to Figure 4. **(A)** The CDKN2A knock out background was provided by the genetic phenotype of the host mice. CDKN2A heterozygous (+/-) knock out embryos were implanted in a receptive female host and the first generation was bred. The progeny was genotyped by PCR as depicted in the picture, where homozygous KO animals show a single 313 bp product (indicated in red). CDKN2A homozygous knock out (-/-) mice were mated, and the progeny (3rd generation) were implanted in the subventricular area of the brains at day 1 post birth with a mixture of plasmids **(B)** composed of a sleeping beauty transposase expressing plasmid, and plasmids expressing interfering short hairpin RNAs against P53 and ATRX. The development of gliomas was monitored by bioluminescence, and when the animals became symptomatic, they were euthanized for tumor dissection. The green fluorescence incorporated in the ATRX KD plasmid was used to identify tumor tissue **(C)**. The tumor tissue dissected was detached and primary tumor cells were grown in vitro **(D)**. The 2,000 glioma cells cultured in vitro were reimplanted in WT mice and developed gliomas with a median survival of 22 days **(E)**.

**Figure S13.**
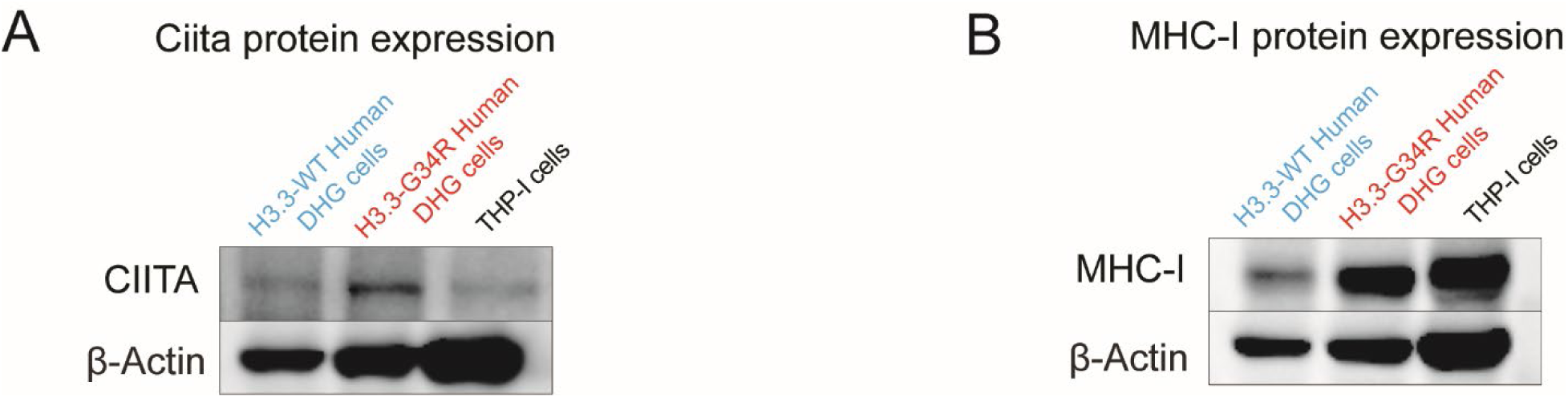
Expression of MHC-I in human DHG, related to Figure 4. **(A)** Western blot for CIITA at basal levels and upon treatment with IFN-γ in H3.3-WT and H3.3-G34R human DHG cells. **(B)** Western blot for MHC-I at basal levels and upon treatment with IFN-γ in H3.3-WT and H3.3-G34R human DHG cells. THP-1 cells were included as positive control.

**Figure S14.**
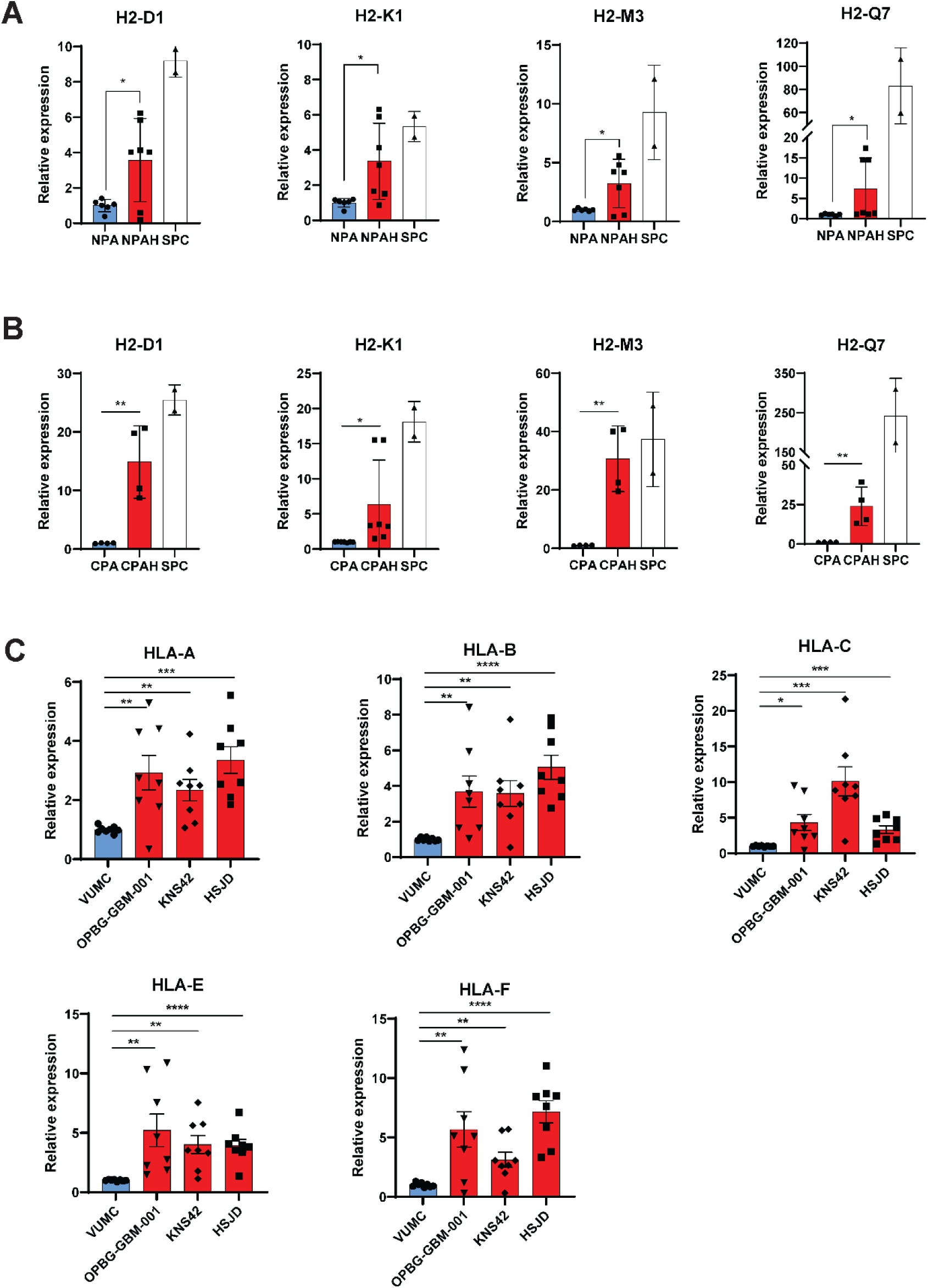
Expression of MHC-I genes, related to Figure 4. **(A)** Analysis of the expression of MHC-I genes in mouse H3.3-G34R versus H3.3-WT cells by real-time PCR. Mouse splenocytes were used as MHC-I-expressing control cells. **(B)** Analysis of the expression of MHC-I genes in mouse H3.3-G34R versus H3.3-WT cells by real-time PCR. Mouse splenocytes were used as MHC-I-expressing control cells. **(C)** Analysis of the expression of MHC-I genes in patient derived H3.3-G34R versus H3.3-WT cells by real-time PCR. *P < 0.05, unpaired t test.

**Figure S15.**
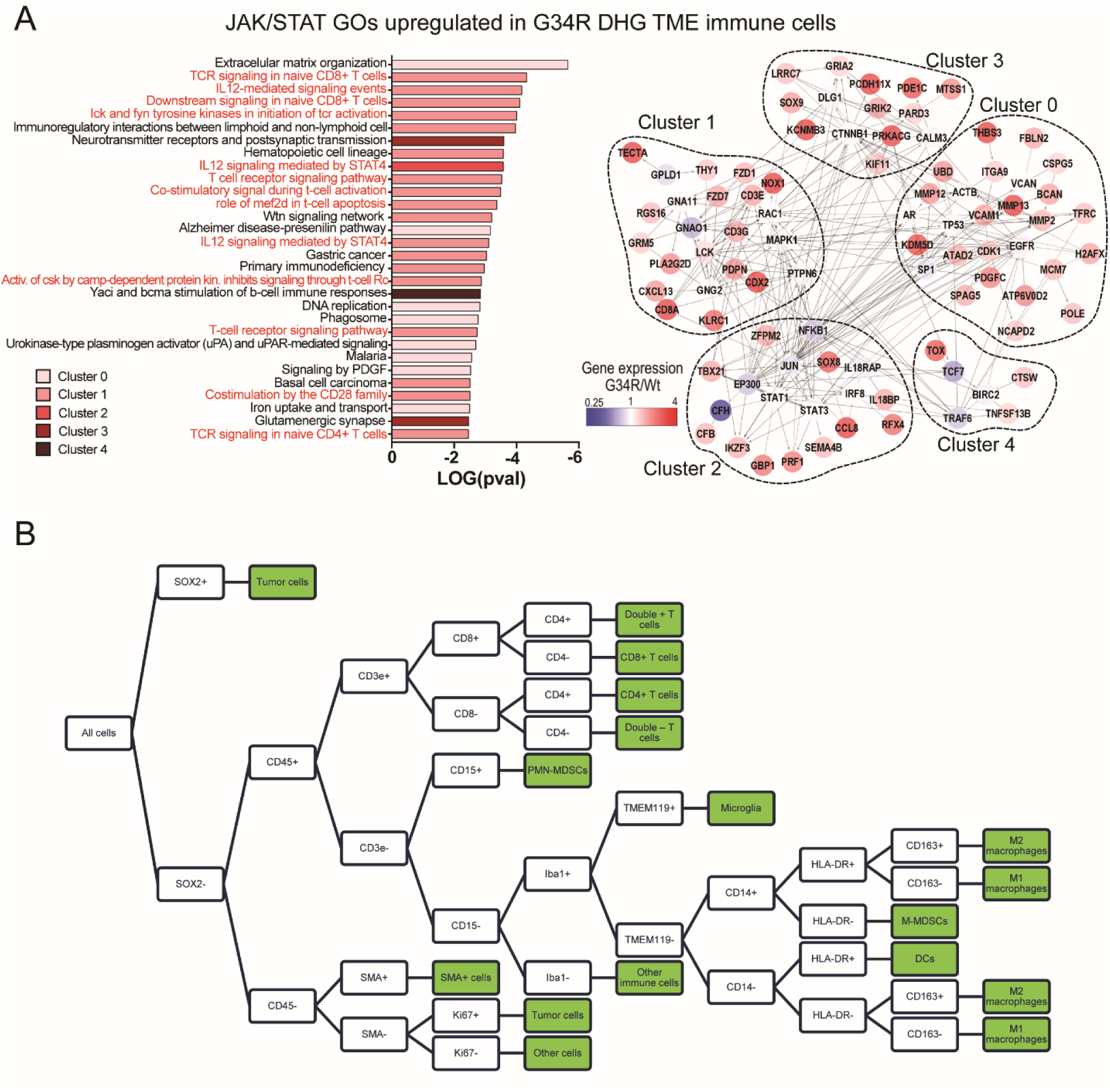
Analysis of H3.3-G34R DHG TME immune cells, related to Figure 5. **(A)** Plot showing the 30 most significant enriched pathways (x-axis) in terms of p-value (-Log(p-value), y-axis). Pathways related to T-cell activation and function are highlighted in red; and network analysis of differentially upregulated genes in H3.3-G34R CD45+ cells with respect to H3.3-WT CD45+ cells. Clusters of nodes of identical color represent a cluster of genes involved in similar biological pathways. Linker genes have been used to construct the network and they are depicted in red. Numbers indicate the cluster number. *** p<0.001; * p<0.05; Student T test. **(B)** Cell classification was done hierarchically, as shown, with a script implementing object classifiers for each marker that were trained using artificial neural networks.

**Figure S16.**
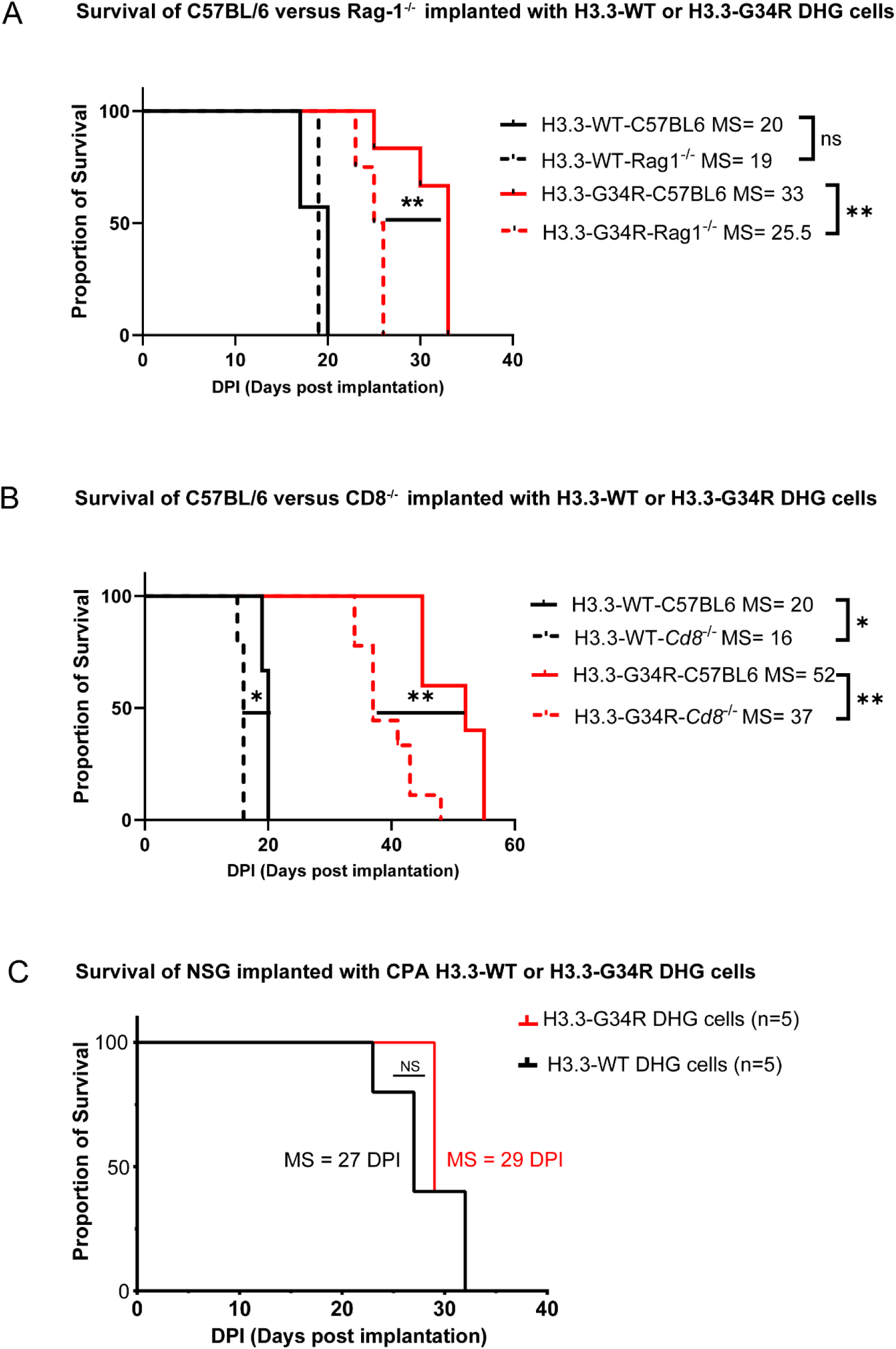
Lymphocytes impact H3.3-G34R tumor progression, related to Figure 6. Fifty thousand H3.3-WT or H3.3-G34R neurospheres were implanted in C57BL6, Rag-1-/- **(A)** or Cd8-/- **(B)** mice (n=5) and, at symptomatic stage, animals were euthanized and survival as days-post implantation (DPI) was documented. Kaplan-Meier survival curves comparing the median survival (MS) of H3.3-WT and H3.3-G34R, implanted in C57BL6 and A) Rag-1-/- or B) Cd8-/- mice are shown. **(C)** Survival of NSG mice implanted with CDKN2A-/-, P53 KD, ATRX KD H3.3-WT or H3.3-G34R DHG cells. *p<0.05; **p<0.01; ***p<0.001; ns=non-significant; Mantel-Cox test.

**Figure S17.**
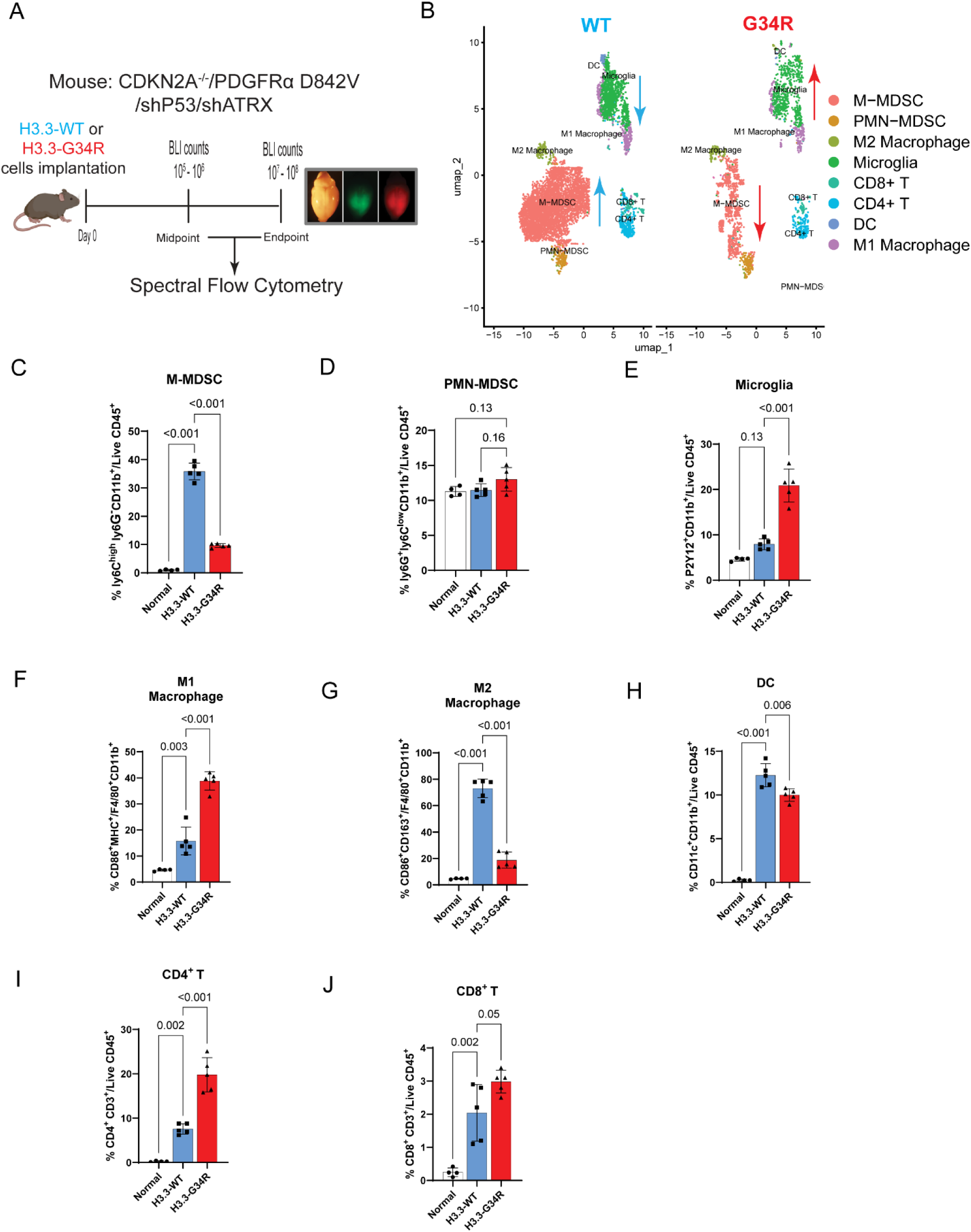
Spectral flow cytometry Analysis of the tumor immune microenvironment (TIME) of CDKN2A^-/-^, PDGFRa D842V H3.3-G34R DHG at <u>end</u> <u>stage</u>, related to Figure 6. **(A)** Illustration of experimental procedure followed for the characterization of immune cells in H3.3-G34R mouse DHG TIME. **(B)** UMAP embedding of H3.3 G34R/V and WT DHG immune cell SFC analysis. **(C-D)** The proportion of cells with Monocytic (CD45+/CD11b+/Ly6G-Ly6Chi) or Polymorphonuclear (CD45+/CD11b+/Ly6G+Ly6Clo) myeloid derived suppressor cell phenotype was performed on H3.3-WT and H3.3-G34R TIMEs. **(E)** The total proportion of microglia is shown as the percentage of CD45 low/CD11b+/TMEM119+ P2Y12+ cells. **(F-G)** Macrophage were further classified as M1 or M2 phenotype (CD45+/Ly6C-F4/80+/CD86+MHC+ or CD45+/Ly6C-F4/80+/CD163+CD206+, respectively). **(H)** The total proportion of DC is shown as the percentage of CD45+/CD11b+/CD11c+ **(I-J)** Total pan T cells were identified as CD45+/CD3+ cells and further classified by the expression of CD8 or CD4 (CD45+/CD3+/CD8+ or CD45+/CD3+/CD4+, respectively). Data are represented as mean±SD. Statistical differences determined with one-way ANOVA Tukey’s post hoc test. ns, not significant.

**Figure S18.**
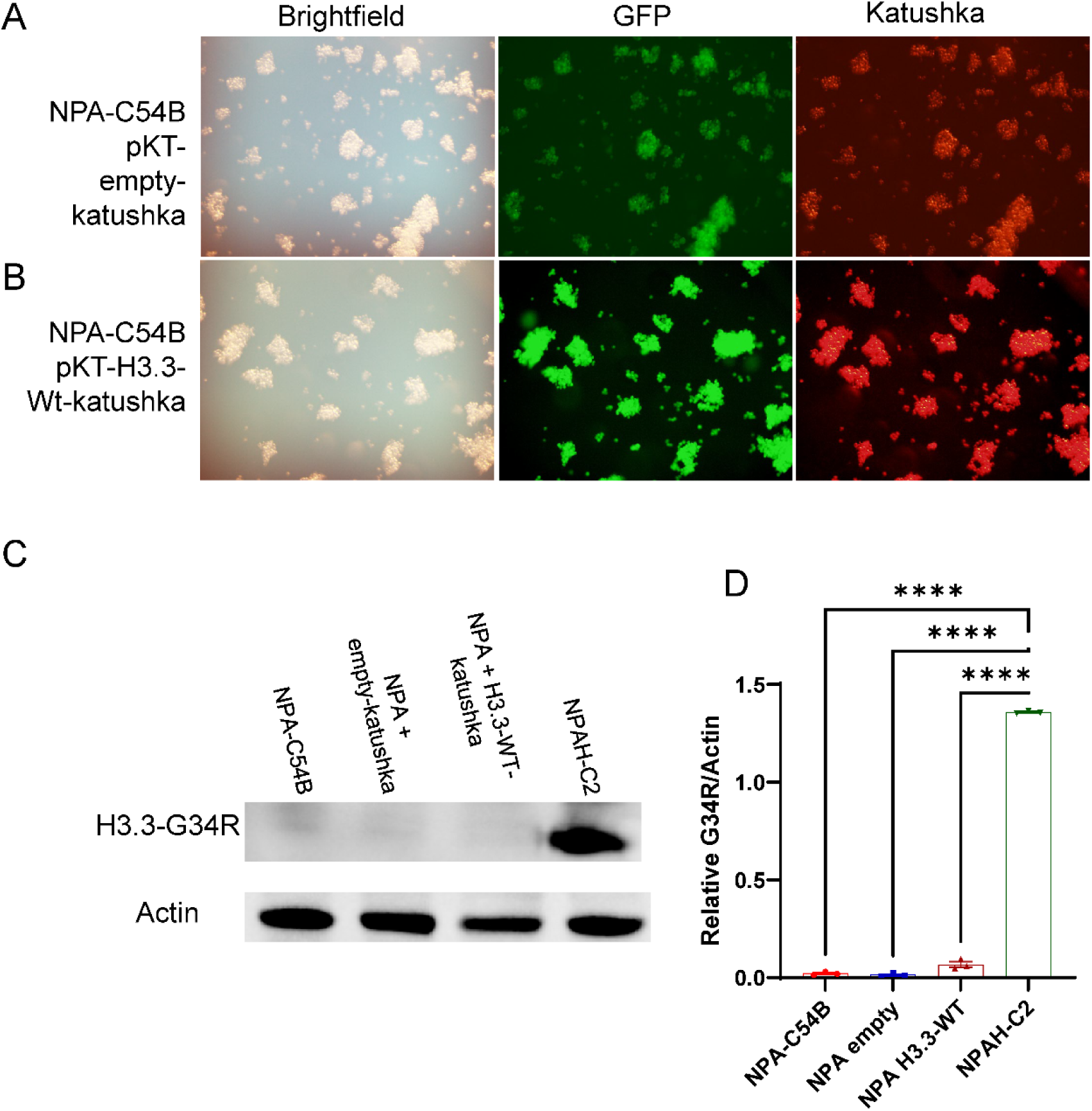
Generation of a mouse DHG model (NPA: NRASG12V, TP53 KD, and ATRX KD) expressing fluorescent marker Katushka, related to Figure 6. Fluorescent microscope images of **(A)** NPA-C54B transfected with pKT-empty-Katushka and **(B)** NPA-C54B transfected with pKT-H3.3-WT-Katushka. Images were taken at 10X magnification. **(C)** Western blot of transfected NPA cells to show expression of H3.3-G34R. **(D)** Densiometric analysis of the Western blot. ****p<0.0001. KD: knockdown.

**Figure S19.**
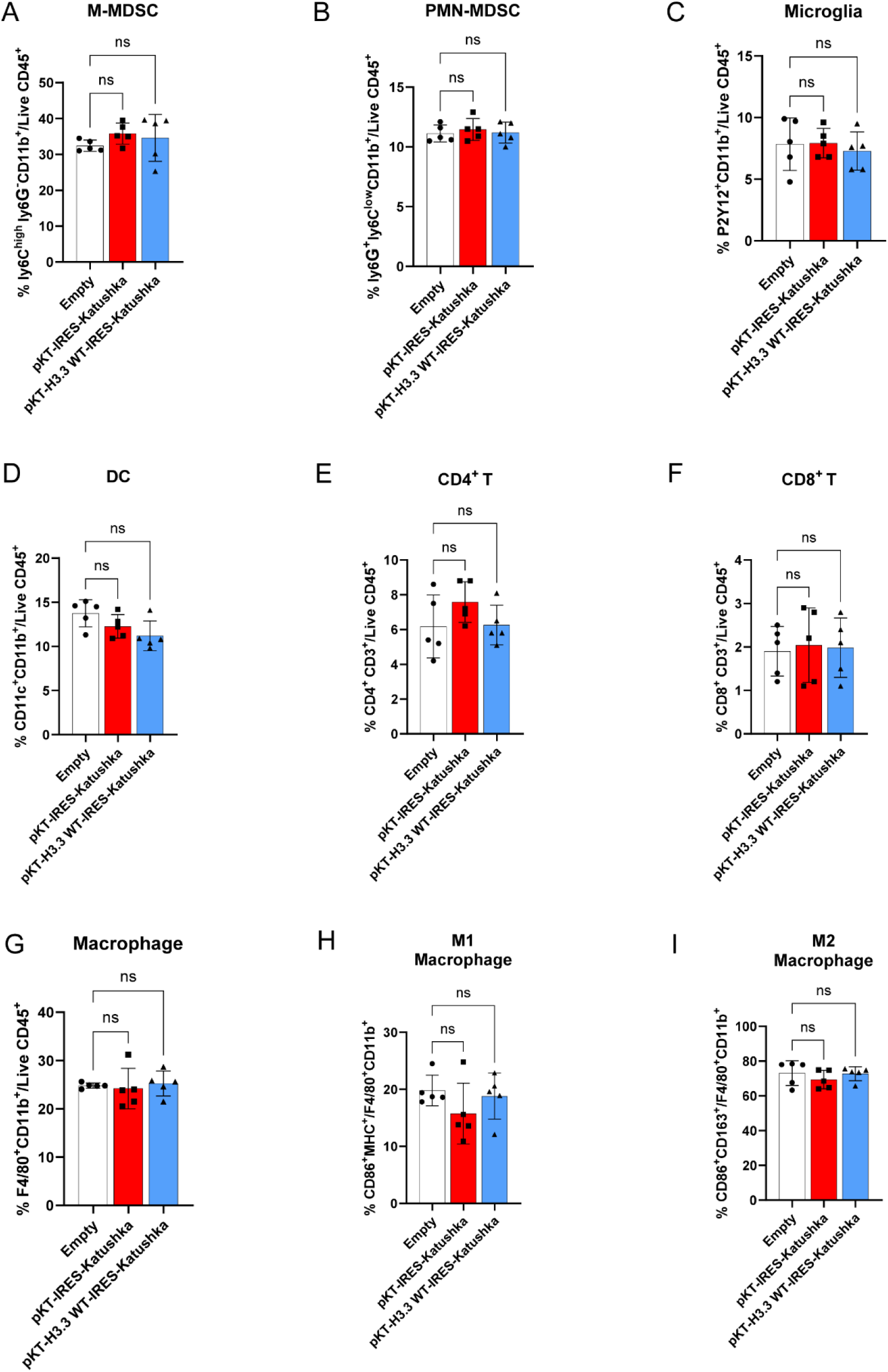
The impact of Katushka expression in the tumor immune microenvironment (TIME) of DHG tumors using spectral flow cytometry analysis, related to Figure 6. **(A-B)** The proportion of cells with Monocytic (CD45+/CD11b+/Ly6G-Ly6Chi) or Polymorphonuclear (CD45+/CD11b+/Ly6G+Ly6Clo) myeloid derived suppressor cells. **(C)** The total proportion of microglia is shown as the percentage of CD45 low/CD11b+/TMEM119+ P2Y12+ cells. **(D)** The total proportion of DC is shown as the percentage of CD45+/CD11b+/CD11c+ **(E-F)** Total pan T cells were identified as CD45+/CD3+ cells and further classified by the expression of CD8 or CD4 (CD45+/CD3+/CD8+ or CD45+/CD3+/CD4+, respectively). **(G)** The total proportion of macrophages is shown as the percentage of CD45+/CD11b+/F4/80+ cells. **(H-I)** Macrophage were were further classified as M1 or M2 phenotype (CD45+/Ly6C-F4/80+/CD86+MHC+ or CD45+/Ly6C-F4/80+/CD163+CD206+, respectively). Data are represented as mean±SD. Statistical differences determined with one-way ANOVA Tukey’s post hoc test. ns, not significant.

**Figure S20.**
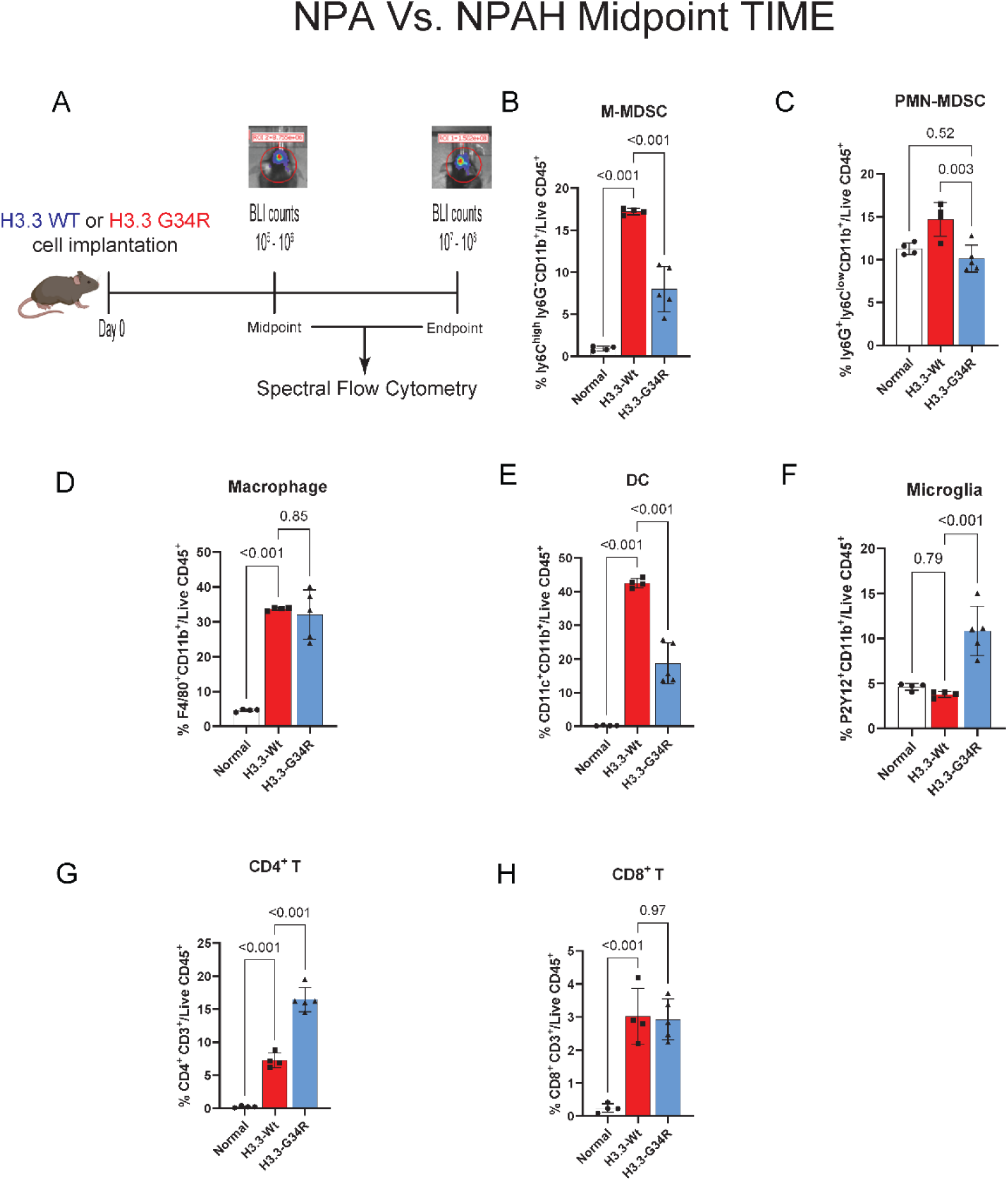
Spectral flow cytometry Analysis of the tumor immune microenvironment (TIME) of DHG <u>midpoint</u> tumors, related to Figure 6. **(A)** Illustration of experimental procedure followed for the characterization of immune cells in H3.3-G34R mouse DHG TIME. **(B-C)** The proportion of cells with Monocytic (CD45+/CD11b+/Ly6G-Ly6Chi) or Polymorphonuclear (CD45+/CD11b+/Ly6G+Ly6Clo) myeloid derived suppressor cells. **(D)** The total proportion of macrophages is shown as the percentage of CD45+/CD11b+/F4/80+ cells. **(E)** The total proportion of microglia is shown as the percentage of CD45 low/CD11b+/TMEM119+ P2Y12+ cells. **(F)** The total proportion of DC is shown as the percentage of CD45+/CD11b+/CD11c+ **(G-H)** Total pan T cells were identified as CD45+/CD3+ cells and further classified by the expression of CD8 or CD4 (CD45+/CD3+/CD8+ or CD45+/CD3+/CD4+, respectively). Data are represented as mean±SD. Statistical differences determined with one-way ANOVA Tukey’s post hoc test. ns, not significant.

**Figure S21.**
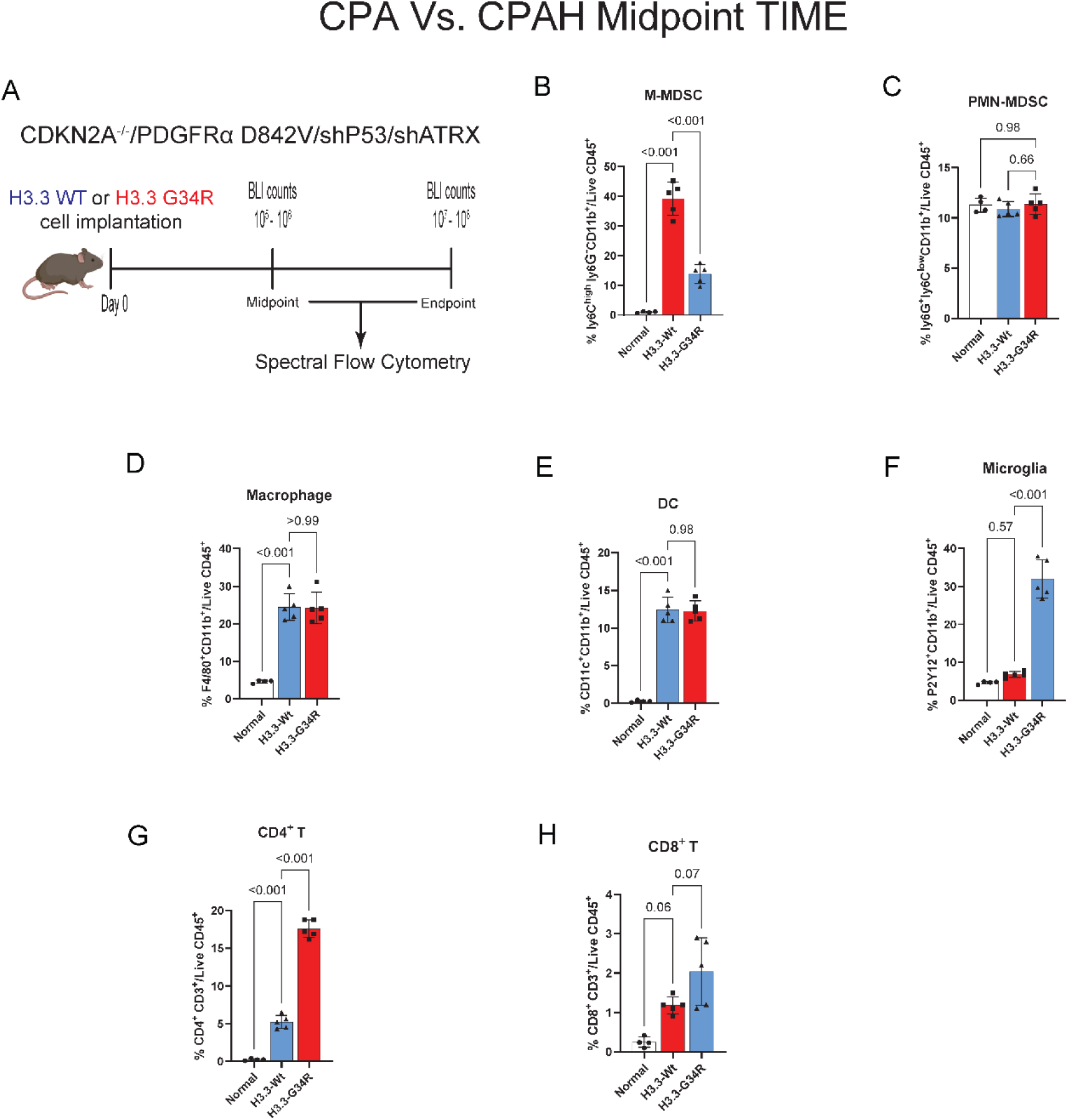
Spectral flow cytometry Analysis of the tumor immune microenvironment (TIME) of CDKN2A^-/-^, PDGFRa D842V driven DHG <u>midpoint</u> tumors, related to Figure 6. **(A)** Illustration of experimental procedure followed for the characterization of immune cells in H3.3-G34R mouse DHG TIME. **(B-C)** The proportion of cells with Monocytic (CD45+/CD11b+/Ly6G-Ly6Chi) or Polymorphonuclear (CD45+/CD11b+/Ly6G+Ly6Clo) myeloid derived suppressor cells. **(D)** The total proportion of macrophages is shown as the percentage of CD45+/CD11b+/F4/80+ cells. **(E)** The total proportion of microglia is shown as the percentage of CD45 low/CD11b+/TMEM119+ P2Y12+ cells. **(F)** The total proportion of DC is shown as the percentage of CD45+/CD11b+/CD11c+ **(G-H)** Total pan T cells were identified as CD45+/CD3+ cells and further classified by the expression of CD8 or CD4 (CD45+/CD3+/CD8+ or CD45+/CD3+/CD4+, respectively). Data are represented as mean±SD. Statistical differences determined with one-way ANOVA Tukey’s post hoc test. ns, not significant.

**Figure S22.**
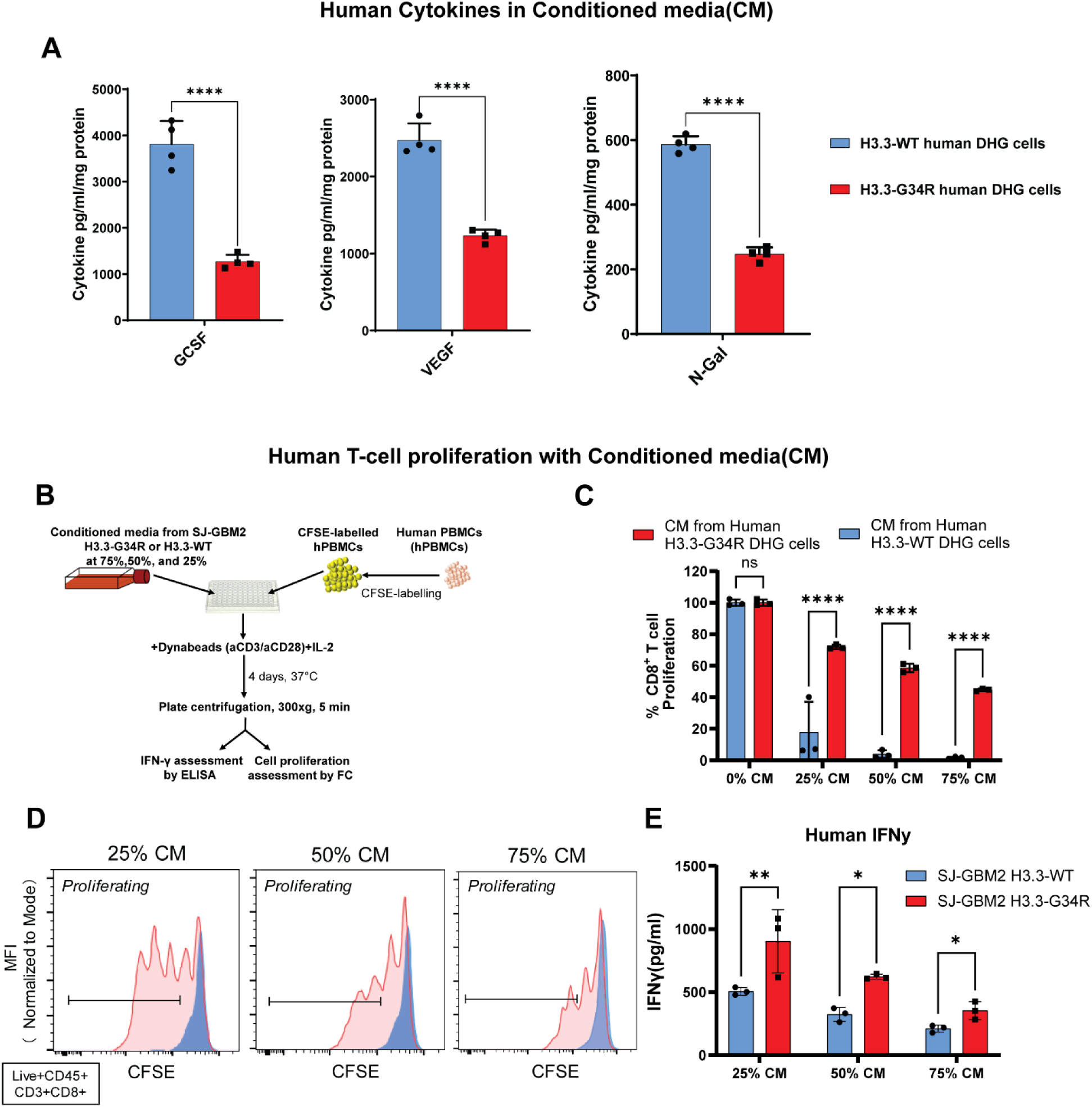
Cytokines profile of human G34R DHG cells and immune properties of the conditioned media, related to Figure 7. **(A)** To assess the concentration of different cytokines, human H3.3-WT and H3.3-G34R cells conditioned media was analyzed by ELISA. Total protein present in each flask was determined and cytokine concentration was normalized to protein quantity. ****p<0.0001, 2-way ANOVA and Sidak’s multiple comparisons test. **(B)** Experimental layout of the T cell proliferation assay to test the immunosuppressive capacity of conditioned media (CM) from Human H3.3-G34R or H3.3-WT DHG cells. **(C)** The percentage of proliferating CD8+ T cells in the presence of CM from Human H3.3-WT or H3.3-G34R DHG cells, with respect to the positive control (0% CM). **(D)** Representative histograms show the CD8+ T cell proliferation in the presence of 25%, 50% or 75% CM from Human H3.3-WT or H3.3-G34R DHGs. **(E)** IFNγ concentration was measured in the supernatant of the hPBMC cultures incubated with the H3.3-WT and H3.3-G34R conditioned media by ELISA. Bars represent cytokine concentration in absolute numbers.

**Figure S23.**
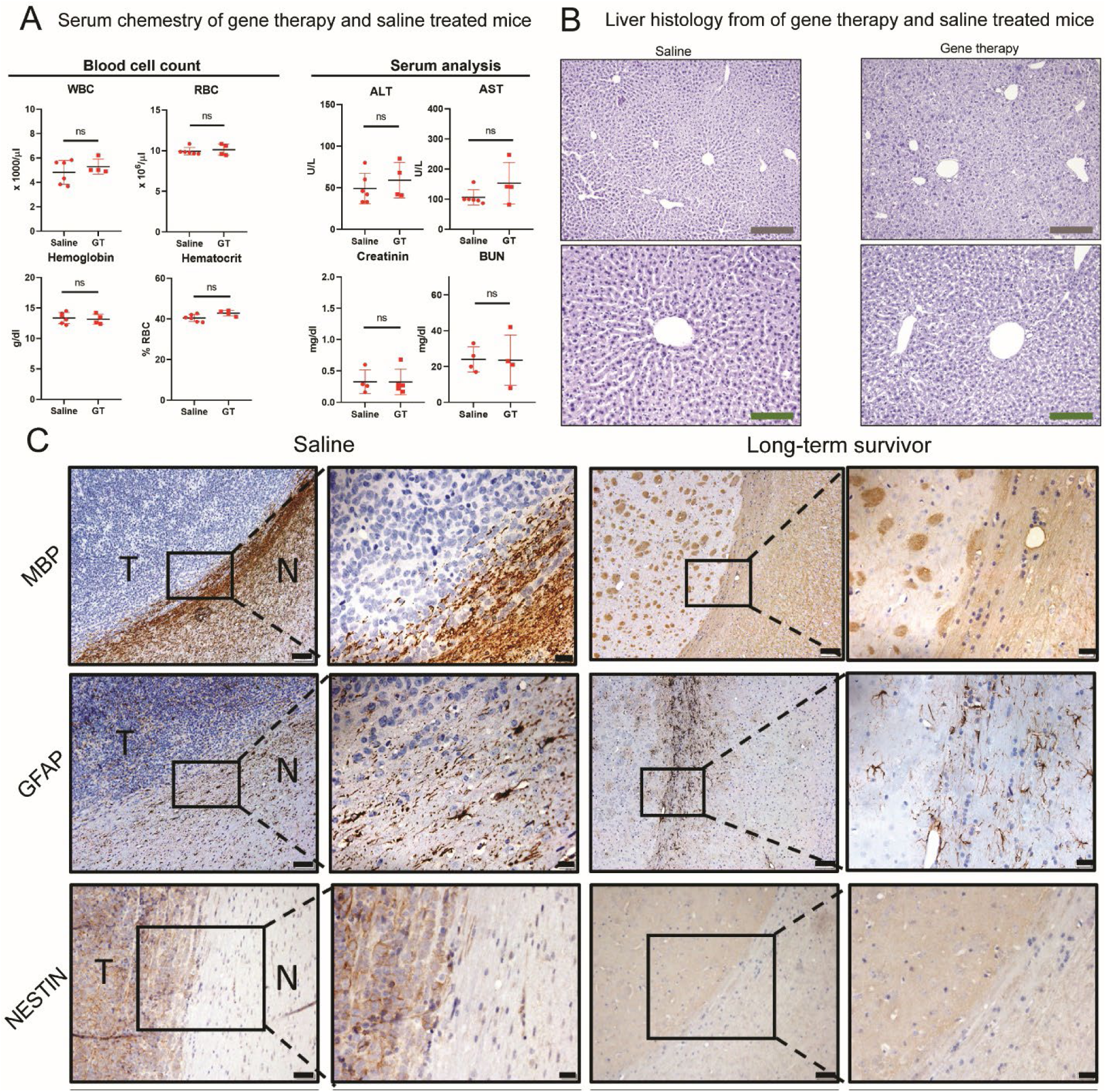
Safety assessment of gene therapy (GT) administration., related to Figure 8. **(A)** Blood cells count and serum analysis of H3.3-G34R tumor bearing mice treated with Saline or GT. Animals harboring H3.3-G34R brain tumors were treated with GT or Saline, and, at symptomatic stage, animals were euthanized, and blood was extracted to perform complete cell blood count and serum chemistry analysis. All parameters were compared between saline and GT groups and their differences were non-significant. ns= non-significant, Student t test. **(B)** Histopathology of livers from H3.3-G34R tumor bearing mice treated with Saline or GT. Animals that succumbed due to tumor burden after Saline or GT treatment was perfused and processed for hematoxylin and eosin (H&E) staining. Grey scale bar: 200 µm. Green scale bar: 100 µm. **(C)** Long term survivors from H3.3-G34R GT treatment were implanted with 50,000 H3.3-G34R neurospheres in the contralateral hemisphere and 70 days post rechallenge animals were euthanized and processed for histological analysis. Paraffin embedded 5 µm E) brain and F) liver sections were stained for hematoxylin and eosin (H&E). Black scale bar: 1 mm. Grey scale bar: 200 µm. Green scale bar: 100 µm.

**Figure S24.**
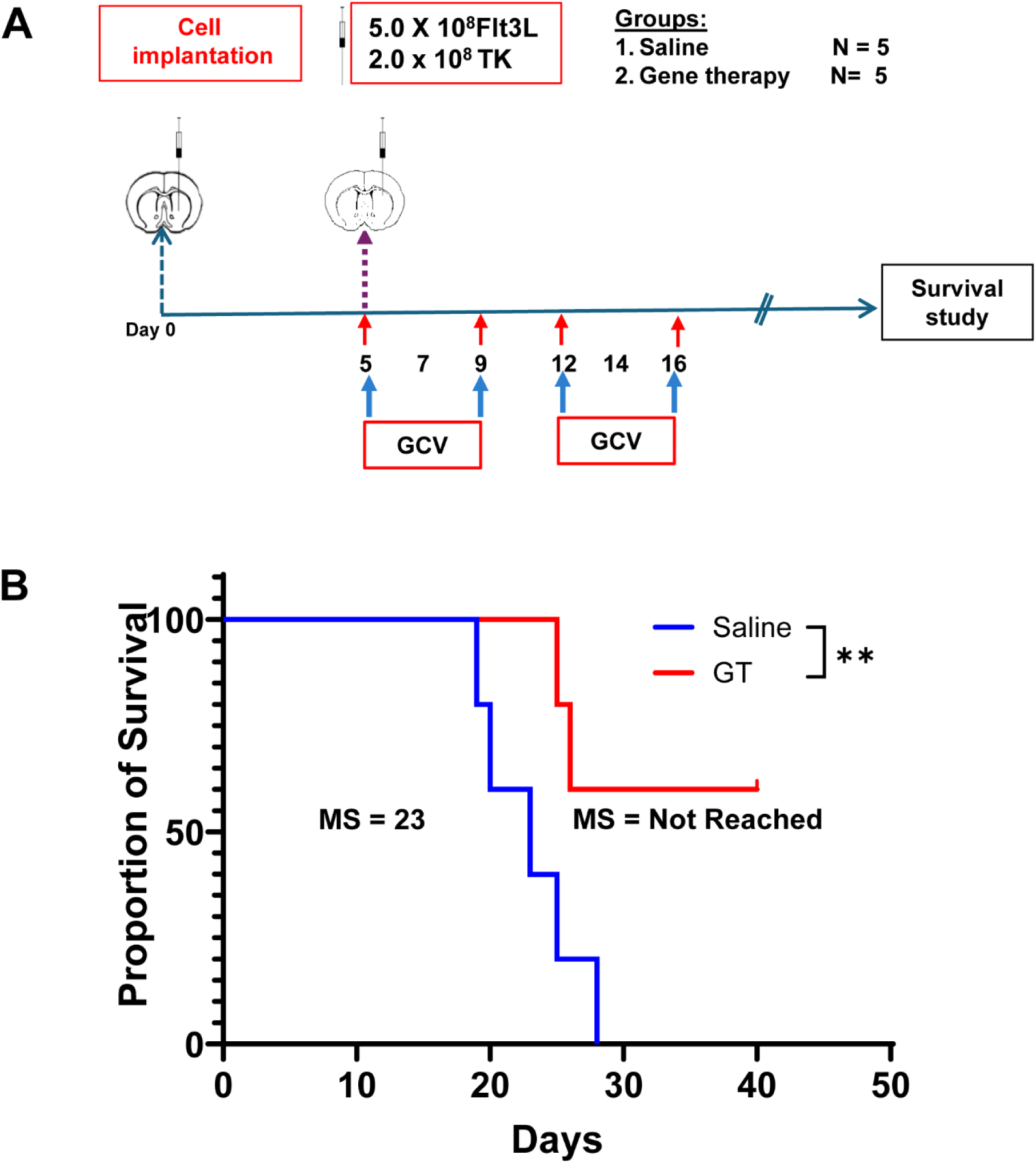
Therapeutic efficacy of gene therapy in CDKN2A^-/-^, PDGFRa D842V H3.3-G34R DHGs, related to Figure 8. **(A)** Schematic representation of immune mediated gene therapy regimen administered to mice. **(B)** Kaplan-Meier survival curve showing the survival of mice treated with saline and gene therapy. Median survival was (MS 23 days) for the saline-treated group and MS not reached for the gene therapy-treated group. Statistical significance was assessed using the Log-rank (Mantel-Cox) test (p=0.0256).

**Figure S25.**
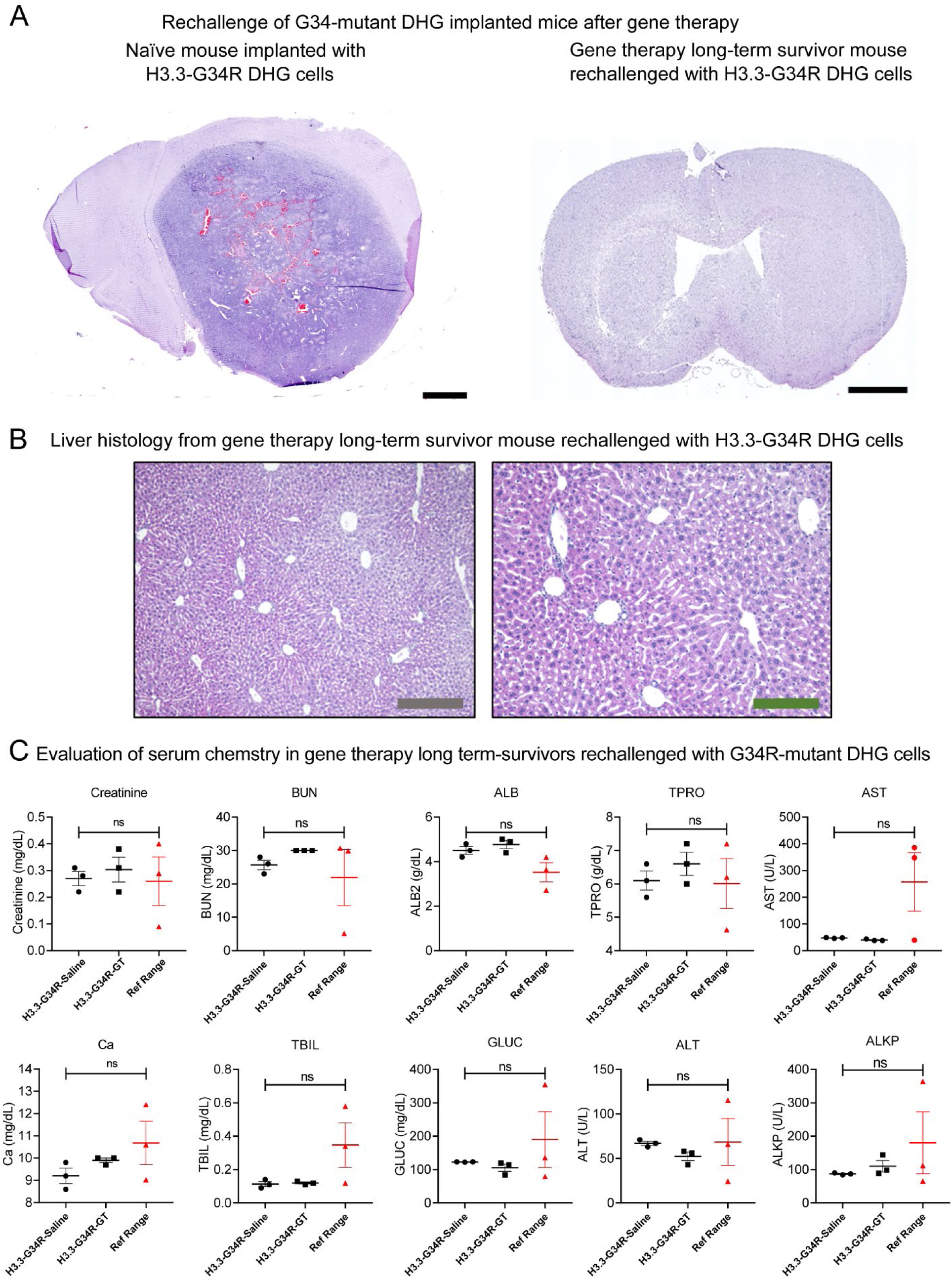
Rechallenge of gene therapy long term survivors, related to Figure 8. **(A)** Brain hematoxylin/Eosin histology from Naïve mouse implanted with H3.3-G34R DHG cells versus gene therapy long-term survivor rechallenged with H3.3-G34R DHG cells. Gene therapy long term survivor does not show signs of tumor development due to development of immunological antitumoral memory. **(B)** Long term survivors from H3.3-G34R GT treatment were implanted with 50,000 H3.3-G34R neurospheres in the contralateral hemisphere and 70 days post rechallenge animals were euthanized and processed for histological analysis. Paraffin embedded 5 µm liver sections were stained for hematoxylin and eosin (H&E). Black scale bar: 1 mm. Grey scale bar: 200 µm. Green scale bar: 100 µm. **(C)** Assessment of serum metabolites in gene therapy long term survivors rechallenged with G34R-mutant DHG cells. BUN= Blood Urea Nitrogen, ALT = alanine transaminase, AST = Aspartate Transferase (AST), ALKP = alkaline phosphatase (ALP), GLUC = glucose, TPRO = total protein, ALB2 = albumin, TBIL = Total Bilirubin, Ca = Calcium. Levels are compared to a reference range (Ref. Range).

## Notes

### Competing Interest Statement

The authors have declared no competing interest.

### Summary of Updates

Please note the usage of diffuse hemispheric glioma (DHG) following the 2021 WHO classification of tumors of the central nervous system.

